# Identification of Determinants that Allow Maintenance of High-Level Fluoroquinolone Resistance in *Acinetobacter baumannii*

**DOI:** 10.1101/2023.10.03.560562

**Authors:** Efrat Hamami, Wenwen Huo, Juan Hernandez-Bird, Arnold Castaneda, Jinna Bai, Sapna Syal, Juan C. Ortiz-Marquez, Tim van Opijnen, Edward Geisinger, Ralph R. Isberg

## Abstract

*Acinetobacter baumannii* is associated with multidrug resistant (MDR) infections in healthcare settings, with fluoroquinolones such as ciprofloxacin being currently ineffective. Clinical isolates largely harbor mutations in the GyrA and TopoIV fluoroquinolone targets, as well as mutations that increase expression of drug resistance-nodulation-division (RND) efflux pumps. Factors critical for maintaining fitness levels of pump overproducers are uncharacterized despite their prevalence in clinical isolates. We here identify proteins that contribute to the fitness of FQR strains overexpressing three known RND systems using high-density insertion mutagenesis. Overexpression of the AdeFGH efflux pump caused hypersensitization to defects in outer membrane homeostatic regulation, including lesions that reduced LOS biosynthesis and blocked production of the major *A. baumannii* porin. In contrast, AdeAB pump hyperexpression, in the absence of elevated *adeC* expression (the outer membrane component of the pump), was relatively tolerant to loss of these functions, consistent with the outer membrane protein being the primary disruptive component. Surprisingly, overexpression of proton-transporting efflux pumps had little impact on cytosolic pH, consistent with a compensatory response to pump activity. The most striking transcriptional changes were associated with AdeFGH pump overexpression, including the activation of the phenylacetate (PAA) degradation regulon. Disruption of the PAA pathway resulted in cytosolic acidification and defective expression of genes involved in protection from oxidative stress. These results indicate that RND efflux pump overproduction is compensated by maintenance of outer membrane integrity in *A. baumannii* to facilitate fitness of FQR isolates.

**Importance:** *Acinetobacter baumannii* is a pathogen that often causes multidrug resistant (MDR) infections in healthcare settings, presenting a threat to the efficacy of known therapeutic interventions. Fluoroquinolones such as ciprofloxacin are currently ineffective against a majority of clinical *A. baumannii* isolates, many of which express pumps that remove this antibiotic class from within the bacterium. Three of these pumps can be found in most clinical isolates, with one of the three often hyperproduced at all times. In this study we identify proteins that are necessary for the fitness of pump hyperproducers. The identified proteins are necessary to stabilize the outer membrane and allow the cytoplasm to tolerate the accumulation of ions as a consequence of excess pump activity. These results point to strategies for developing therapies that combine known antibiotics with drugs that target proteins important for survival of strains hyperexpressing efflux pumps.

## Introduction

*Acinetobacter baumannii* is a Gram-negative bacterium primarily associated with diseases in healthcare settings (1–3). The microorganism has the potential to cause a diverse range of diseases including pneumonia, wound, and urinary tract infections (1–3). Of great concern is its role in causing ventilator-associated pneumonia, which may be closely tied to its ability to grow in biofilms on indwelling devices (2–6). Particularly challenging is the high incidence of multidrug resistant (MDR) *A. baumannii* in hospitalized patients, leaving afflicted individuals with few treatment options (1–3). Although there was a reduction in nosocomial cases in the US from 2013 to 2019, the SARS-CoV2 pandemic saw a large jump in cases that illustrates the difficulties in suppressing diseases by this organism (3). Furthermore, the increasingly common resistance profiles of clinical isolates that are refractory to most antimicrobial therapy regimens indicate that the pathogen poses a continued threat to successful treatment (1–3, 7–10). These clinical isolates include a high proportion that are multi-drug resistant (MDR) or have decreased susceptibility to various single agents (1–3, 7–10). This has prompted the World Health Organization to report that *A. baumannii* is a high priority candidate for research and development of new antimicrobials (8). For this reason, there is a high demand for novel antimicrobials or improved combination therapies with which we can combat this pathogen.

There are a variety of means by which *A. baumannii* acquires resistance, including the acquisition of drug resistance elements via horizontal gene transfer (HGT), the selection of mutations that inactivate drug target sites, and hyper-expression of preexisting drug efflux pumps (11–15). Pathogens not only harbor a multitude of efflux systems, some of which are acquired by HGT, but many have broad substrate specificities (11–15), stimulating efforts to discover effective pump inhibitors (16). Unfortunately, there has been no indication of progress in bringing these inhibitors to the clinic, although several have proven effective in interfering with drug action or synergizing with clinically useful antibiotics using laboratory culture conditions or in animal models (16). Nevertheless, given the relentless patterns of resistance acquisition after the use (and misuse) of single-target antibiotics, a treatment regimen that uses a combination of readily available antimicrobials working in synergy could be effective in interfering with resistance evolution. This study approaches this concept by determining candidate proteins that could be targeted together with clinically-proven classes of antibiotics.

One drug class that has reduced effectiveness against nosocomial pathogens is the fluoroquinolones (2, 17). Resistance in clinical isolates primarily selects for mutations that disrupt drug binding to the targets topoisomerases II and IV, encoded for by *gyrA* and *parC* genes respectively (17). In *A. baumannii*, the highest levels of fluoroquinolone resistance (FQR) are found in isolates that have increased expression of drug efflux pumps in addition to mutations in the drug targets. These drug efflux pumps belong to the resistance-nodulation-division (RND) family, in which overexpression is commonly driven by mutations in regulatory elements that result in constitutive expression (18–23). The three major efflux pumps commonly associated with FQR in *A. baumannii* clinical isolates are encoded for by *adeAB*, *adeFGH*, and *adeIJK*, which are controlled by *adeRS* (encoding a two-component system), *adeL* (a LysR type regulator), and *adeN* (a TetR repressor) regulators respectively (18–23). Among these three, the AdeN-regulated pump is associated with intrinsic resistance to fluoroquinolones, as this is the only RND pump in *A. baumannii* that has measurable levels of expression in the absence of activating mutations (17, 22, 23).

These pump systems traverse both membranes (24–30), and overproduction requires that the envelope accommodates an elevated population of protein complexes not present in drug-sensitive isolates, putting potential stress on the inner and outer membrane. Therefore, the stability and assembly of these proteins, as well as the fitness of *A. baumannii* during exponential growth may rely upon bacterial functions that work to maintain cell envelope integrity or prevent intrinsic toxicity of the overexpressed efflux system. Our work focuses on identifying genetic interactions in these high-level FQR strains that allow them to maximize fitness, with the goal of identifying proteins necessary to buffer the deleterious effects of pump hyperexpression that could lead to decreased fitness levels in these strains. Such mutations are predicted to identify targets for drugs that are specifically toxic for pump overproducers.

In this study, we used dense transposon sequencing and transcriptomics strategies to identify proteins necessary to maintain fitness of *A. baumannii* efflux pump overexpressing strains. We demonstrate that strains harboring overexpressing mutations in *adeN* and *adeS* show surprising robustness and identify targets of vulnerability in overexpressers of the AdeL-regulated pump. These targets are central players in maintaining cell envelope homeostasis. We also connect the expression of the phenylacetate catabolic pathway as a response to pump hyperexpression that leads to maintaining cytosolic pH and maximizing protection from reactive oxygen species.

## Results

### Identification of mutations that depress fitness of fluoroquinolone-resistant strains overexpressing RND efflux pumps

To identify genes that are important for maintaining the fitness of strains having high-level fluoroquinolone resistance (FQR), we isolated mutations in the *A. baumannii* ATCC 17978UN strain background that are viable in high concentrations of ciprofloxacin (CIP; Materials and Methods). These strains were then subjected to a mutagenesis strategy (Fig. 1A) that identified insertion mutations that had lowered viability in an FQR strain background compared to the wild type (called strain WT) (Fig. 1B). Three FQR strains were interrogated, each of which harbored *gyrA*(S81L) and *parC*(S84L) mutations that are responsible for fluoroquinolone resistance in most Gram-negative organisms and are commonly found in the clinic (called strain GP). The three strains were distinguished from each other by having distinct mutations in loci that regulate the expression of clinically relevant *A. baumannii* resistance-nodulation-division (RND) drug efflux pumps. The mutation in a component of a two-component regulatory system *adeS*(R152S) upregulates the AdeAB pump, the activating *adeL*(ΔI335A336) mutation upregulates the AdeFGH pump, while a knockout in the repressor *adeN*::ISAba1 increases expression of the AdeIJK pump (Fig. 1C). It should be noted under conditions of AdeAB pump overexpression, AdeK is used as the outer membrane component which is expressed at wild type levels in the presence of the *adeS*(R152S) pump activating mutation (Fig. 1C) used in this study (31, 32).

**Figure 1.**
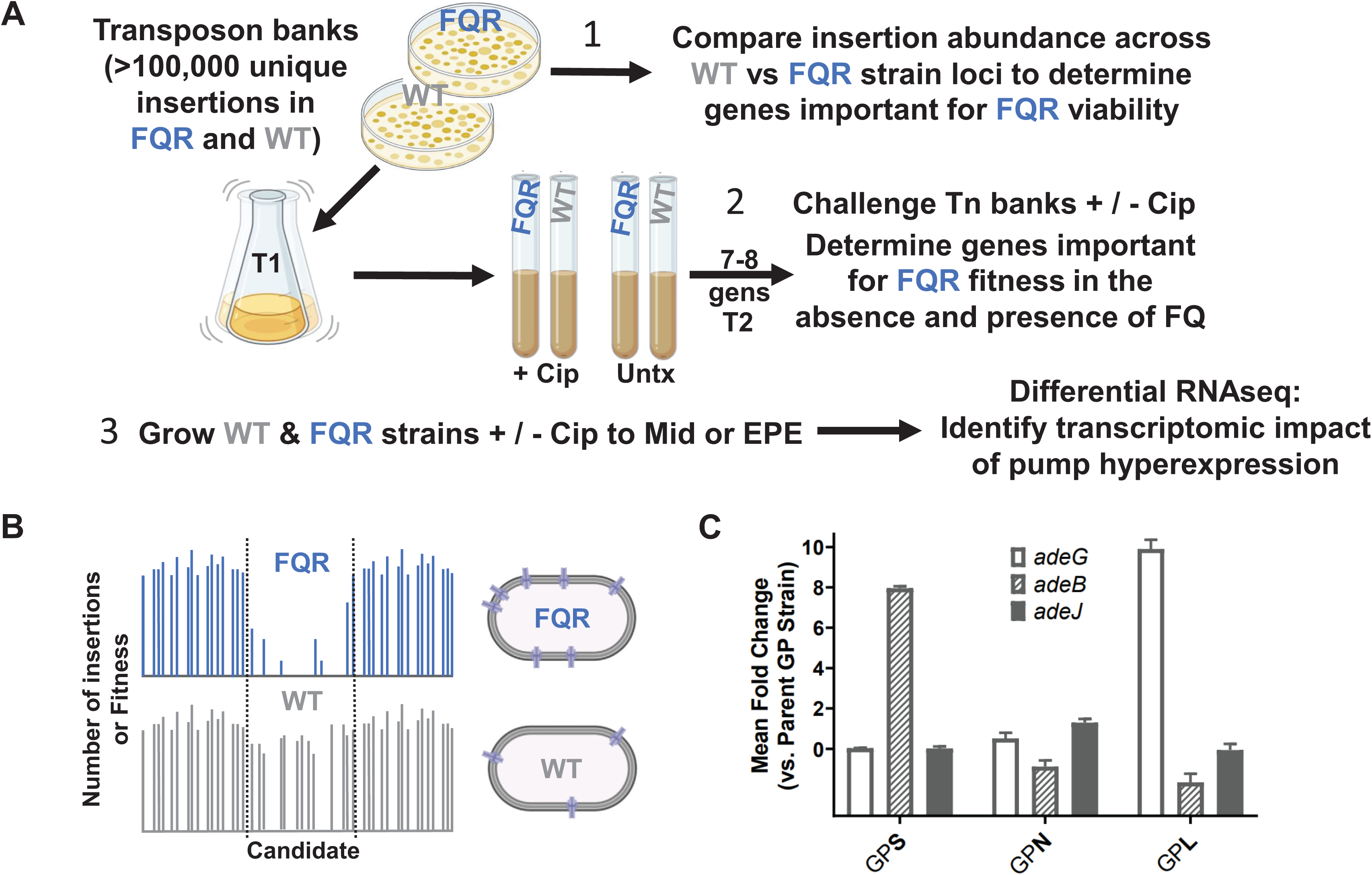
Uncovering the limitations of high-level FQR strains with enhanced RND pump expression. (A) Diagram for the three experimental approaches as described (Materials and Methods; created with BioRender.com). 1) Approach #1: Transposon insertion libraries were created in FQR strains overexpressing efflux pumps and the drug susceptible WT strain. Insertion abundance across the genome between the two strain types was compared to identify genes with fewer insertions in the FQR strain background than in WT (see B for example). 2) Approach #2: Transposon insertion banks were challenged with sub-MIC levels of CIP, parallel to those grown without drug. Fitness of all transposon mutants was calculated and compared between the strain types. 3) Approach #3: Strains were grown in presence or absence of CIP to exponential or post-exponential phase in presence or absence of drug and transcriptomic data was obtained to discover differentially regulated pathways. (B) Example of insertion abundance in a single gene cluster in two strain backgrounds. (C) Activating mutations in efflux pump regulatory mutants specifically activate their cognate pumps. RNA was isolated from the noted strains and relative transcription determined by q-rtPCR (Materials and Methods). Displayed is mean fold change ± SEM of each target as compared to the parental GP strain (Log2 scale). At least three independent biological replicates were used for each strain. FQR: fluoroquinolone resistant. Mid: mid exponential phase. EPE: early post-exponential phase. CIP: ciprofloxacin. Untx: untreated. WT: wildtype. GPL: *gyrAparCadeL*. GPN: *gyrAparCadeN*. GPS: *gyrAparCadeS*. GP: *gyrAparC*.

Transposon (Tn) mutagenesis was used to determine candidates important for viability or fitness in either the absence or presence of drug. Between 110,000-150,000 unique transposon mutants were created in all three FQR and the WT strain backgrounds, reaching pool saturation levels of about 37-50% based on targets predicted for the *himar-1* transposon (Materials and Methods). Each bank was sequenced post-harvest and transposon-proximal sequences were amplified to identify the locations of insertions in each strain background and to determine overall pool saturation. This sequence data was also utilized to identify insertion site distributions that were present in each strain, because insertions in genes predicted to be essential or cause fitness defects in a specific pump overproducer strain are predicted to be underrepresented relative to the WT strain. Per our first approach (Fig. 1A), insertion abundance throughout the genome was compared between strain backgrounds using the TRANSIT resampling method to identify genes that have fewer insertions relative to WT in the absence of drug (33) (Fig. 2, Data Set S1), as candidates for genes that are putative conditionally essential targets. To focus on strain-dependent conditional essentiality, 445 genes characterized as essential by TRANSIT’s Gumbel method (34, 35) were excluded from this and downstream analyses using transposon mutagenesis. Data were displayed by the statistical significance as a function of the insertion abundance in each of the pump overproducer strains relative to WT (Fig. 2A-C). Among genes identified in this fashion (passing significance criteria adj. p<0.05) in the absence of fluoroquinolones were *psiE* (*adeS* background), *sodB* (*adeN* background), and “nuc-diph-s-epim” or “n.d.s.e” (*adeL* background), respectively encoding a phosphate starvation inducible locus, superoxide dismutase, and a nucleoside diphosphate sugar epimerase more recently implicated in LOS assembly (36) (Fig. 2D-F).

**Figure 2:**
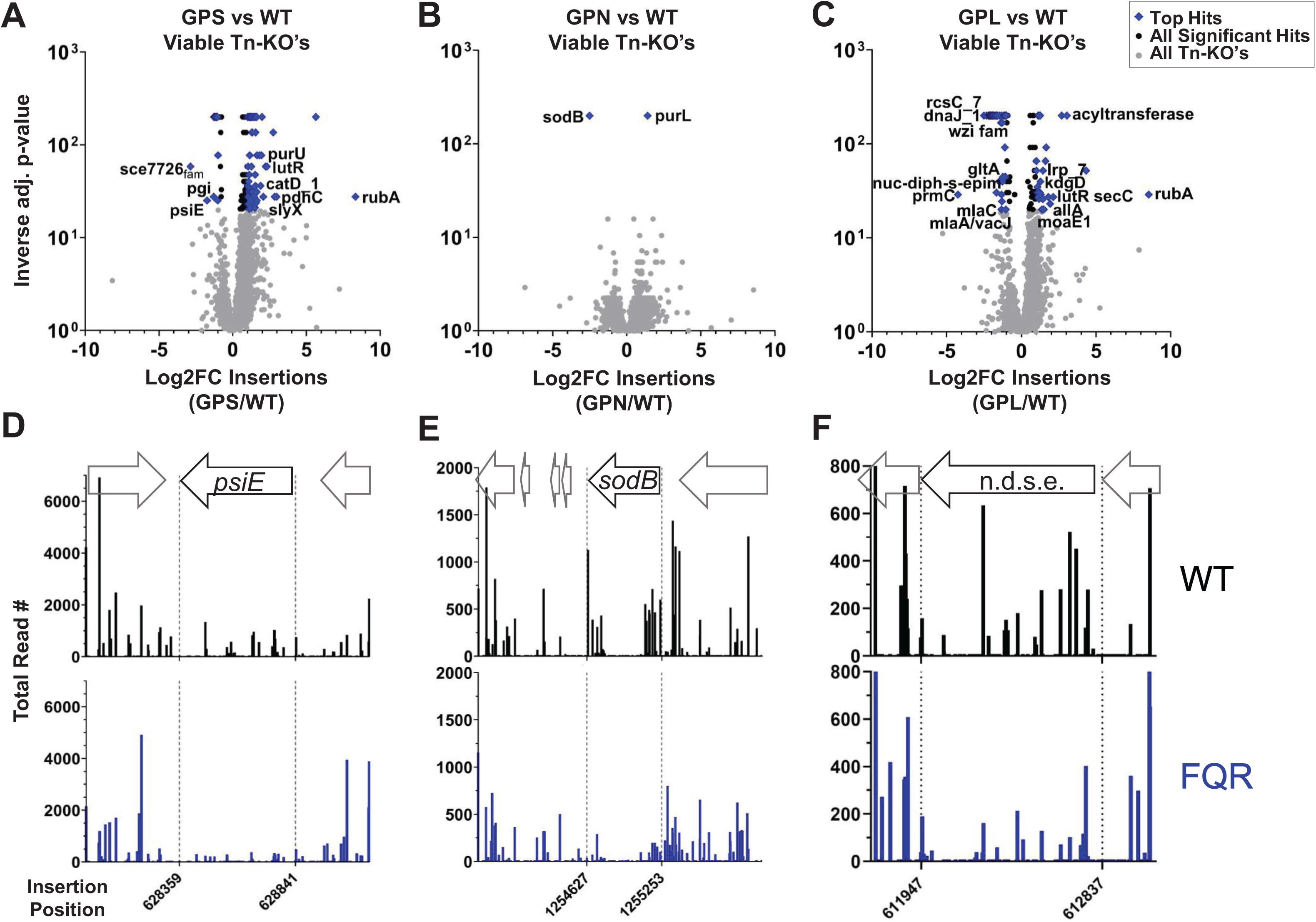
Discovery of genes with lowered tolerance for insertions in the high-level FQR strains. (A-C) Volcano plots in which statistical significance is plotted as a function of the differences in insertion abundance between each FQR strain and WT based on TRANSIT algorithm (33). All adjusted inverse p-values = 0 were assigned the value of 0.005 to allow graphing. Black & blue: genes with adjusted p-values<0.05. Values for all other genes are shown in grey. (D-F) Number of reads plotted as a function of insertion site for noted genes as well as flanking regions. Shown are: FQR: fluoroquinolone resistant. WT: wildtype. GPL: *gyrAparCadeL*. GPN: *gyrAparCadeN*. GPS: *gyrAparCadeS*.

Prior to the exclusion of essential genes, this analysis revealed that *cyoAB* encoding components of the essential ubiquinol oxidase complex harbored significantly fewer insertions in GPL than in WT (Fig. S1A, B). Its function during aerobic respiration as a generator of the proton motive force may play a role in allowing the bacterium to tolerate proton influx resulting from the drug-proton antiport activity of the RND efflux pump (37, 38). To provide support that lowered Cyo activity reduces fitness in the presence of *adeL*ΔI335A336, we used CRISPRi to knock down expression of the *cyoABCDE* operon in the GPL and WT strains and observed growth in broth culture. To this end, an anhydrotetracycline (aTc) –inducible dCas9 was inserted at the *att*Tn7 locus in both strains and transformed with a plasmid harboring a constitutive single guide RNA (sgRNA) (Materials and Methods). As a control, the non-targeting original vector (labeled NC-sgRNA; Table S1) was used. Growth was monitored in cultures grown for 5.5 or 6 hours after dilution to low density in LB and aTc. We saw a fitness defect when expression of the ubiquinol oxidase system was knocked down in both the WT and the GPL strains, with the defect enhanced in WT treated with 100ng/mL aTc (Fig. S1C). Quantification of *cyoABCDE* operon transcript levels after aTc addition showed that expression was silenced in both strains at comparable efficiency compared to controls (Fig. S1D).

Notably, there were very few insertions in both the WT and GPL strains which is an indication of gene essentiality. As a consequence, although the locus was distinguished among those showing the most highly significant difference in insertion efficiencies (Fig. S1B), these differences were measured by degrees of essentiality (Fig. S1E). Furthermore, the average number of viable insertion mutants per *himar-1* transposon location (TA) among the cumulative WT pools was approximately 258 and in GPL about 440. While the loci highlighted from these analyses demonstrate significantly different insertion efficiencies, these differences were likely artifacts of plating efficiency rather than conditional essentiality. Therefore, a second strategy was pursued to measure fitness defects during growth in broth culture to attempt to identify more clearcut differences between the two strain backgrounds.

### The FQR mutants have increased vulnerability to disruption of cell envelope homeostasis

To attack this problem using a second approach, multiple small Tn insertion pools (∼8-16k unique insertions) were constructed in each background, and each was grown for 7-8 generations in the absence of antibiotics to determine the relative fitness levels of individual gene disruptions (>79,000 unique insertions in total for each background; Materials and Methods). The difference in fitness levels between an individual pump overproducer and the wild type was then determined, and statistical significance was plotted as a function of fitness differences in the two backgrounds (Fig. 3A-C). In the absence of drug, the GPS mutant was particularly tolerant of insertion mutations relative to the WT control (Fig. 3A; Data Set S2). For the GPN and GPL strains in the absence of drug, among the proteins identified are those affecting membrane integrity including those that modulate lipid trafficking and LOS synthesis (Fig. 3B-C; Data Set S2). Notably, the GPS strain which shows no upregulation of the outer membrane component of the RND systems was the most tolerant of disruption in these pathways (Fig. 3A).

**Figure 3:**
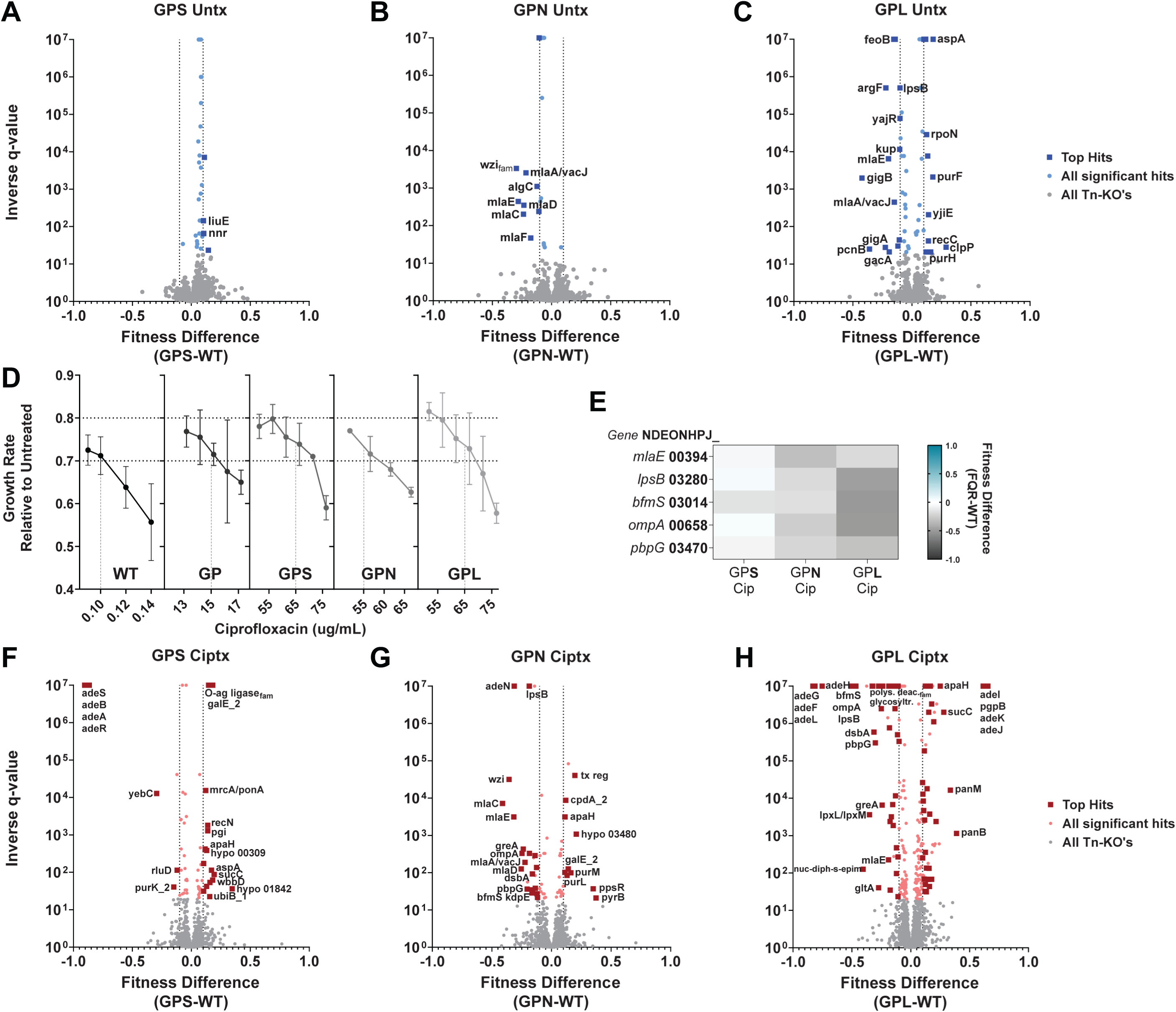
Maintenance of cell envelope homeostasis supports the fitness of efflux pump overproducer strains. (A-C, F-H): Statistical significance plotted as a function of relative fitness comparing identical mutations in FQR and WT strains grown in the absence (A-C) or presence (F-H) of sub-MIC levels of CIP. (D) Growth rates at noted CIP concentrations to identify conditions allowing 20-30% growth inhibition in the FQR and WT strains. (A-C) Blue or (F-H) Red: Mutations above the FDR = 5%, q<0.05 as determined by Benjamini and Hochberg (74). Notable findings are labeled. All q-values < 0.000001 assigned the value of 0.000001 for graphing purposes. (E) Heatmap showing fitness of insertions in each gene in noted strain backgrounds. Selected genes with Tn-disruptions yielding lowered fitness in the FQR strains are shown in the presence of CIP. Gene names are followed by accession numbers which are preceded by “NDEONHPJ_” per the ATCC 17978 strain annotations. This data excludes genes characterized as essential and genes pertaining to prophages or plasmids. Shown are: WT: wildtype. GP: *gyrAparC.* GPS: *gyrAparCadeS.* GPN: *gyrAparCadeN*. GPL: *gyrAparCadeL*.

To determine if antibiotic treatment increased the spectrum of mutations that sensitize pump overproducers, each of the strains was treated at various CIP concentrations to identify conditions that resulted in 20-30% growth rate inhibition (Fig. 3D). Using drug concentrations that reproduced these growth rates, each of the insertion banks was grown 7-8 generations in the presence of CIP (17), cultures were harvested, gDNA was isolated, insertion sites were selectively amplified and sequenced, and the abundance of each insertion was determined using the MAGenTA pipeline (17, 39, 40) (Materials and Methods). From these datasets, the fitness scores of gene disruptions were normalized to a LOWESS curve to avoid the position bias exhibited with CIP treatment (Fig. S2). The differences in fitness levels between an individual pump overexpresser and the wild type were then used to identify mutants that preferentially hypersensitize the pump overexpressing strain to CIP (Fig. 3E-H; Data Set S2).

In general, treatment with drug increased the number of insertion mutations that resulted in fitness defects in the pump hyper expressing strains compared to WT (compare Figs. 3A-C and 3E-H). Primary targets that preferentially sensitize at least two FQR strains during CIP treatment were concentrated in loci related to envelope homeostasis, including *bfmS* which affects a collection of cell envelope processes (41) and *pbpG* which is involved in peptidoglycan degradation (40) (Figs. 3E-H). Even so, the magnitude of defects was consistently higher for mutations in GPL compared to the GPS background, with mutations in the GPN background often showing intermediate effects (Fig. 3E). As in the absence of drug, the GPS strain showed higher tolerance than the GPL and GPN strains for mutations in the same pathways (Fig. 3E-H).

Knockout mutations of candidates were generated as gene deletions in both the FQR and parent strains via homologous recombination to verify their phenotypes (their genotypes confirmed by whole genome sequencing, Materials and Methods). The targets for deletion were chosen based on the following criteria: 1) they resulted in significant fitness defects in at least two of the FQR strains (q ≤ 0.05); and 2) they had severe fitness defects in at least one FQR strain (fitness defect ≥ 0.25). Bacterial growth was then evaluated in the presence of CIP (Figs. 4, 5) at concentrations that yielded comparable growth inhibition across strains (Fig. 3D). Based on Tn-seq analysis (Figs. 3E-H), insertion mutants in *bfmS* and *ompA_Ab_* showed increased CIP sensitivity in GPN and GPL compared to the WT strain background (Fig. 4A). Consistent with these findings, growth defects caused by the deletion mutations were exacerbated in the pump overexpressing strains after CIP treatment as compared to the WT strain background (Fig. 4B). Of particular interest was the behavior of each of the mutants in the GPL background. Both the Δ*bfmS::aacC1* and Δ*ompA* mutants were highly attenuated for growth in the presence of CIP, with growth plateauing far below that observed for controls (Fig. 4B). For instance, when Δ*bfmS::aacC1* was crossed into the GPL background, growth plateau was observed at about 10X lower culture densities than that observed for the GPL parent and 12-20X lower than the same mutation crossed into WT (Fig. 4B). Similarly, when the Δ*ompA* mutation was crossed into GPL, plateaus occurred at 5-8X lower culture density than the GPL parent and about 8X lower than the same mutation crossed into WT (Fig. 4B).

**Figure 4:**
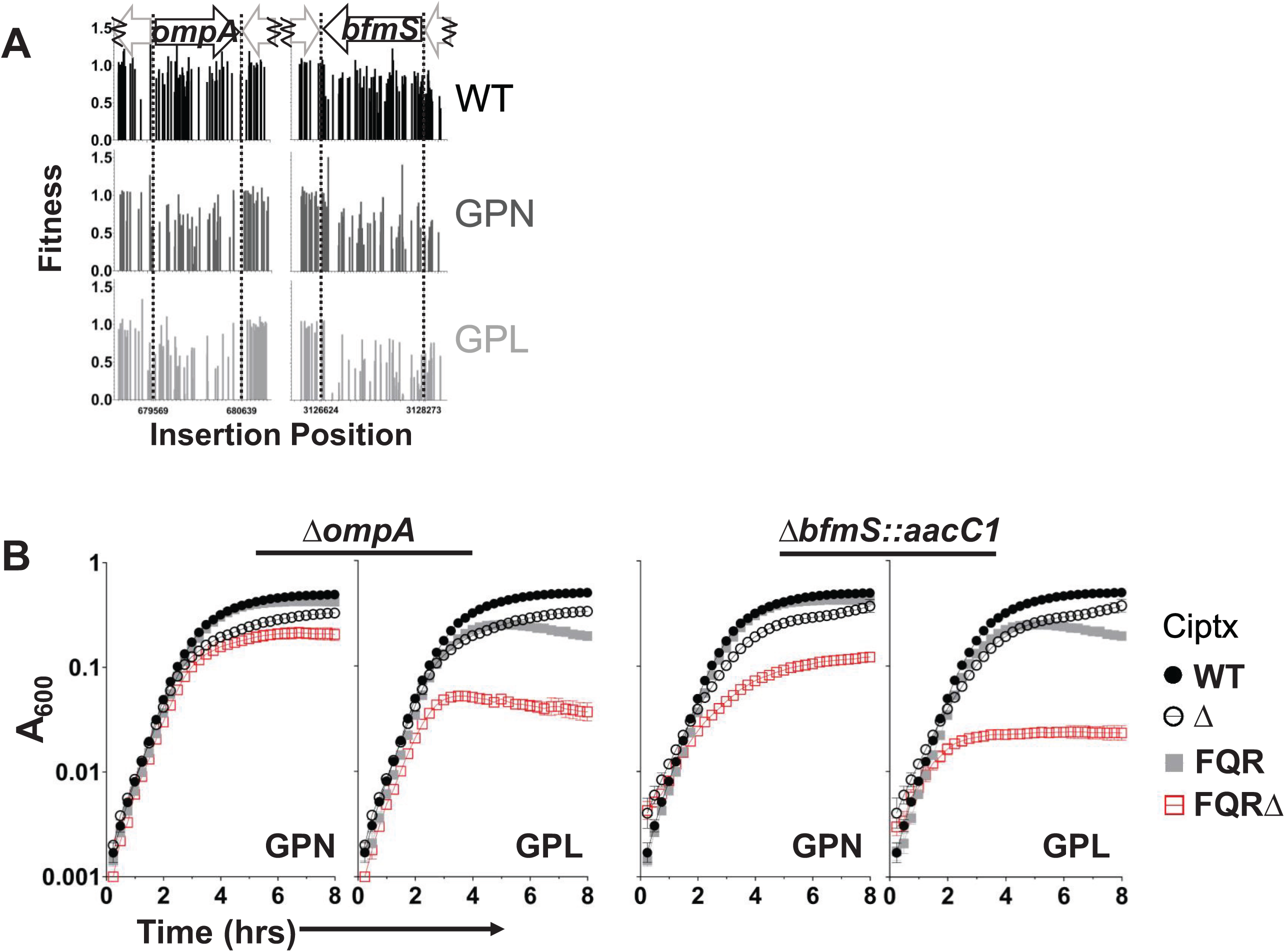
OmpA and BfmS are critical fitness determinants for *adeL*ΔI335A336 strains during CIP exposure. (A) The Single Fitness pipeline of MAGenTA was used to evaluate the fitness of Tn mutants at each position across the loci shown, post CIP treatment. (B) Null mutations in the FQR strain backgrounds grown in the presence of CIP compared to the WT background. Same experimental data for the WT strain and single deletion mutants are shown in each graph for comparative purposes. Growth was monitored by taking absorbance measurements every 15 minutes in 96 well plates in presence of CIP (0.1 µg/ml for the WT, 55 µg/ml for the GPN, and 65 µg/ml for the GPL parent strains and deletion mutants in those respective backgrounds). Displayed are means ± SEM (n≥3). Of note, GPLΔ*ompA* has an additional mutation in *lpxK* (I283V). Ciptx: CIP treated. WT: wildtype. GPL: *gyrAparCadeL*. GPN: *gyrAparCadeN*.

To ensure the growth inhibition was a result of the pump regulator mutation, the consequences of crossing the mutations into the lower-level resistance GP strain were analyzed in comparison to GP carrying a pump-activating mutation in each of the pump regulators (Fig. S3). This allowed a direct comparison between strains differing by a single FQR mutation. The increase in drug sensitivity was more pronounced in the FQR pump mutants carrying the Δ*ompA_Ab_* and Δ*bfmS::aacC1* mutations relative to GP with the same lesions (Fig. S3A).

The pump-overexpressing mutations alter regulatory elements that could be modulating the expression of other unidentified genes that cause fitness alterations. To eliminate this possibility, we determined if the fitness defect of the GPL strain harboring the Δ*ompA_Ab_* and Δ*bfmS::aacC1* mutations was reversed by deleting the *adeFGH* efflux pump genes. Deletion of *adeFGH* reversed the fitness defects caused by the Δ*ompA_Ab_* and Δ*bfmS::aacC1* mutations, consistent with pump overproduction causing the observed fitness defects (Figs. S4 F and S5 F respectively). Furthermore, for each mutation tested, the GP strain background showed comparable fitness to the GPLΔ*adeFGH* strain background (Figs. S4 H & S5 G).

The GPLΔ*ompA_Ab_* strain (Fig. 4B) was found by whole genome sequencing (WGS) to harbor an additional mutation in *lpxK* (I283V), so we created additional strains harboring Δ*ompA_Ab_* via homologous recombination in the GPL background (Materials and Methods) and tested their phenotypes relative to that of GP. The heightened sensitivity of GPL to CIP after introduction of the Δ*ompA_Ab_* lesion was retained (Fig. S4 B-D), indicating that the fitness defect was due to the loss of OmpA. Similarly, the GPΔ*bfmS::aacC1* strain in Fig. S3A was found to have an additional mutation upstream of the *bfmR* gene. This mutation dampened the expression of *bfmR* (Fig. S5H; strain #272 GPΔ*bfmS::aacC1* (Fig. S3A) compared to strains #271 GPΔ*bfmS::aacC1* (Fig. S5B) and #304 Δ*bfmS::aacC1* (Fig. 4B)). Additional clones cured of this *bfmR* promotor mutation were created in the GP strain background, and knockout of *bfmS* reproducibly showed minimal fitness alterations of the GP parent in the presence of CIP, while still sensitizing the *adeL* variant that overexpresses the efflux pump (Fig. S5B-E).

Given the strong defects observed with these two mutants, other candidate mutations predicted to interfere with envelope biogenesis were interrogated, including *lpsB* and *pbpG*, which showed lowered insertion mutation fitness in the pump hyper-expressors when compared to WT, particularly in the GPL strain background (Fig. 5A). Deletion mutations were constructed in WT, GP, GPN and GPL backgrounds, and growth of the eight mutant strains was monitored in the presence of CIP. Loss of PbpG from either the GPN or GPL strain background was particularly detrimental to growth in the presence of low CIP concentrations (Fig. 5). In both GPN and GPL backgrounds harboring Δ*pbpG*, strains initiated growth but culture densities stopped increasing after about 3 hours (Fig. 5B; Fig. S3B). In contrast, the parental GP mutant harboring Δ*pbpG* did not show this phenotype (Fig. S3B). In the case of the Δ*lpsB* mutant, growth plateaued at lower levels when the deletion was analyzed in the GPL mutant (8X lower than parent GPL) compared to GPN (5-7X lower than GPN), consistent with the *adeL*(ΔI335A336) mutation being particularly susceptible to envelope stress (Fig. 5B).

**Figure 5:**
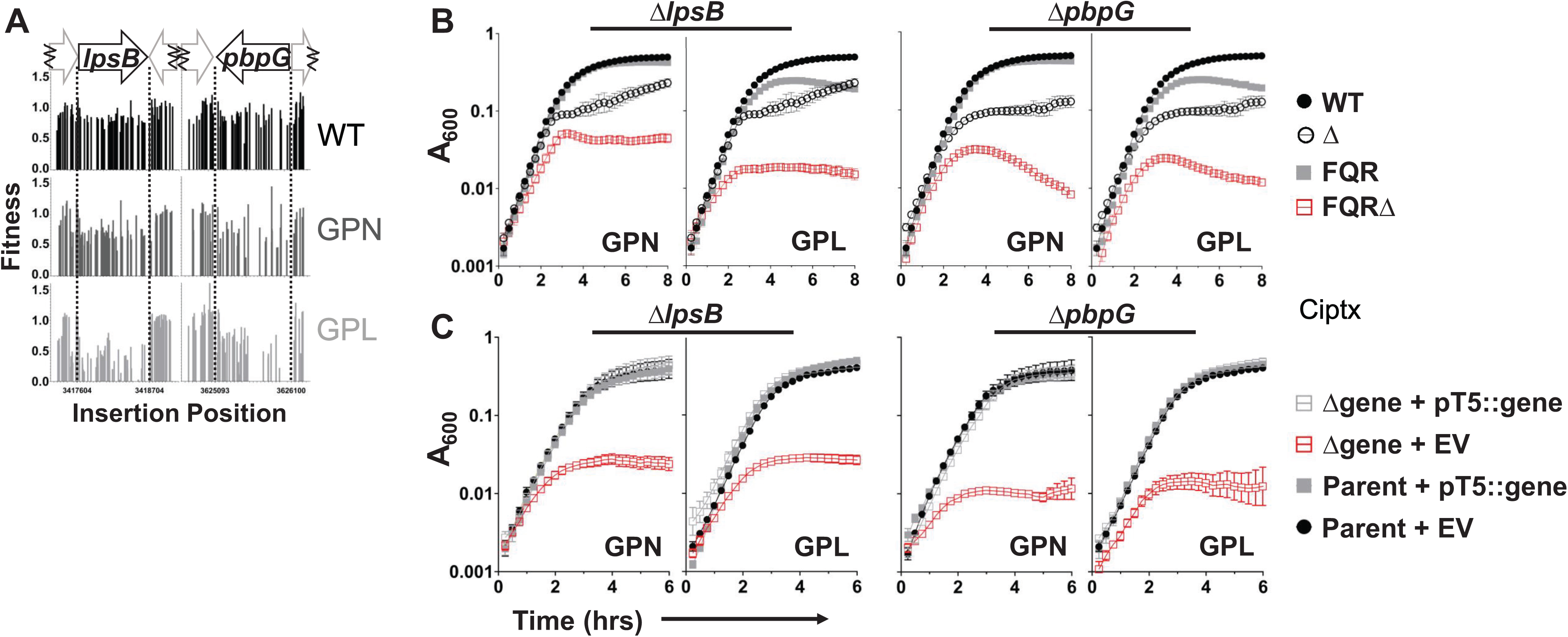
LpsB and PbpG are required for the AdeN (GPN) and AdeL (GPL) mutants for optimal fitness in presence of CIP. (A) Transposon mutant fitness post CIP treatment at each insertion position across the selected loci. (B) Growth of the indicated strains in the presence of CIP was monitored as in Fig. 4, displayed as mean ± SEM (n≥3). Data for the WT strain and single deletion mutants from identical cultures are shown in each graph for comparative purposes. (C) Growth of deletion mutants in LB + 65 µg/ml CIP + 1mM IPTG to induce expression of the indicated genes complemented *in trans* in GPN (column 2) or GPL (column 4) backgrounds. Data are means ± SEM (n≥6). Ciptx: CIP treated. IPTG: Isopropyl β-d-1-thiogalactopyranoside. WT: wildtype. GP: *gyrAparC.* GPL: *gyrAparCadeL*. GPN: *gyrAparCadeN*. EV: empty vector.

To ensure that this phenotype was due to these null mutations, both deletions were complemented *in trans*. The presence of plasmids harboring these genes restored the fitness of both pump overexpression strains, with growth in the presence of CIP indistinguishable from the appropriate parents harboring empty vector (Fig. 5C). WGS of these strains uncovered an *mrcB* (G539C) mutation in the GPNΔ*lpsB* background. Complementation of the GPNΔ*lpsB* strain, however, restored growth to levels of the parental strain, indicating that the growth defect was caused by loss of *lpsB* and not due to the secondary missense mutation (Fig. 5C). Furthermore, when a deletion of the *adeFGH* efflux pump operon was introduced into the GPL strain harboring either the Δ*lpsB* or the Δ*pbpG* mutations, both growth defects were alleviated, indicating that defective growth was linked to the overproduction of the efflux pump (Figs. S6-7).

The absence of PbpG or LpsB has been associated with alterations in *A. baumannii* LOS homeostasis (36, 40). Therefore, LOS profiles of strains disrupted in *pbpG* and *lpsB* were determined by fractionation on SDS-polyacrylamide gels (Fig. S8; Materials and Methods). No significant differences in LOS profiles were detected between the WT or FQR strain backgrounds (Fig. S8A). For the various strains harboring the Δ*pbpG* mutant, there were decreased levels of a species unrelated to LOS, but otherwise the migration and relative concentration of LOS species appeared unaffected (Fig. S8B). In contrast, as expected, Δ*lpsB* mutations resulted in truncated LOS, with similar profiles in each of the efflux pump backgrounds (Fig. S8C). Therefore, LOS is altered by these mutations as previously described, and these alterations were independent of the strain background tested (40).

As reduced A_600_ yields could be indicative of morphological differences between mutants, the various strains were imaged. As expected, the *bfmS*::*aacC1* and Δ*pbpG* mutants showed morphological alterations that have been previously documented, ranging from stubbier to slightly rounded and enlarged, dependent on the mutation (41, 42). The latter was observed with Δ*lpsB* as well. These alterations were independent of pump overproduction (Fig. S9). In contrast, the Δ*ompA* mutation resulted in no morphological changes (Fig. S9). In response to CIP, the parental strain elongated as anticipated (43, 44), while the mutations that showed morphological changes continued to appear different from their parent strain, with changes ranging from reduced length to wider cells (Fig. S10). While structural deformation from loss of PbpG appears slightly more severe in the pump overexpressing strain background (Fig. S11), the specific morphological alterations caused by the mutants were observed both in the presence and absence of pump hyperexpression (Figs. S11-14). Critically, what differentiated the morphotypes of these lesions between the pump overexpressing strain and parent strain is that cells in the pump overexpressing strain background consistently displayed larger volumes (Figs. S11-14). We conclude that although the mutations that resulted in enhanced CIP sensitivity in pump overproducer strains resulted in altered morphologies, it is unlikely these changes in morphology explain for the reduced culture densities observed in growth curves of strains that overproduce efflux pumps (Figs. 4 and 5).

### Maintenance of pH homeostasis in the presence of pump overproduction

The RND-class efflux pumps are known to function as drug-proton antiporters (45). With this in mind, we investigated whether FQR pump hyper-expression causes *A. baumannii* cytosolic pH to become more acidic in the presence or absence of CIP substrate. WT, the high-level FQR strains, and the GP precursor strains were grown to mid-exponential phase and loaded with BCECF-AM dye to determine internal pH (pHi). The pHi values were then determined from a standard curve of cells incubated in pH-adjusted buffers in the presence of CCCP to equilibrate external and internal pH (Materials and Methods). In the absence of antibiotic, the cytosol of GPN, the AdeIJK pump over-expressor, was significantly more basic than the GP parent strain (Fig. 6A), while the cytosolic pH values of the other pump overproducers were otherwise comparable to GP or WT controls (Fig. 6A; one-way ANOVA, multiple comparisons between all strains and GP, * where p<0.05). To determine if the presence of CIP, which should result in proton-CIP antiport, affects internal pH, the experiment was repeated after exposure to CIP at concentrations yielding comparable levels of growth inhibition across strains. CIP treatment lowered the pHi compared to the untreated conditions for all strains. The cytosols of the GPS and GPL strains were fractionally more acidic than the parental GP strain after CIP addition, but the differences did not rise to the level of statistical significance (Fig. 6B; same tests as in 6A). Imaging shows these strains exhibit similar cell morphologies both in the absence (Fig. S15) and presence of CIP (Fig. S16). These data indicate there may be homeostatic mechanisms to neutralize the effects of pump overexpression that act at either transcriptional or post-transcriptional levels, so the global transcriptional effects of pump overexpression were analyzed in the presence and absence of CIP (Fig. 1A, third approach).

**Figure 6:**
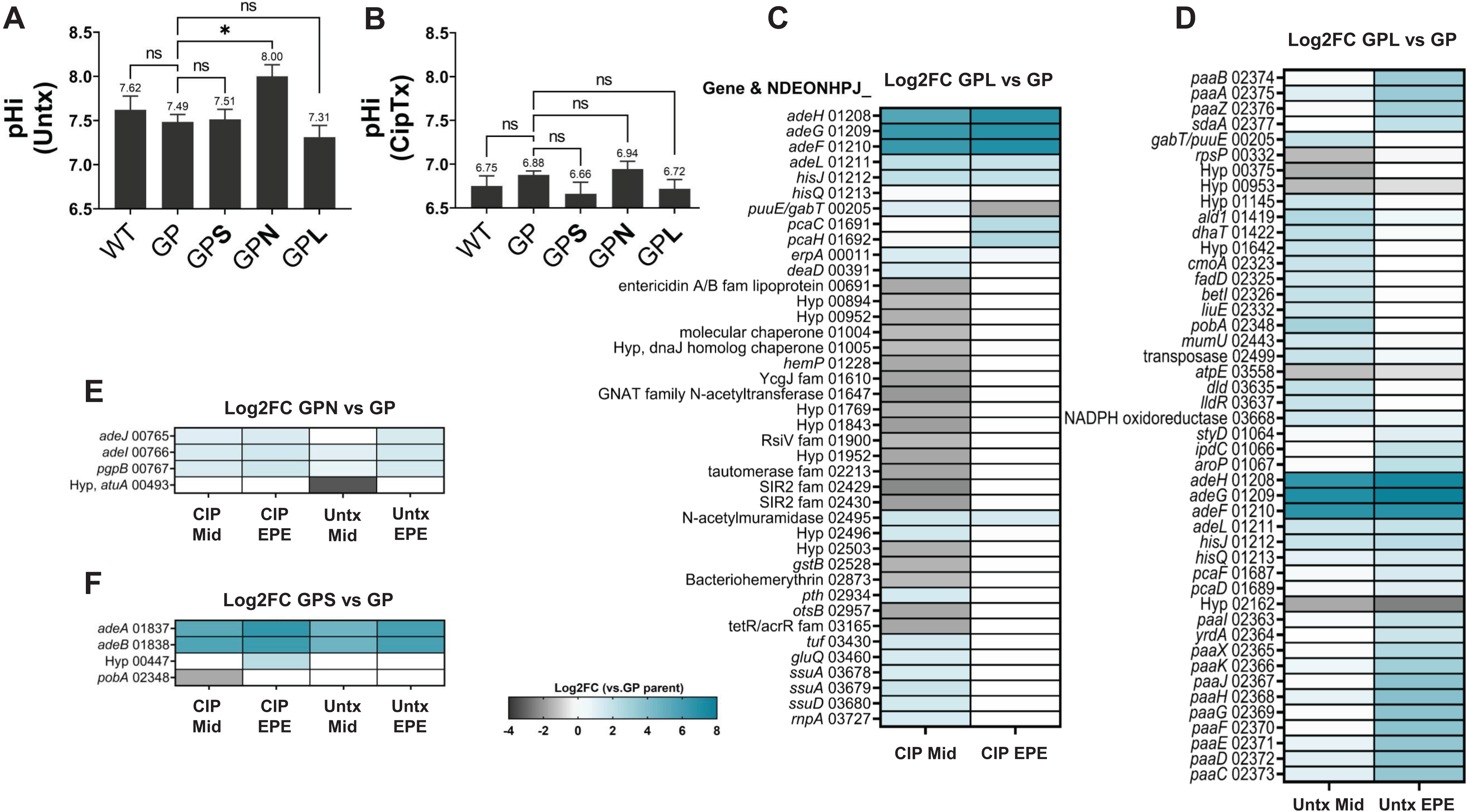
Transcriptional and physiological response to pump overproduction. (AB) Alterations in intracellular pH (pHi) in response to pump overproduction. Noted strains grown to mid-exponential phase in the absence (A) or presence (B) of CIP. Internal pH (pHi) was measured using 20µM BCECF-AM as described (Materials and Methods). Displayed are the means ± SEM for each strain from n=5-8 biological replicates, with P values derived from one-way ANOVA making multiple comparisons. * = p < 0.05 (0.0259); ns=not significant. (C-F) Transcriptional alterations in response to pump overproductions are magnified in presence of *adeL*ΔI335A336 mutation. Transcriptomic data (n=3) was analyzed comparing high-level FQR pump hyper-expressing strains and the GP parent strain (Materials and Methods). Differences in transcript levels for each comparison are shown via the heat maps with GP vs GPL (C, +CIP) and (D, –CIP), GP vs GPN (E) and GP vs GPS in (F). This data excludes genes pertaining to prophages or plasmids. A gene is listed if a significant change in gene expression was observed in at least one condition within the panel. WT: wildtype. GPL: *gyrAparCadeL.* GPN: *gyrAparCadeN*. GPS: *gyrAparCadeS*. GP: *gyrAparC*. Mid: mid-exponential phase. EPE: early post-exponential phase.

### Large transcriptional changes in response to pump overproduction are limited to the AdeL-regulated pump

To determine if pump overproduction results in transcriptional changes that support fitness, the FQR, low-level resistance GP, and WT strains were cultured in the absence or presence of CIP at concentrations causing 20-30% growth inhibition and analyzed by RNAseq (Materials and Methods). Samples were collected during exponential phase (A_600_ ∼0.5) as well as early post-exponential phase (A_600_ ∼1), and differentially regulated genes were identified by comparing the high-level FQR strains and the GP parent strain (Fig. 6C-F; GP vs GPL (C, +CIP) and (D, –CIP), GP vs GPN (E), and GP vs GPS (F), Data Sets S3 and S4). In either the presence or absence of CIP, the GPL strain had the greatest number of differentially expressed genes relative to the control GP strain when compared to the other pump-overexpressing strains (Fig. 6C-F). In fact, hyperexpression of either the AdeS-or AdeN-regulated pumps only altered the gene expression patterns of very few genes in the presence or absence of CIP (Fig. 6E, F). This result echoes the Tn-seq analysis of the three overproducer strains, as the *adeL* mutant had the largest set of mutations that lowered fitness relative to WT (Figs. 2, 3, and Figs. S17-19).

Among the various transcriptional changes resulting from AdeL-regulated pump hyper-expression, the most notable was upregulation of the *paa* operon, which encodes proteins involved in degradation of phenylacetate (Fig. 6C, D). In the absence of drug, 14/46 genes that were differentially regulated in the two backgrounds were from the *paa* regulon (Fig. 6D). Furthermore, in early post-exponential phase, genes rank-ordered as the 12 most highly overexpressed in the GPL strain compared to the GP parent were also in the *paa* regulon (Supplemental Dataset 4), so the functional importance of this upregulation in the absence of drug was pursued further. As confirmation of the overexpression of *paa*, RNA was isolated from parallel cultures of GP and GPL grown in the absence of drug and analyzed by q-rtPCR (Materials and Methods). Consistent with the RNAseq results, transcript levels of the first gene of the operon, *paaA*, were significantly elevated in the GPL strain compared to the GP parent, with a 23-fold increase during exponential phase and a 6-fold increase in early post-exponential phase (Fig. 7A, B). Another transcriptomic change of note in the GPL strain was the downregulation of the *atpE* gene encoding a component of the F_1_F_0_ proton translocating ATPase/synthase in the absence of drug (Fig. 6D). This parallels other work, which shows that depletion of the ATPase/synthase increases fitness of the GPL strain in the presence of CIP (35).

**Figure 7:**
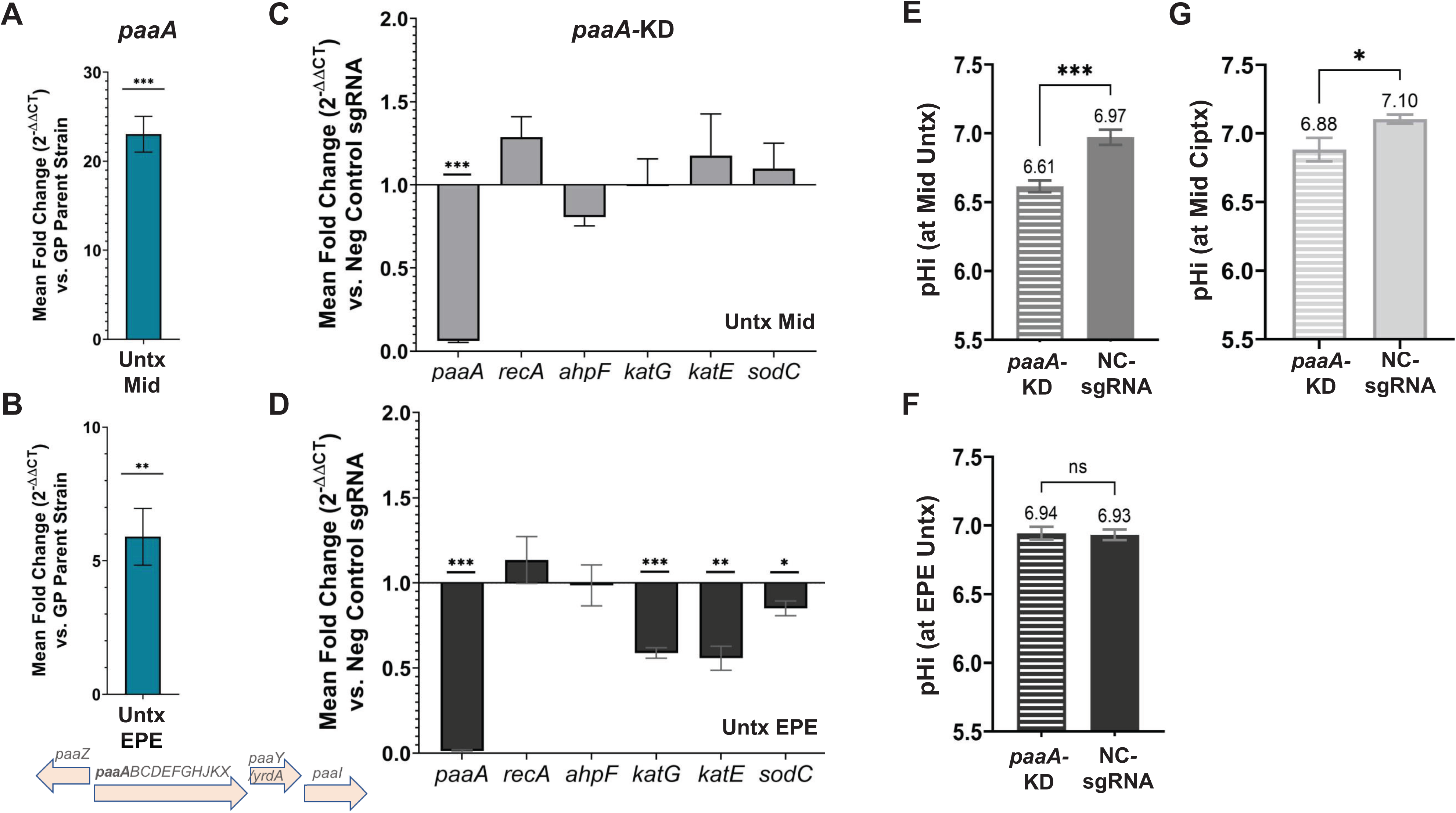
The presence of *adeL*ΔI335A336 in GPL results in overexpression of the *paa* operon. RNA was isolated from GP and GPL strains and transcription of *paaA* gene was quantified by q-rtPCR. Strains grown to exponential (A) or early post-exponential phase (B) in absence of drug. Data are mean fold change in transcript levels (2^-ΔΔCT^) ± SEM of each target (n=3) compared to parental GP strain, analyzed by two-tailed t-test. Significance marked by asterisks; **, p=0.0099; ***, p=0.0004. (C, D) Alterations in transcript levels in CRISPRi-depleted *paaA* strain, with knockdown verified as described (Materials and Methods). RNA was prepared from cultures grown to mid-exponential (C) or early post-exponential phase (D). Data are transcript levels of each target relative to nontargeting negative control (2^-ΔΔCT^) ± SEM (n=3), analyzed by multiple unpaired t test analyses following the two-stage step-up Benjamini, Krieger, and Yekutieli, false discovery rate of 5.00% (75). Significance: *, p=0.020-0.034; **, p=0.002-0.003; ****, p<0.001. (E-G). Acidification of cytosol as a consequence of *paaA* depletion. Internal pH (pHi) determined during mid-exponential phase in absence (E, F) or presence (G) of CIP. (F). Intracellular pH in early post-exponential phase. Data are mean ± SEM from n=5. P values were derived from an unpaired t test comparing the mean pHi values between the two strains. (***, p=0.0009 in E; *, p=0.0420 in G) & (ns, p=0.8479 in F). Untx: Untreated. Ciptx: CIP treated at 45µg/mL. Mid: grown to mid-exponential phase. EPE: grown to early post-exponential phase. GPL: *gyrAparCadeL*. GP: *gyrAparC*.

### Depletion of *paaA* alters expression of stress-related genes and lowers intracellular pH in the GPL pump hyperexpressor

The phenylacetate catabolic pathway breaks down the substrate phenylacetate to generate end products that feed into the TCA cycle (46). Previous studies indicate that induction of the *paa* pathway in *A. baumannii* may be a response to stress observed after antibiotic treatment (47). Therefore, upregulation of this operon may similarly support overproduction of the AdeL-regulated pump. It should be emphasized that *paa* mutants showed no significant fitness defect in our TnSeq screen. Consistent with the Tn-seq findings, targeting *paaA* by CRISPRi caused no growth defect in the GPL strain as compared to a parallel non-targeting control strain. Depletion was efficient, as there was a 20-fold knockdown in exponential phase and almost 200-fold in early post-exponential phase compared to the negative control (Fig. 7C, D). This argues that in the absence of phenylacetate degradation, other compensatory pathways may allow fitness to be maintained.

The *paa* pathway has been shown to promote resistance to hydrogen peroxide (47, 48), so enhanced expression of the *paa* pathway in the AdeFGH pump hyperexpresser may facilitate a response to oxidative stress and/or other stressors. To test this idea, transcription of a set of oxidative stress genes was probed in the GPL strain by q-rtPCR analysis. Knockdown of *paaA* expression in early post exponential phase yielded reduced transcript levels of *katG, katE*, and *sodC* (encoding two peroxidases and superoxide dismutase, respectively; Fig. 7D). This diminished expression is consistent with previous observations that *paa* null mutants are more sensitive to hydrogen peroxide stress through some previously uncharacterized mechanism (47, 48).

Internal pH was determined after *paaA* knockdown in the GPL mutant (Fig. 7E, F). Knockdown of *paaA,* the first gene of the large operon, resulted in a significant acidification during exponential growth compared to a non-targeting control (Fig. 7E) (A_600_ ∼0.52-0.62). By early post-exponential phase (A_600_∼1.22-1.39), the effect of the knockdown was no longer statistically significant compared to the non-targeting control (Fig. 7F). When the *paa* knockdown cultures were grown to exponential phase and exposed to sub-MIC levels of CIP for the last hour, the pHi was lower than the non-targeting control (Fig. 7G; significance determined via unpaired t tests). These data indicate a role for this pathway in maintaining pH homeostasis during exponential growth of strains overexpressing the AdeL-regulated pump.

Given that the *paa* operon has been documented to be associated with a drug stress response, we determined if the expression of the *paa* operon is altered in response to CIP in each of the strains. Based on the RNAseq data in exponential (Fig. 8A) and early post-exponential (Fig. 8B) phase, *paaA* transcript levels in all the strains were significantly upregulated in response to CIP exposure as compared to growth in the absence of drug (Fig. 8A, B). These results are consistent with previous results showing that upregulation of the *A. baumannii paa* catabolon occurs in response to drug stress (47). Notably, upregulation of *paa* transcript levels in the absence of drug is a special feature of the GPL strain when compared to the GP parent, and this is not observed in the GPS or GPN strains (Fig. 6D).

**Figure 8:**
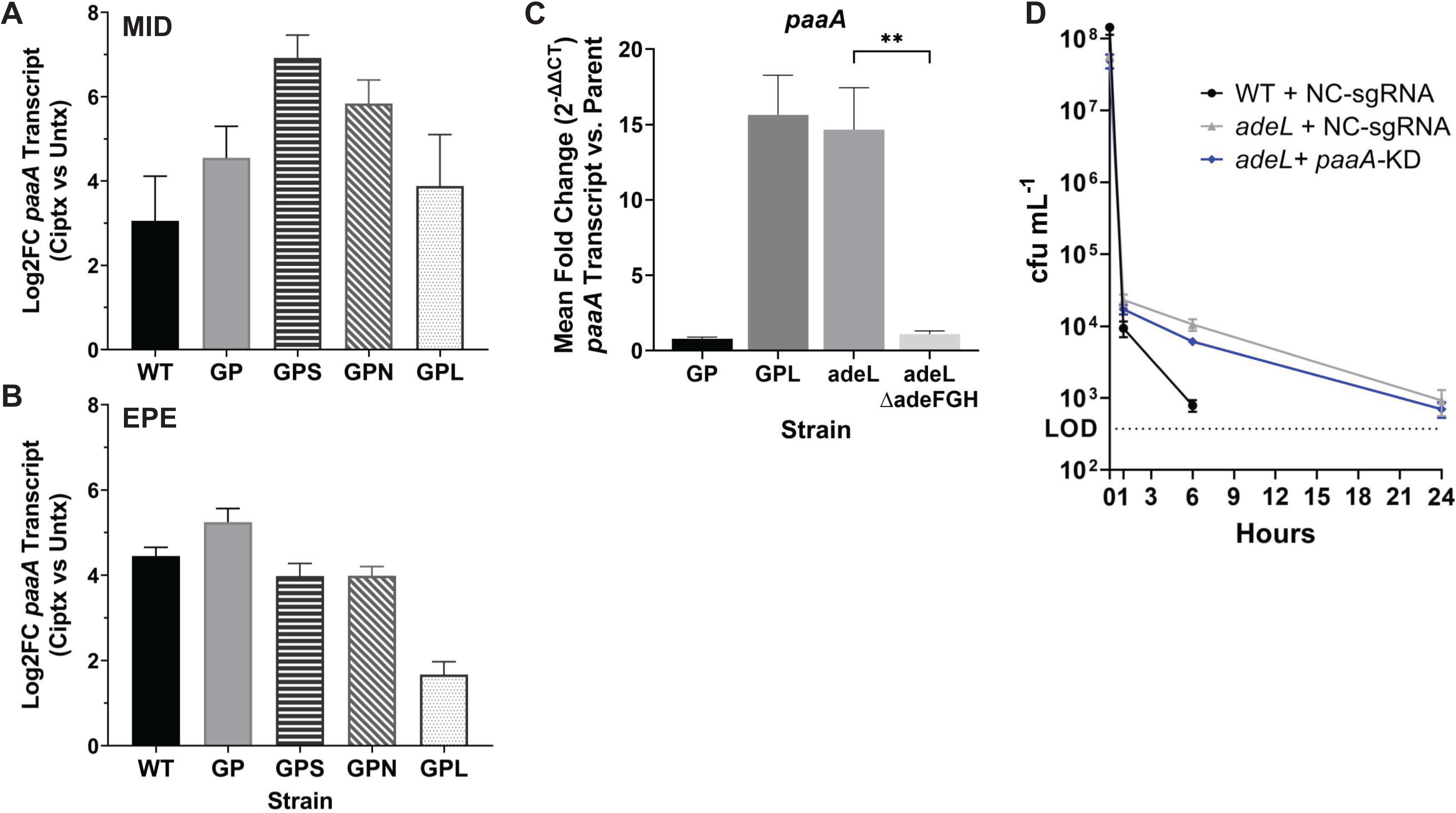
*paaA* is significantly upregulated during CIP exposure. (A,B) The *paaA* transcript (shown with L2FCSE) is upregulated after CIP exposure independently of strain background. Strains were grown to noted growth phases, with data derived from RNAseq experiments (Fig. 6; Materials and Methods). Significance determined by s values from triplicate cultures for each (s ≤ 1.11E-03). (C) *paaA* upregulation is pump dependent. Strains were grown to mid exponential phase and their RNA were used to quantify *paaA* transcript levels by q-rtPCR. Bars show mean transcript levels (2^-ΔΔCT^) ± SEM (n=3) relative to parent strains (WT or GP), significance determined by Ordinary one-way Anova (**, p=0.0027). (D) Persistence of *adeL*ΔI335A336 single mutant (in WT strain background) is unaffected by depletion of *paaA*. The indicated strains were grown to exponential phase with aTc inducer to deplete *paaA* transcript, then exposed to CIP at ∼20X MIC for the noted times prior to plating on LB agar plates in absence of drug to determine CFU/mL. The mean CFU/mL ± SEM are shown (n=3). The dotted line indicates LOD, the limit of detection for the assay (375 CFU/mL). Mid: grown to mid-exponential phase. EPE: grown to early post-exponential phase. aTc: anhydrotetracycline. WT: wildtype. GPL: *gyrAparCadeL*. GPN: *gyrAparCadeN*. GPS: *gyrAparCadeS*. GP: *gyrAparC*.

To determine if this upregulation is in response to overexpression of the RND efflux pump, rather than a consequence of a mutation in the regulatory element, the AdeL single mutant and AdeL mutant with the AdeFGH pump deleted were grown to mid-exponential phase with WT, GP, and GPL cultured in parallel. RNA was extracted and used for q-rtPCR to quantify *paaA* transcript levels. As anticipated, *paaA* transcript levels were elevated in the AdeL mutant. These levels dropped, however, in the absence of the AdeFGH RND efflux pump (Fig. 8C). This result supports the hypothesis that the *paa* operon is involved in a stress response to efflux pump hyperexpression.

Previous work has shown that pump hyper-expression and the oxidative stress response are contributors to the development of persistence in the presence of supra-MIC levels of antibiotics (49–52). In particular, the single AdeL mutant promotes persistence in the presence of about 20X MIC levels of CIP (52). We, therefore, determined if depleting *paaA* expression would interfere with persistence. Depleting *paaA* transcript did not alter the ability of a single *adeL* mutation in the WT background to persist in high levels of CIP when compared to the non-targeting control in AdeL (Fig. 8D). These results indicate that during exponential growth, upregulation of phenylacetate metabolism plays a role in maintaining cellular homeostasis in response to pump expression, mimicking the response of *A. baumannii* to antibiotic exposure.

## Discussion

In this work, we set out to identify proteins that were required to maintain the fitness of *Acinetobacter baumannii* drug efflux pump-activated strains. The most surprising result was that in the AdeN and AdeS mutants, efflux pump overexpression was well-tolerated, with few mutations causing defects specific to these strain backgrounds (Fig. 3) and little compensatory transcriptional changes (Fig. 6). In contrast, the high-level fluoroquinolone resistant (FQR) strain expressing the AdeL-regulated pump showed a range of mutations that lowered fitness relative to the WT control and a more pronounced shift in its transcriptome relative to a *gyrAparC* FQR mutant (GP). An obvious difference between these mutants is that the *adeL*-activating mutations resulted in a higher level of expression of the AdeH outer membrane component than the corresponding AdeS– and AdeN-regulated outer membrane components (Fig. 6). The AdeS-regulated pump that was the most tolerant of insertion mutations has an outer membrane component that is not subject to AdeS-mediated upregulation (31, 32).

We previously observed that AdeL mutations confer persistence during high-level CIP exposure (52). In the mouse pneumonia model, *adeL*-activating mutants are outcompeted by strains that have increased fitness in the absence of antibiotic (52), consistent with *adeL-* activating mutations imparting a fitness cost. We suspect that the lowered growth rate is a consequence of envelope stress (Fig. 3) and could have clinical corollaries. The inability of *adeL* mutants to efficiently compete in immunocompetent animals (52) may mirror drug resistance evolution in hospitals, where immunocompromised patients may act as reservoirs for these variants that eventually act as evolutionary precursors to more fit and higher resistance variants. In this fashion, evolutionary precursors that cause cell envelope disruptions may be compensated by further mutations that allow spread in individuals with intact immune systems.

Fitness costs and resistance drivers are important for the evolution of MDR clinical isolates and it is common for a pathogen to employ multiple drug efflux systems at a time, which has the potential for accumulated envelope stress (19). Of note, while it appears there is no advantage to overexpressing both AdeIJK and AdeFGH efflux systems simultaneously during CIP challenge (Figs. 3H, S17, GPNL mutant in Fig. S20), this was not investigated further in the presence of other drugs nor with AdeAB. This work does emphasize, however, that there is an interplay between fitness costs and fractional differences in drug resistance, and that strains having multiple antibiotic efflux systems may require compensatory mutations to reverse fitness costs. The work described here identified points of vulnerability in the high-level FQR strains that may be exploited to sensitize MDR strains to drug. It is evident that GPL heavily relies on determinants that stabilize the cell envelope and an effective alarmone response in the presence of drug. Furthermore, it may explain why *adeS* and *adeN*-regulated pump overproducers appear to be particularly common in clinical isolates.

BfmS, identified in our Tn-seq screen, is the histidine kinase sensor that negatively regulates the BfmR response regulator in the BfmRS two-component regulatory system (TCS). BfmR controls a large transcriptional network including proteins linked to cell wall biogenesis, capsule production, and outer membrane protein assembly (41). It is intriguing that disruption of *bfmS* is detrimental to the fitness of the RND pump hyper-expressors, because disruption in a drug-sensitive isolate has been shown to increase drug resistance and the expression of genes such as *pbpG* that support RND pump expression (41). Presumably, proteins known to be preferentially downregulated or mis-localized when the TCS is activated by loss of BfmS may, in bulk, contribute to the fitness of the FQR strains. One such downregulated gene is *ompA*, encoding a major outer membrane protein of *A. baumannii* (53, 54), which apparently functions as a low-efficiency non-selective ion channel similar to *E.coli* OmpA and *Pseudomonas* OprF (55, 56).

If there is a single feature linking together the Tn-seq mutant candidates, it is that they hit proteins associated with maintenance of outer membrane integrity. Various studies have demonstrated that defects causing instability to the outer membrane can be overcome by compensatory changes in envelope components, particularly associated with the outer membrane (57, 58). The functionality of the RND transporters is likely to be similarly supported by these components. We postulate that PG, LPS, and other OMPs are central players in allowing overproduction of the outer membrane component of the RND pump while simultaneously supporting outer membrane barrier integrity. As it provides a selective permeability barrier against antimicrobials, any destabilization of this barrier may simultaneously depress drug efflux pump function or lead to a compromised drug barrier, leading to drug sensitivity. Most of the insertions that caused fitness defects in pump-overexpressing strains were revealed only after the addition of CIP (Fig. 3). This means that either of these two mechanisms could come into play, with high pump concentrations combined with a partially dysfunctional outer membrane barrier collaborating to drive drug sensitivity.

There were gene disruptions that yielded significantly higher fitness in all FQR strain backgrounds when compared to the drug-susceptible WT strain, particularly after CIP exposure. The most striking knockouts were in *apaH,* encoding a dinucleoside polyphosphate hydrolase. ApaH degrades (p)ppGpp alarmones, which accumulate in response to stress as a consequence of RelA synthetase activity (59–61). Consistent with this result, we have recently shown that CRISPRi depletion of SpoT, which also degrades alarmones, increases fitness of GPL during CIP treatment (35). Further supporting the model that (p)ppGpp accumulation increases fitness during CIP treatment, transposon insertions in RelA, predicted to eliminate alarmone synthesis, resulted in a fitness reduction in the GPL background (Fig. S17A, C). Therefore, it is likely that accumulation of signals activating a stress response increase fitness specifically in the RND hyper-expressing strains, particularly in GPL.

The necessity of alarmone synthesis for optimum fitness was consistent with our observation that there were spikes in insertion mutation abundance at two prophage locations post-drug exposure in all strain backgrounds (Figs. S17-19C). These peaks were more prominent in the FQR strains, especially GPL, than in the WT strain and were responsible for many of the overrepresented insertions in these backgrounds (Fig. 3). Our previous work indicates that the abundance of these sequences is due to prophage induction after CIP exposure (17), and is associated with an amplified SOS stress response. Therefore, RND overproduction, particularly in the GPL strain, requires an effective stress response in the presence of CIP under conditions in which all strains are growing at equivalent rates.

Analysis of the transcriptome of the RND overproducers provides further evidence for the induction of cell stress as a consequence of AdeL-regulated pump overexpression (Fig. 6). Most notably, transcription of the phenylacetate operon (*paa*) was induced in response to pump overexpression in the absence of antibiotic treatment. This catabolon has been documented to protect against antimicrobial treatment (47) and supports virulence in *A. baumannii*, *Burkholderia cenocepacia,* and *Pseudomonas aeruginosa* (47, 62–64). Although the consequences of loss of *paaA* expression appear mild, there may be redundant pathways that come into play in the presence of drug pressure to support fitness of the AdeL mutant. The absence of *paaA* led to lowered cytoplasmic pH (Fig. 7E, G), so the production of substrates from the Paa pathway, which feed into the TCA cycle, could provide relief from proton influx produced by the overproduced pump. Alternatively, accumulation of phenylacetic acid could lead to lowered cytosolic pH upon knockdown of the *paa* operon. The importance of maintaining pH homeostasis in the presence of pump overproduction is further emphasized by the result that loss of the proton-pumping ubiquinol oxidase complex was predicted to cause lowered fitness of the GPL strain.

In summary, this work indicates that targets involved in supporting the integrity of the *A. baumannii* envelope are associated with maintaining fitness of strains that overexpress efflux pumps. The most highly expressed pump, regulated by AdeL, was associated with the largest number of genetic requirements for fitness, and was the most disruptive to the transcriptome. The genetic loci identified can be viewed as drug targets for synergy with fluoroquinolones as well (Fig. S21), defining pathways that may buffer the stress that accompanies increased pump activity. Still unclear is how the phenylacetate degradation pathway is induced in response to the stress identified here, and what physiological role, if any, it plays in supporting the fitness of the strains analyzed in this study. Future work will attempt to expand these studies into analyzing the role of essential proteins in supporting pump overproduction in hopes of identifying potential targets for multidrug therapy targeting drug-resistant mutants.

## Materials and Methods

### Bacterial strains and growth conditions

Bacterial strains used in this work are listed in Table S1. All *Acinetobacter baumannii* strains are derived from ATCC 17978UN. Cultures were grown in lysogeny broth (LB) (10 g/L tryptone, 5 g/L yeast extract, 10 g/L NaCl) or on LB agar plates (LB supplemented with 15g/L agarose) and incubated at 37°C with aeration, as well as shaking or on a roller drum if grown in broth. Unless grown the same day, cultures were typically diluted, incubated to recover after overnight growth, then diluted to A_600_=0.1-0.001 into either 5-10mL broth or into 96-well plates (Costar), and growth was monitored by measuring A_600_ every hour (broth culture) or every 15min (Epoch 2 or SynergyMx, (Biotek) plate readers.) Growth is then plotted as log-transformed absorbance values, with the mean ± SEM shown. Superoptimal broth with catabolite repression (SOC) was used for recovery following all electroporations of *E. coli* and *A. baumannii* during cloning or library preparations. Electroporations of *A. baumannii* were performed using 0.1cm gap cuvettes (0.2cm for *E. coli*) and the Gene Pulser (Bio-Rad) at the settings 200 Ω, 25 µF, and 1.8 kV (at 2.2kVd). As needed, LB was supplemented with 10% Sucrose or antibiotics at the following concentrations: ampicillin (Amp): 100µg/mL; carbenicillin (Carb): 50-1000µg/mL; gentamicin (Gent): 100µg/mL; chloramphenicol (Cm): 100µg/mL; kanamycin (Kan): 20μg/mL; or CIP: varying concentrations (Sigma).

### Molecular cloning and mutant construction

Primers (IDT) and plasmids described in this study as well are listed in Table S1. The AdeS (R152S) mutant was isolated by drug selection of a precursor strain by plating on LB with 10 μg/mL gentamicin. Single colonies were purified on LB agar plates, and then *adeR* and *adeS* were amplified by colony PCR and sequenced (Genewiz) to identify mutations. The selected *adeS* mutation (R152S) was introduced into *A. baumannii gyrA(S81L)parC(S84L)* (also called GP (GVA37)) to create *gyrA(S81L)parC(S84L)adeS(R152S)* in a fashion similar to that described previously (also called GPS (EHA105)) (65). Briefly, the mutant allele was amplified with upstream and downstream flanking sequences to generate a product of about 1500bp in length and introduced first into pUC18 by restriction enzyme digestion and T4 DNA ligation. Once propagated in DH5α and sequence confirmed (Genewiz), the mutant allele was introduced into the pJB4648 plasmid by restriction enzyme digestion and T4 DNA ligation and propagated in DH5α λpir. The mutant allele in pJB4648 was then introduced into *A. baumannii* by electroporation and homologous recombination with two selection steps, the first selecting for transformants on LB+10 μg/mL gentamicin on bacteriological plates and the second counterselection for Gent^S^ on LB+10% Sucrose plates at either room temperature or at 30°C. The internal, in-frame gene deletion mutants were previously created or constructed during this study using the same methodology after three-way restriction enzyme digestions and ligations (40, 41, 65). The GPL (EHA23) strain (*gyrA(S81L)parC(S84L)adeL(ΔI335A336)*) was constructed using the gentamicin selection/sucrose counterselection. The *adeL(ΔI335A336)* allele was PCR amplified from gDNA isolated from P9 Cy3.2 (52) with flanking regions, cloned into pUC18 then moved into pJB4648, propagated in XL1 Blues and DH5α λpir *E. coli*, sequence confirmed, then introduced into the GP background by electroporation and homologous recombination. GP was created previously (17) and *gyrA(S81L)parC(S84L)adeN*::ISAba1 (GVA47), also known as GPN, was generated by plating GP on LB+32μg/mL CIP.

### WGS

Strains used in this study were sequence verified by whole genome sequencing (WGS) as previously described (52). Illumina Nextera reagents were used for library preparation and 100bp single-end reads were obtained using the Tufts University Core Facility’s HiSeq2500. BRESEQ 0.35.5 was used to align reads to the *Acinetobacter baumannii* 17978UN reference genome (GenBank Accession: CP079212.1), and mutations, those of which are absent in the parent strains, were identified. All sequences are deposited in the SRA database under BioProject number PRJNA1068501.

### LOS analysis

LOS profiles were examined using Tricine gel electrophoresis of cell lysates as previously described (40), using lysates from cells grown to mid-logarithmic phase. After fractionation on Novex 16% tricine gels (Invitrogen), the gels were stained with Pro-Q Emerald 300 (Invitrogen) for LOS staining, and Coomassie Brilliant Blue for total protein staining for sample quantification normalization (40).

### pHi assay

Buffers of varying pH were made using 50mL combinations of 0.2M Na_2_HPO_4_ and 0.2M NaH_2_PO_4_ (66). 5mM BCECF/AM (ThermoFisher) and 0.5mM carbonyl cyanide 3-chlorophenylhydrazone (CCCP; Sigma) were prepared in DMSO and stored at –20°C until use. Methods were adapted with modifications from previous protocols (67). Overnight cultures prepared from individual colonies were diluted for recovery to allow growth, then diluted again to A_600_ ∼0.003 in 8mL LB broth (with or without drug as indicated). In the case of dCas9 strains, single colonies were inoculated into 2mL LB in capped tubes and incubated to late exponential phase before diluting into LB + 100ng/mL anhydrotetracycline (aTc). The cultures that were exposed to sub-MIC levels of CIP were treated with 45 μg/mL of CIP an hour prior to harvest. Cultures were grown to exponential phase (A_600_=0.4-0.7) or early post exponential phase (A_600_=1.2-1.4) after which samples (about 5mL) were pelleted in 15mL conical tubes at 7500g for 5 min, resuspended in sterile phosphate-buffered saline (PBS, Thermo Scientific), loaded with 20µM BCECF/AM in PBS followed by incubation at 30°C for 30 min in a 96-well v-bottom plate protected from light.

Cells in the 96 well plate were diluted 1:4, pelleted at 2000rpm for 5 min, resuspended in LB broth and incubated at 37°C for 5 min, before pelleting once more and re-suspending in 200-400µl previously prepared mono– and di-basic sodium phosphate buffer at neutral pH (experimental samples) or at a range of pH values (*in situ* standard curve samples). Samples were diluted 1:1 in the neutral or pH adjusted buffers to a final volume of 200µl in a 96-well black plate with a clear, flat bottom (Costar). About 10µM CCCP stock was added to the *in situ* standard curve wells. All samples were mixed via pipetting before fluorescence was read at 490ex/535em and 440ex/535em after 5 minutes (Biotek SynergyMx with Gen5 software). Background fluorescence from buffer alone was subtracted from each sample before calculating the ratiometric fluorescence of the calibration curve samples and experimental samples. The experimental sample internal pH was interpolated from the *in-situ* calibration curve using standard curve fitting.

### Preparation of transposon mutant banks

Transposon banks with *Himar-1* transposon were created as described (17, 40). 50µl electrocompetent *A. baumannii* cells were electroporated with 300ng pDL1100 (*Himar-1-*Kan^R^). Cells were recovered in 1mL SOC broth, placed in 14mL snap cap tubes (Corning) and aerated on a drum roller for 15min at 37°C. The cells were pelleted and resuspended in 400µl SOC, then spread onto two 150 mm culture dishes containing mixed cellulose ester filters (0.45µm pore size, 137mm diameter; Advantec) that had been placed on top of the SOC agar. Plates were incubated for 1hr at 37°C before filters were transferred to 150 mm culture dishes containing LB+20µg/mL kanamycin and incubated overnight (∼16hrs) at 37°C.

Mini-*Tn10* banks were also made in the GPN strain background. GPN electrocompetent cells were transformed with 75ng pDL1073 (mTn*10*-*neo*), with subsequent steps following the same procedure as used for *Himar-1.* CFU counts were determined from plate images using an Image J counting routine. Mutants were pooled from filters by gentle shaking in 50-75mL sterile PBS. Glycerol was added for a final concentration of around 10% and aliquots stored at –80oC until use. About 21 independent subpools were created in each strain background (producing around 100,000-150,000 cumulative unique mutants).

### CIP challenge of transposon banks

Independent subpools comprising of approximately 7,500-16,000 unique transposon mutants were challenged with antibiotic as described (17, 40). Mutant banks were thawed on ice, diluted in 25mL LB broth to an A_600_=0.1, grown to A_600_=0.2 at 37°C, then diluted to A_600_=0.003 in 10mL LB broth with or without CIP of varied concentrations added in parallel cultures. The remaining culture (T_1_) was harvested to determine what mutants were present prior to outgrowth and drug challenge. Cultures were incubated for about 8 generations (around 170min if untreated, 225min if CIP treated), then harvested (T_2_).

Drug-treated cultures that exhibited 20-30% growth inhibition were harvested. Cultures were pelleted and stored at –20°C until all samples were collected. The concentration ranges at which comparable growth inhibition was met for each strain background as well as the average population expansions between T_1_ and T_2_ with each condition (according to A_600_) are noted in Data Set S2.

### Preparation of transposon libraries for sequencing

As described (17, 40), the gDNA was extracted using the DNeasy Blood and Tissue Kit (Qiagen). Samples were treated with proteinase K during resuspension and incubated at 56°C with shaking for 1hr before treatment with RNase cocktail (Fisher Scientific) for 8min at 62°C. The gDNA was quantified with SYBR green then diluted to 7ng/µl in UltraPure water (Fisher Scientific). About 28ng of gDNA of each sample was tagmented using TD buffer and TDE1 enzyme (Illumina) and incubated in a thermocycler at 55°C for 5min prior to melting at 95°C for 0.5min. Transposon insertion sites were then amplified using Q5 hot start polymerase (NEB) and 0.6µM Nextera 2A-R and olj638 (for *Himar-1* libraries), or olj928 (for Tn*10* libraries). The 50µl reactions were incubated at 98°C for 1min, cycled 30 times through 98°C for 10sec, 65°C for 20 sec, and 72°C for 1min, then finally at 72°C for 2min. 0.5µl of the first PCR products were used for a second PCR reaction used to introduce indexes with leftward *Himar-1-* or Tn10-specific indexed primers and rightward Nextera indexed primers. The reaction and settings were the same except cycling was shortened to 12 cycles. Amplified products were quantified via 2% agarose tris-acetate-EDTA (TAE) gels using SYBR Safe (Invitrogen) for visualization. The gels were loaded with a known amount of 100bp DNA ladder (NEB) alongside samples of each reaction to quantify their concentrations based on band intensity around 250-600bp using a gel imaging system. Equal measures of all samples were thereafter pooled and purified via PCR column clean-up (Qiagen). About 17.5 pmol of each pooled sample was reconditioned in an additional 50µl reaction with primers P1 and P2. Thermocycler settings were set with a heated lid and incubations at 95°C for 1min, with slow ramp of 0.1°C/sec to 64°C then held for 20sec, and finally 72°C for 10min. The products were purified once more with the PCR column clean-up kit (Qiagen). Primer olj115 (Tn*10*) or mar512 (*Himar-1*) was used for sequencing at the Tufts University Core Facility of Genomics with the Illumina HiSeq 2500 (SE50, post PippinPrep (Sage Science) size selection for 250-600bp).

### Sequence analysis and assessment of mutant fitness

Sequenced reads were processed and analyzed as done previously (17, 39, 40, 68). After demultiplexing, the Nextera adapter sequence was clipped from reads using the fastx_clipper, quality filtered, and reads shorter than 20nt were discarded. Reads were mapped to the *A. baumannii* ATCC 17978 reference genome (https://genomes.atcc.org/genomes/e1d18ea4273549a0) using Bowtie 1.2.2, formatted so only unique and best-mapped reads were aligned, allowing up to 1 maximum mismatch. Fitness scores were then calculated for each transposon mutant using MAGenTA (68), with scores averaged and aggregated at the gene level across replicates. Insertions within 10% of 3’ end of genes were excluded and fitness scores normalized to a set of neutral/pseudo genes. Of note, fitness scores of GPN insertion mutants were determined using data from both *Himar-1* and mini-*Tn10* banks. All other results were from *Himar-1* mutagenesis. Fitness scores of gene disruptions in all strain backgrounds were normalized to a LOWESS curve to avoid the position bias exhibited post CIP exposure. The difference in fitness scoring for each gene disruption was assessed between each FQR strain and the WT drug susceptible strain. Significant fitness differences yielded from the same gene disruption between the different strain backgrounds were identified via multiple t test analysis using the original FDR method of Benjamini and Hochberg with significance falling within the FDR of 5% (q<0.05).

### RNAseq experiment to uncover transcriptional differences between FQR and parent strains

Cultures were grown overnight, diluted to an A_600_ of 0.1 in 10mL LB broth, grown for about 1.5hrs at 37°C on a drum wheel to approximately mid-log growth, then diluted once more to A_600_ of 0.003 in fresh LB broth with or without CIP. Some of these were grown to approximately mid-log phase by incubation for 150-165min (untreated) or for 195-225min (CIP treated) as similarly done before (17) and reaching A_600_∼0.445-0.68 and A_600_∼0.39-0.66 respectively. Others were grown for 195-210min (untreated) reaching A_600_∼0.85-1.195 or for 255-270min (CIP treated) to A_600_∼0.82-1.125. Of the cultures treated with CIP, those that exhibited 20-30% growth inhibition (compared to the untreated culture grown in parallel) were harvested (details provided in Data Sets S3 and S4). About 1-1.5 A_600_ units of all cultures were harvested and frozen at –80°C until all samples were collected.

### Preparing RNA for sequencing

RNA was isolated from the samples using the RNeasy Mini kit (Qiagen) and prepared for sequencing as described (17, 69, 70) following published protocols (71). Approximately 600ng of RNA was fragmented at 94°C with 10x FastAP buffer for 2.5min then treated with TURBO DNase, FastAP (Thermo Scientific) (and murine RNase inhibitor; NEB) for 30min at 37°C. Samples were then purified using Agencourt RNAClean XP (Beckman Coulter) and eluted with 12.5µl of DEPC UltraPure water (Fisher Scientific). 5µl of sample was incubated with 1µl of 100µM barcoded RNA adapters at 72°C for 2 min before mixed with DMSO (American Bioanalytical), 10x T4 RNA ligase buffer, 100mM ATP, RNase inhibitor murine, T4 RNA ligase, and 50% PEG8000 (NEB) to allow for 3’ linker ligation at room temperature for 1.5 hours. The 60 samples were pooled into 5 groups of 12 samples each during purification using RNA Clean & Concentrator (Zymo Research) and all eluted with 16µl DEPC UltraPure water. Ribosomal rRNA was depleted using RNase-H as described (72). Probes complementary to *A. baumannii* ATCC 17978UN 23S, 16S, and 5S rRNA sequences (CP079931.1) were created using 0.design_probe.py and off-target binding predicted using 2.predict_probe_offtarget.py (https://github.com/hym0405/RNaseH_depletion). The 89 devised oligonucleotides (IDT) were resuspended into 100µM in UltraPure water, and a master-mix created with 5µl of each probe.

For each depletion, the RNA was mixed with 1µl of the probe mixture, 0.6µl 1M Tris-HCl pH 7.5, 1.5µl 5M NaCl, and DEPC UltraPure water for a final 15µl reaction. They were incubated in a thermocycler with a heated lid at 95°C for 2min, temperature reduced to 45°C in 0.1°C/sec increments and incubated a further 5 min at 45°C. Meanwhile, RNase-H master mix was prepared for each depletion (3µl Hybridase Thermostable RNase H; Epicentre), 0.5µl 1M Tris-HCl pH=7.5, 0.2µl 5M NaCl, 0.4µl 1M MgCl2, and 0.9µl DEPC UltraPure water) and preheated to 45°C. Post probe and RNA hybridization, 5µl prewarmed RNase-H mixture was added to each probe-RNA mixture and incubated at 45°C for 30min. These were then purified twice with 2x SPRI RNAClean XP (Agencourt) and eluted with 13µl DEPC UltraPure water.

The RNA preparation was then used as a template for cDNA synthesis. 12µl of sample was incubated with 2µl of 25µM AR2 primer at 72°C for 2min using 10x AffinityScript RT buffer and enzyme (Agilent Technologies), 0.1M DTT, 100mM dNTPs, and 40U/µl murine RNase inhibitor. After 55min at 55°C, RNA was then denatured in a 10% reaction volume (adding 2µl to the 20µl cDNA reaction product) of 1N freshly prepared NaOH at 70°C for 12min before neutralization with 4µl of freshly prepared 0.5M acetic acid. The products were then subjected to Agencourt RNAClean XP, eluted with 5µl DEPC UltraPure water, incubated with 2µl of 40µM 3Tr3 adaptor at 75°C for 3min with the beads still present, and mixed with 10x T4 ligase buffer, DMSO, ATP, PEG8000, and T4 RNA ligase for an overnight 20µl ligation reaction at room temperature. The products were purified with SPRI RNAClean XP (Agencourt) beads twice and finally eluted with 25µl H_2_0. DNA was amplified from cDNA using 10µl of 5x Q5 reaction buffer and 2.5µl Q5 enzyme (NEB), 0.5µl 100mM dNTPs, 10µl cDNA, 2.5µl primers RTS_Enr_P5 and RTS_Enr_P7_BCXX, and water for 50µl reaction. The reactions were incubated at 98°C for 3min, cycled at 98°C for 30 sec, 67°C for 25sec, and 72°C for 15sec for 14 cycles, and completed at 72°C for 2min. PCR products were purified once with AxyPrep Magnetic PCR beads (Axygen; Corning). DNA was then quantified by Qubit with the dsDNA HS Assay kit (RNA HS Assay kit was used previously for the quantification of the RNA samples; Thermo Scientific) and about 100nM of each of the 5 subpools was pooled together into 1 sample for sequencing.

Throughout sample preparation, 1µl SUPERase-IN RNase inhibitor (Thermo Scientific) was added to samples if they were stored at –80°C in between steps. RNAse Zap was used continuously on surfaces until cDNA was synthesized. Samples were routinely analyzed by submitting samples for QC fragment analysis at the TUCF Genomics Core. The pooled samples were sequenced at this facility, using a NextSeq550 Single-End High output, with 75 cycles following nanoQC.

### RNAseq analysis

Sequences that had been de-multiplexed for the i7 barcodes were then de-multiplexed for the i5 barcodes using fastx_barcode_splitter.pl (https://github.com/agordon/fastx_toolkit/). The 5’ anchored adaptor sequences were then trimmed *in silico* using Cutadapt (73). Truseq adaptor sequences were also trimmed *in silico*, disallowing any error, and dropping resulting sequences shorter than 20nt. Sequence data were thereafter analyzed following and modifying published scripts (https://github.com/huoww07/Bioinformatics-for-RNA-Seq). The Acinetobacter_baumannii_ATCC_17978.fasta and Acinetobacter_baumannii_ATCC_17978.gbk files were downloaded from the ATCC genome portal (https://genomes.atcc.org/genomes/e1d18ea4273549a0?tab=overview-tab) to create the STAR genome index (via genomeGenerate with STAR). Annotation GTF files were constructed using Geneious, then cufflinks gffread for conversion to a strict gtf file. Alignments were then performed using STAR. Subread/1.6.3 was used to get a summary of the sequence data features, then featurecount_stat.R as well as mapping_percentage.R were adapted to analyze and visualize features including mapping efficiency and overview of mapped vs. unmapped reads. Using the Tufts Cluster on-demand R studio app, DESeq2, ggplot2, dplyr, tidyverse, ggrepel, DEGreport, and pheatmap tools were employed to analyze the quality of the dataset as well as visualize and deconvolve the contrasting transcriptomic data between the high-level FQR pump-overproducing strains and the GP parental strain.

### qPCR analysis for transcript abundance

Cultures derived from individual colonies were grown in LB broth to mid-or early post-exponential phase. About 1 A_600_ unit of culture was harvested and RNA was extracted using RNeasy (Qiagen). cDNA was synthesized after ezDNase treatment using the SuperScript VILO cDNA kit (Thermo Fisher), then amplified using PowerUp SYBR Green Master Mix (Applied Biosystems) and StepOnePlus Real-Time PCR system (Applied Biosystems) per manufacturer’s instructions. The efficiency of target amplification was tested using a standard curve of cDNA at varying dilutions and consequently verified to be greater than 95% for each primer pair used. Three biological replicates were evaluated for each strain and three technical replicates examined per biological sample. To confirm a lack of gDNA contamination, control samples tested in parallel which lacked reverse-transcriptase during cDNA synthesis were included. Transcript levels for each indicated target were compared between each mutant strain and the parental strain (via the comparative 2^-ΔΔCT^ method) after normalization to that of 16S, the endogenous control. The data are represented in bar plots showing the mean fold difference in transcript levels (2^-ΔΔCT^) ± SEM.

### MIC determination

MICs were determined by broth microdilutions. Overnight cultures were diluted to A_600_=0.1 in 3mL LB broth, incubated to early exponential phase (A_600_=0.2-0.3), then diluted to a final A_600_=0.003 in 200ul LB broth in the presence of different dilutions of CIP in a 96-well plate (Costar). Growth was monitored by measuring A_600_ every 15min (Epoch 2 or SynergyMx, (Biotek) plate readers). At least three replicates were tested for all strains. CIP MIC was determined as the lowest concentration that prevented growth up to 12 hours, where geometric mean A_600_ units did not exceed 0.035, and are not significantly different from the LB broth-only negative controls. CIP MICs were 256µg/mL (GPS and GPN) and 384µg/mL (GPL).

### Persistence assay

The *A. baumannii* strains EHA417 (*adeL*ΔI335A336 *P_Tet_::dCas9* (pWH1266-*paaA*-sgRNA)), EHA418 (*adeL*ΔI335A336 *P_Tet_::dCas9* (pWH1266-NC-sgRNA)), and EHA254 (WT *P_Tet_::dCas9* (pWH1266-NC-sgRNA)) were incubated in 1.5mL LB in triplicate for 2-3 hours. They were then diluted 1/1000 in 8mL LB + 100ng/mL anhydrotetracycline (aTc) to induce dCas9 expression and grown to mid-exponential phase. Each culture was then normalized to A_600_∼0.5 with fresh LB containing inducer aTc, then treated with 10 μg/mL CIP. Samples of culture (500µl) were removed at timepoints=0, 1, 6, and 24 hours, washed with PBS, then resuspended in PBS and serial dilutions were spotted on LB agar plates for overnight incubation at 37°C and CFU enumeration the next day.

### Microscopy

Culture samples were placed on 1% agarose in PBS pads and imaged using DIC at 100X magnification on the Zeiss Axiovert Observer.Z1.

### Accession numbers

All analyzed sequences are in the SRA database under BioProject numbers PRJNA1067619 (TnSeq), PRJNA1067535 (RNAseq), and PRJNA1068501 (WGS).

## Acknowledgements

This work is supported by NIAID awards U01AI124302 and U19AI158076 (RRI, TvO), U19AI142780 and R21AI128328 (RRI), and R01AI162996 (EG). We thank David Lazinski (Tn plasmids), Germán Vargas-Cuebas (for the GPN mutant), Rebecca Batorsky (bioinformatic help with TRANSIT), Yunfei Dai (for communications regarding dCas9 strain generation, sgRNA design, and TRANSIT, and for the pUC-Tn7-GentR-pTet::dCas9 and pWH1266-sgRNA plasmids), Matthew Hall (for help with RNAseq library preparation protocols), Albert Tai and Irina V. Grinvald for communications regarding sequencing and library preparation, as well as the next-generation sequencing support provided by S10OD032203 via Tufts University Core Facility Genomics Core.

## Author Contributions

R.R.I and E.G. designed the TnSeq study and provided training and support with experimental methodologies. S.S. and A.C. created mariner banks in the WT and GPN strains. A.C. performed TnSeq experiments with GPN and WT. J.B. performed the LOS experiments and analyses. W.H. provided bioinformatic support and ran the DESeq2 bioinformatic workflow for the comparative RNAseq study. J.H-B. performed the WGS library prep and sequence analysis for all deletion mutants. J.C.O-M. and TvO provided support with RNAseq library preparation methodologies. E.G. created the vectors used to make two deletion mutants and the backbone vector used for gene complementation. E.H. and R.R.I devised the research, interpreted the results, and wrote the manuscript. E.G reviewed and revised the manuscript.

## References

1. Peleg AY, Seifert H, Paterson DL. 2008. *Acinetobacter baumannii*: emergence of a successful pathogen. Clin Microbiol Rev 21:538–82.

2. CDC. 2019. Antibiotic Resistance Threats in the United States. Atlanta, GA: US Department of Health and Human Services, CDC.

3. CDC. 2022. COVID-19: U.S. Impact on Antimicrobial Resistance, Special Report 2022. Atlanta, GA: US Department of Health and Human Services, CDC.

4. Tomaras AP, Dorsey CW, Edelmann RE, Actis LA. 2003. Attachment to and biofilm formation on abiotic surfaces by *Acinetobacter baumannii*: involvement of a novel chaperone-usher pili assembly system. Microbiology (Reading) 149:3473–3484.

5. Badave GK, Kulkarni D. 2015. Biofilm Producing Multidrug Resistant *Acinetobacter baumannii*: An Emerging Challenge. J Clin Diagn Res 9:DC08–10.

6. Sanchez CJ, Jr., Mende K, Beckius ML, Akers KS, Romano DR, Wenke JC, Murray CK. 2013. Biofilm formation by clinical isolates and the implications in chronic infections. BMC Infect Dis 13:47.

7. Gonzalez-Villoria AM, Valverde-Garduno V. 2016. Antibiotic-Resistant *Acinetobacter baumannii* Increasing Success Remains a Challenge as a Nosocomial Pathogen. J Pathog 2016:7318075.

8. Tacconelli E, Carrara E, Savoldi A, Harbarth S, Mendelson M, Monnet DL, Pulcini C, Kahlmeter G, Kluytmans J, Carmeli Y, Ouellette M, Outterson K, Patel J, Cavaleri M, Cox EM, Houchens CR, Grayson ML, Hansen P, Singh N, Theuretzbacher U, Magrini N, Group WHOPPLW. 2018. Discovery, research, and development of new antibiotics: the WHO priority list of antibiotic-resistant bacteria and tuberculosis. Lancet Infect Dis 18:318–327.

9. Lob SH, Hoban DJ, Sahm DF, Badal RE. 2016. Regional differences and trends in antimicrobial susceptibility of *Acinetobacter baumannii*. Int J Antimicrob Agents 47:317–23.

10. Giammanco A, Cala C, Fasciana T, Dowzicky MJ. 2017. Global Assessment of the Activity of Tigecycline against Multidrug-Resistant Gram-Negative Pathogens between 2004 and 2014 as Part of the Tigecycline Evaluation and Surveillance Trial. mSphere 2.

11. Poole K. 2004. Efflux-mediated multiresistance in Gram-negative bacteria. Clin Microbiol Infect 10:12–26.

12. Sun J, Deng Z, Yan A. 2014. Bacterial multidrug efflux pumps: mechanisms, physiology and pharmacological exploitations. Biochem Biophys Res Commun 453:254–67.

13. Coyne S, Courvalin P, Perichon B. 2011. Efflux-mediated antibiotic resistance in *Acinetobacter* spp. Antimicrob Agents Chemother 55:947–53.

14. Blair JM, Webber MA, Baylay AJ, Ogbolu DO, Piddock LJ. 2015. Molecular mechanisms of antibiotic resistance. Nat Rev Microbiol 13:42–51.

15. Geisinger E, Huo W, Hernandez-Bird J, Isberg RR. 2019. *Acinetobacter baumannii*: Envelope Determinants That Control Drug Resistance, Virulence, and Surface Variability. Annu Rev Microbiol 73:481–506.

16. Compagne N, Vieira Da Cruz A, Muller RT, Hartkoorn RC, Flipo M, Pos KM. 2023. Update on the Discovery of Efflux Pump Inhibitors against Critical Priority Gram-Negative Bacteria. Antibiotics (Basel) 12.

17. Geisinger E, Vargas-Cuebas G, Mortman NJ, Syal S, Dai Y, Wainwright EL, Lazinski D, Wood S, Zhu Z, Anthony J, van Opijnen T, Isberg RR. 2019. The Landscape of Phenotypic and Transcriptional Responses to Ciprofloxacin in *Acinetobacter baumannii*: Acquired Resistance Alleles Modulate Drug-Induced SOS Response and Prophage Replication. mBio 10.

18. Naidu V, Bartczak A, Brzoska AJ, Lewis P, Eijkelkamp BA, Paulsen IT, Elbourne LDH, Hassan KA. 2023. Evolution of RND efflux pumps in the development of a successful pathogen. Drug Resist Updat 66:100911.

19. Pagdepanichkit S, Tribuddharat C, Chuanchuen R. 2016. Distribution and expression of the Ade multidrug efflux systems in *Acinetobacter baumannii* clinical isolates. Can J Microbiol 62:794–801.

20. Magnet S, Courvalin P, Lambert T. 2001. Resistance-nodulation-cell division-type efflux pump involved in aminoglycoside resistance in *Acinetobacter baumannii* strain BM4454. Antimicrob Agents Chemother 45:3375–80.

21. Coyne S, Rosenfeld N, Lambert T, Courvalin P, Perichon B. 2010. Overexpression of resistance-nodulation-cell division pump AdeFGH confers multidrug resistance in *Acinetobacter baumannii*. Antimicrob Agents Chemother 54:4389–93.

22. Yoon EJ, Chabane YN, Goussard S, Snesrud E, Courvalin P, De E, Grillot-Courvalin C. 2015. Contribution of resistance-nodulation-cell division efflux systems to antibiotic resistance and biofilm formation in *Acinetobacter baumannii*. mBio 6.

23. Rosenfeld N, Bouchier C, Courvalin P, Perichon B. 2012. Expression of the resistance-nodulation-cell division pump AdeIJK in *Acinetobacter baumannii* is regulated by AdeN, a TetR-type regulator. Antimicrob Agents Chemother 56:2504–10.

24. Wang Z, Fan G, Hryc CF, Blaza JN, Serysheva, II, Schmid MF, Chiu W, Luisi BF, Du D. 2017. An allosteric transport mechanism for the AcrAB-TolC multidrug efflux pump. Elife 6.

25. Murakami S, Nakashima R, Yamashita E, Yamaguchi A. 2002. Crystal structure of bacterial multidrug efflux transporter AcrB. Nature 419:587–93.

26. Murakami S, Nakashima R, Yamashita E, Matsumoto T, Yamaguchi A. 2006. Crystal structures of a multidrug transporter reveal a functionally rotating mechanism. Nature 443:173–9.

27. Koronakis V, Sharff A, Koronakis E, Luisi B, Hughes C. 2000. Crystal structure of the bacterial membrane protein TolC central to multidrug efflux and protein export. Nature 405:914–9.

28. Yu EW, Aires JR, McDermott G, Nikaido H. 2005. A periplasmic drug-binding site of the AcrB multidrug efflux pump: a crystallographic and site-directed mutagenesis study. J Bacteriol 187:6804–15.

29. Yu EW, McDermott G, Zgurskaya HI, Nikaido H, Koshland DE, Jr. 2003. Structural basis of multiple drug-binding capacity of the AcrB multidrug efflux pump. Science 300:976–80.

30. Higgins MK, Bokma E, Koronakis E, Hughes C, Koronakis V. 2004. Structure of the periplasmic component of a bacterial drug efflux pump. Proc Natl Acad Sci U S A 101:9994–9.

31. Sugawara E, Nikaido H. 2014. Properties of AdeABC and AdeIJK efflux systems of *Acinetobacter baumannii* compared with those of the AcrAB-TolC system of *Escherichia coli*. Antimicrob Agents Chemother 58:7250–7.

32. Leus IV, Weeks JW, Bonifay V, Smith L, Richardson S, Zgurskaya HI. 2018. Substrate Specificities and Efflux Efficiencies of RND Efflux Pumps of *Acinetobacter baumannii*. J Bacteriol 200.

33. DeJesus MA, Ambadipudi C, Baker R, Sassetti C, Ioerger TR. 2015. TRANSIT--A Software Tool for Himar1 TnSeq Analysis. PLoS Comput Biol 11:e1004401.

34. Bai J, Dai Y, Farinha A, Tang AY, Syal S, Vargas-Cuebas G, van Opijnen T, Isberg RR, Geisinger E. 2021. Essential Gene Analysis in *Acinetobacter baumannii* by High-Density Transposon Mutagenesis and CRISPR Interference. J Bacteriol 203:e0056520.

35. Hamami E, Huo W, Neal K, Neisewander I, Geisinger E, Isberg RR. 2024. Identification of essential genes that support fitness of Acinetobacter baumannii efflux pump overproducers in the presence of fluoroquinolone. bioRxiv doi:10.1101/2024.01.04.574119:2024.01.04.574119.

36. VanOtterloo LM, Macias LA, Powers MJ, Brodbelt JS, Trent MS. 2024. Characterization of *Acinetobacter baumannii* core oligosaccharide synthesis reveals novel aspects of lipooligosaccharide assembly. mBio 15:e0301323.

37. Arai H, Kawakami T, Osamura T, Hirai T, Sakai Y, Ishii M. 2014. Enzymatic characterization and in vivo function of five terminal oxidases in *Pseudomonas aeruginosa*. J Bacteriol 196:4206–15.

38. Borisov VB, Verkhovsky MI. 2015. Oxygen as Acceptor. EcoSal Plus 6.

39. van Opijnen T, Bodi KL, Camilli A. 2009. Tn-seq: high-throughput parallel sequencing for fitness and genetic interaction studies in microorganisms. Nat Methods 6:767–72.

40. Geisinger E, Mortman NJ, Dai Y, Cokol M, Syal S, Farinha A, Fisher DG, Tang AY, Lazinski DW, Wood S, Anthony J, van Opijnen T, Isberg RR. 2020. Antibiotic susceptibility signatures identify potential antimicrobial targets in the *Acinetobacter baumannii* cell envelope. Nat Commun 11:4522.

41. Geisinger E, Mortman NJ, Vargas-Cuebas G, Tai AK, Isberg RR. 2018. A global regulatory system links virulence and antibiotic resistance to envelope homeostasis in *Acinetobacter baumannii*. PLoS Pathog 14:e1007030.

42. Russo TA, MacDonald U, Beanan JM, Olson R, MacDonald IJ, Sauberan SL, Luke NR, Schultz LW, Umland TC. 2009. Penicillin-binding protein 7/8 contributes to the survival of *Acinetobacter baumannii* in vitro and in vivo. J Infect Dis 199:513–21.

43. Bulssico J, Papukashvil II, Espinosa L, Gandon S, Ansaldi M. 2023. Phage-antibiotic synergy: Cell filamentation is a key driver of successful phage predation. PLoS Pathog 19:e1011602.

44. Bos J, Zhang Q, Vyawahare S, Rogers E, Rosenberg SM, Austin RH. 2015. Emergence of antibiotic resistance from multinucleated bacterial filaments. Proc Natl Acad Sci U S A 112:178–83.

45. Nikaido H, Takatsuka Y. 2009. Mechanisms of RND multidrug efflux pumps. Biochim Biophys Acta 1794:769–81.

46. Teufel R, Mascaraque V, Ismail W, Voss M, Perera J, Eisenreich W, Haehnel W, Fuchs G. 2010. Bacterial phenylalanine and phenylacetate catabolic pathway revealed. Proc Natl Acad Sci U S A 107:14390–5.

47. Hooppaw AJ, McGuffey JC, Di Venanzio G, Ortiz-Marquez JC, Weber BS, Lightly TJ, van Opijnen T, Scott NE, Cardona ST, Feldman MF. 2022. The Phenylacetic Acid Catabolic Pathway Regulates Antibiotic and Oxidative Stress Responses in *Acinetobacter*. mBio 13:e0186321.

48. Green ER, Juttukonda LJ, Skaar EP. 2020. The Manganese-Responsive Transcriptional Regulator MumR Protects *Acinetobacter baumannii* from Oxidative Stress. Infect Immun 88.

49. Vega NM, Allison KR, Khalil AS, Collins JJ. 2012. Signaling-mediated bacterial persister formation. Nat Chem Biol 8:431–3.

50. Wu Y, Vulic M, Keren I, Lewis K. 2012. Role of oxidative stress in persister tolerance. Antimicrob Agents Chemother 56:4922–6.

51. Lee HH, Molla MN, Cantor CR, Collins JJ. 2010. Bacterial charity work leads to population-wide resistance. Nature 467:82–5.

52. Huo W, Busch LM, Hernandez-Bird J, Hamami E, Marshall CW, Geisinger E, Cooper VS, van Opijnen T, Rosch JW, Isberg RR. 2022. Immunosuppression broadens evolutionary pathways to drug resistance and treatment failure during *Acinetobacter baumannii* pneumonia in mice. Nat Microbiol 7:796–809.

53. Park JS, Lee WC, Yeo KJ, Ryu KS, Kumarasiri M, Hesek D, Lee M, Mobashery S, Song JH, Kim SI, Lee JC, Cheong C, Jeon YH, Kim HY. 2012. Mechanism of anchoring of OmpA protein to the cell wall peptidoglycan of the gram-negative bacterial outer membrane. FASEB J 26:219–28.

54. Gribun A, Nitzan Y, Pechatnikov I, Hershkovits G, Katcoff DJ. 2003. Molecular and structural characterization of the HMP-AB gene encoding a pore-forming protein from a clinical isolate of *Acinetobacter baumannii*. Curr Microbiol 47:434–43.

55. Sugawara E, Nikaido H. 2012. OmpA is the principal nonspecific slow porin of *Acinetobacter baumannii*. J Bacteriol 194:4089–96.

56. Iyer R, Moussa SH, Durand-Reville TF, Tommasi R, Miller A. 2018. *Acinetobacter baumannii* OmpA Is a Selective Antibiotic Permeant Porin. ACS Infect Dis 4:373–381.

57. Powers MJ, Trent MS. 2018. Phospholipid retention in the absence of asymmetry strengthens the outer membrane permeability barrier to last-resort antibiotics. Proc Natl Acad Sci U S A 115:E8518–E8527.

58. Gwin CM, Prakash N, Christian Belisario J, Haider L, Rosen ML, Martinez LR, Rigel NW. 2018. The apolipoprotein N-acyl transferase Lnt is dispensable for growth in *Acinetobacter* species. Microbiology (Reading) 164:1547–1556.

59. Haas TM, Laventie BJ, Lagies S, Harter C, Prucker I, Ritz D, Saleem-Batcha R, Qiu D, Huttel W, Andexer J, Kammerer B, Jenal U, Jessen HJ. 2022. Photoaffinity Capture Compounds to Profile the Magic Spot Nucleotide Interactomes. Angew Chem Int Ed Engl 61:e202201731.

60. Diez S, Ryu J, Caban K, Gonzalez RL, Jr., Dworkin J. 2020. The alarmones (p)ppGpp directly regulate translation initiation during entry into quiescence. Proc Natl Acad Sci U S A 117:15565–15572.

61. Levenson-Palmer R, Luciano DJ, Vasilyev N, Nuthanakanti A, Serganov A, Belasco JG. 2022. A distinct RNA recognition mechanism governs Np(4) decapping by RppH. Proc Natl Acad Sci U S A 119.

62. Law RJ, Hamlin JN, Sivro A, McCorrister SJ, Cardama GA, Cardona ST. 2008. A functional phenylacetic acid catabolic pathway is required for full pathogenicity of *Burkholderia cenocepacia* in the *Caenorhabditis elegans* host model. J Bacteriol 190:7209–18.

63. Musthafa KS, Sivamaruthi BS, Pandian SK, Ravi AV. 2012. Quorum sensing inhibition in *Pseudomonas aeruginosa* PAO1 by antagonistic compound phenylacetic acid. Curr Microbiol 65:475–80.

64. Lightly TJ, Frejuk KL, Groleau MC, Chiarelli LR, Ras C, Buroni S, Deziel E, Sorensen JL, Cardona ST. 2019. Phenylacetyl Coenzyme A, Not Phenylacetic Acid, Attenuates CepIR-Regulated Virulence in *Burkholderia cenocepacia*. Appl Environ Microbiol 85.

65. Geisinger E, Isberg RR. 2015. Antibiotic modulation of capsular exopolysaccharide and virulence in *Acinetobacter baumannii*. PLoS Pathog 11:e1004691.

66. Gomori G. 1955. Preparation of buffers for use in enzyme studies. Methods in Enzymology 1:143.

67. Clementi EA, Marks LR, Roche-Hakansson H, Hakansson AP. 2014. Monitoring changes in membrane polarity, membrane integrity, and intracellular ion concentrations in *Streptococcus pneumoniae* using fluorescent dyes. J Vis Exp doi:10.3791/51008:e51008.

68. McCoy KM, Antonio ML, van Opijnen T. 2017. MAGenTA: a Galaxy implemented tool for complete Tn-Seq analysis and data visualization. Bioinformatics 33:2781–2783.

69. Jensen PA, Zhu Z, van Opijnen T. 2017. Antibiotics Disrupt Coordination between Transcriptional and Phenotypic Stress Responses in Pathogenic Bacteria. Cell Rep 20:1705–1716.

70. Zhu Z, Surujon D, Ortiz-Marquez JC, Huo W, Isberg RR, Bento J, van Opijnen T. 2020. Entropy of a bacterial stress response is a generalizable predictor for fitness and antibiotic sensitivity. Nat Commun 11:4365.

71. Shishkin AA, Giannoukos G, Kucukural A, Ciulla D, Busby M, Surka C, Chen J, Bhattacharyya RP, Rudy RF, Patel MM, Novod N, Hung DT, Gnirke A, Garber M, Guttman M, Livny J. 2015. Simultaneous generation of many RNA-seq libraries in a single reaction. Nat Methods 12:323–5.

72. Huang Y, Sheth RU, Kaufman A, Wang HH. 2020. Scalable and cost-effective ribonuclease-based rRNA depletion for transcriptomics. Nucleic Acids Res 48:e20.

73. Martin M. 2011. Cutadapt removes adapter sequences from high-throughput sequencing reads. 2011 17:3.

74. Benjamini Y, & Hochberg, Y. 1995. Controlling the false discovery rate: a practical and powerful approach to multiple testing. Journal of the Royal Statistical Society Series B (Methodological):289–300.

75. Benjamini Y, Krieger, A. M., & Yekutieli, D. 2006. Adaptive linear step-up procedures that control the false discovery rate. Biometrika 93:491–507.

