## Supplementary Figure Legends for "Identification of Determinants that Allow Maintenance of High-Level Fluoroquinolone Resistance in *Acinetobacter baumannii*"

**Supplementary Data Figure Legends**

**Supplemental Figure 1:** CyoA-targeted KD causes a fitness defect. (A) Volcano plot in which statistical significance is plotted as a function of the differences in insertion abundance between GPL and WT based on TRANSIT algorithm (33). All adjusted p-values = 0 were assigned the value of 0.005 to allow graphing. Blue: genes with adjusted p-values<0.05. Values for all other genes are shown in grey. (B,E) Number of reads plotted as a function of insertion site for noted genes as well as flanking regions. (C) Indicated strains grown in LB broth + 100ng/mL aTc. A_600_ for culture density are displayed and their mean ± SEM (n=4-5). *cyoA*-KD: strain harboring pWH1266 plasmid with *cyoA*-sgRNA vs NC-sgRNA is with nontargeting sgRNA. (D) Culture samples harvested in exponential phase and transcript levels determined by q-rtPCR (Materials and Methods). Displayed is mean fold change in transcript levels (log2 scale) ± SEM of each target compared to NC-sgRNA strain (n=3). KD: knockdown. aTc: anhydrotetracycline. WT: wildtype. GPL: *gyrAparCadeL*.

**Supplemental Figure 2**: Shown are scatter plots displaying the mean fitness of each gene disruption across the genome in GPL (A), WT (B), GPS (C), and GPN (D) strain backgrounds post CIP exposure. First panel shows the original mean fitness values. Red line at the neutral fitness value of 1 used to highlight the curvature of fitness scoring across the genome suggesting a positional bias. The red line in the middle panel shows the LOWESS curve used to normalize fitness scores and smooth the fitness landscape as seen in the last panel. WT: wildtype. GPL: *gyrAparCadeL*. GPN: *gyrAparCadeN*. GPS: *gyrAparCadeS*.

**Supplemental Figure 3:** OmpA, BfmS, LpsB, and PbpG are critical fitness determinants for *adeL*∆I335A336 strains during CIP exposure. Null mutants in FQR strain backgrounds compared to the GP parent strain background. Data for the GP strain and deletion mutants in the GP background from identical cultures are shown in each graph for comparative purposes. Growth was monitored every 15 minutes in 96 well plates. Displayed are means ± SEM (n≥3). To note, WGS data revealed the GP∆*bfmS::aacC1* strain has a single nucleotide insertion 97 bp upstream of *bfmR*. Ciptx: CIP treated. GP: *gyrAparC*. GPL: *gyrAparCadeL*. GPN: *gyrAparCadeN*.

**Supplemental Figure 4:** Loss of OmpA results in a fitness defect in the pump hyperexpresser which is linked to the presence of the efflux pump. Culture density was monitored every 15min overnight via plate reader. At least three biological replicates of each strain were tested. (A, E, G) All were grown in LB with CIP concentrations yielding comparable growth inhibition of the parent strains: 12.5µg/mL (GPL∆*adeFGH* and GP) or 65µg/mL (GPL). The first 8 hours of growth are shown, displayed as the mean ± SEM. (B-D) Additional clones of the GPL∆*ompA* genotype were tested to confirm OmpA’s role in maintaining fitness of the GPL mutant. (F, H) Both the GPL∆*ompA* and GPL∆*adeFGH*∆*ompA* clones contain a silent mutation in a SGNH/GDSL hydrolase family protein NDEONHPJ_01675 (ACX60_RS08320). GP: *gyrAparC*. GPL: *gyrAparCadeL*.

**Supplemental Figure 5:** The BfmS mutation causes lowered fitness in GPL, an effect that is linked to the presence of the AdeFGH efflux pump. Culture density was monitored every 15min overnight via plate reader. At least three biological replicates of each strain were tested. All were grown in LB with CIP concentrations yielding comparable growth inhibition of the parent strains: 12.5µg/mL (GPL∆*adeFGH* and GP) or 65µg/mL (GPL). The first 8 hours of growth are shown, displayed as the mean ± SEM. (B-E) Additional clones of GP∆*bfmS::aacC1* were tested to confirm minimal to no impact of the mutation on the growth of GP. (F, G) Growth of GPL∆*adeFGH* and GPL∆*adeFGH*∆*bfmS::aacC1* strains were evaluated against GPL∆*bfmS::aacC1* and GP∆*bfmS::aacC1.* (H) RNA was isolated from the noted strains (catalog numbers, Table S1) and relative transcript levels of BfmR (compared to the parental strains) were determined by q-rtPCR (Materials and Methods). Displayed is the mean fold change ± SEM (N=3). 304: ∆*bfmS::aacC1* (evaluated in Fig. 4B). 272: GP∆*bfmS::aacC1* (shown in Fig. S3A). 271: GP∆*bfmS::aacC1* (alternate clone displayed in B, & G). GP: *gyrAparC*. GPL: *gyrAparCadeL*.

**Supplemental Figure 6:** Poorer growth of the GPL strain in the absence of LpsB during CIP exposure is dependent on the expression of the AdeFGH efflux pump in GPL. Culture density was monitored every 15min overnight via plate reader, the first 8 hours shown here displayed as the mean ± SEM of at least three biological replicates for each strain. All were grown in LB with CIP concentrations yielding comparable growth inhibition of the parent strains: 12.5µg/mL (GPL∆*adeFGH* and GP) or 65µg/mL (GPL). Growth is compared between LpsB deletion mutants in the GPL∆*adeFGH* vs GPL strain backgrounds (B) and in the GPL∆*adeFGH* background assessed parallel to that in the GP strain background (C). The GPL∆*adeFGH*∆*lpsB* clone contains an additional mutation, N10D, in a hypothetical protein NDEONHPJ_02492 (ACX60_RS12045). GP: *gyrAparC*. GPL: *gyrAparCadeL*.

**Supplemental Figure 7:** Poorer growth of the GPL strain in the absence of PbpG during CIP exposure is likely attributable to the expression of the AdeFGH efflux pump in GPL. Culture density was monitored every 15min overnight via plate reader, the first 8 hours shown here displayed as the mean ± SEM of at least three biological replicates for each strain. All were grown in LB with CIP concentrations yielding comparable growth inhibition of the parent strains: 12.5µg/mL (GPL∆*adeFGH* and GP) or 65µg/mL (GPL). Growth is compared between PbpG deletion mutants in the GPL∆*adeFGH* vs GPL strain backgrounds (B) and GPL∆*adeFGH* vs GP strain backgrounds (C). GP: *gyrAparC*. GPL: *gyrAparCadeL*.

**Supplemental Figure 8:** Any impact on the LOS profile by the absence of PbpG or LpsB is independent of the strain background tested (A) Extracts of noted strains were fractionated on Novex 16% tricine gel and stained with Pro-Q Emerald 300 to reveal LOS. Representative images of three independent experiments are shown (left). Quantification of gel fractionated LOS, normalized to total protein (right). Shown are means relative to WT ± SD (n=3), individual LOS fractions in FQR strains vs in GP are compared using two-way ANOVA with Dunnett's multiple comparisons (right). (A) Parent strains. (B) *pbpG* deletion mutants. (C) *lpsB* deletion mutants. Significant differences are marked with an asterisk, *=p-value of 0.0498. F: Full length LOS. I: Intermediate. U: Unspecified LMM (low molecular mass) species. WT: wildtype. GPL: *gyrAparCadeL*. GPN: *gyrAparCadeN*. GPS: *gyrAparCadeS*. GP: *gyrAparC*.

**Supplemental Figure 9:** LpsB, PbpG, and BfmS mutants display morphotypes distinct from the parent strain. Images taken of samples from late log phase, grown in the absence of drug, using DIC at 100X. Scale bar added manually (10µm). Subscripts refer to different clones harboring the same mutation. GPL: *gyrAparCadeL*.

**Supplemental Figure 10:** LpsB, PbpG, and BfmS deletion mutants exhibit disparate structural responses to CIP exposure. Samples were taken during exponential growth with 65µg/mL CIP and imaged with DIC at 100X. Scale bar added manually (10µm). Subscripts refer to different clones harboring the same mutation. GPL: *gyrAparCadeL*.

**Supplemental Figure 11:** Structural changes in PbpG mutants during CIP exposure. Samples were taken during growth in LB media supplemented with 15µg/mL CIP (GP strain background) or 65µg/mL CIP (GPL strain background) and imaged with DIC at 100X after 2 and 4 hours of treatment. Scale bar added manually (10µm). GP: *gyrAparC*. GPL: *gyrAparCadeL*.

**Supplemental Figure 12:** Morphology of OmpA deletion mutants during CIP exposure. Samples were taken during growth in LB media supplemented with 15µg/mL CIP (GP strain background) or 65µg/mL CIP (GPL strain background) and imaged with DIC at 100X after 2 and 4 hours of treatment. Scale bar added manually (10µm). Subscripts refer to different clones harboring the same mutation. GP: *gyrAparC*. GPL: *gyrAparCadeL*.

**Supplemental Figure 13:** Morphological alterations due to lesions in BfmS during CIP pressure. Representative images of samples taken after 2 and 4 hours of growth in LB media supplemented with 15µg/mL CIP (GP strain background) or 65µg/mL CIP (GPL strain background), imaged using DIC at 100X magnification. Scale bar added manually (10µm). Subscripts refer to different clones harboring the same mutation. GP: *gyrAparC*. GPL: *gyrAparCadeL*.

**Supplemental Figure 14:** Morphotype exhibited by LpsB mutants during CIP treatment. Images were taken after 2 and 4 hours of growth in LB media with 15µg/mL CIP (GP strain background) or 65µg/mL CIP (GPL strain background), imaged using DIC at 100X magnification. Scale bar added manually (10µm). GP: *gyrAparC*. GPL: *gyrAparCadeL*.

**Supplemental Figure 15:** The pump overexpressing strains exhibit similar morphology as the parental strain with lower CIP resistance and the WT drug susceptible strain during exponential growth in the absence of drug. Representative images taken with DIC at 100X. Scale bar added manually (10µm). WT: wildtype. GP: *gyrAparC*. GPL: *gyrAparCadeL*. GPS: *gyrAparCadeS.* GPN: *gyrAparCadeN*.

**Supplemental Figure 16:** CIP exposure causes cell elongation that is comparable across the pump overexpressing and parental strains during logarithmic growth. Images taken with DIC at 100X. Scale bar added manually (10µm). CIP (µg/mL): 0.1 (WT), 15 (GP), 55 (GPN), & 65 (GPS & GPL). WT: wildtype. GP: *gyrAparC*. GPL: *gyrAparCadeL*. GPS: *gyrAparCadeS.* GPN: *gyrAparCadeN*.

**Supplemental Figure 17:** Fitness of transposon insertion mutants in GPL vs WT. Heatmaps of the normalized fitness of each transposon gene knockout in GPL and WT grown in the presence of CIP (A) or in the absence of drug pressure (B) alongside the difference in fitness scoring between the two strain backgrounds. Scatter plots show the fitness scores for each gene transposon knockout across the genome post CIP treatment (C) or without drug (D) for the drug susceptible WT strain alongside GPL. The noted genes are those that cause significant fitness changes in GPL compared to WT (where fitness differences ≥10%), with the exclusion of genes characterized as essential and genes associated with prophages or plasmids. Gene names are followed by accession numbers which are preceded by “NDEONHPJ_” per the ATCC 17978 strain annotations. Genes highlighted in teal are transposon disruptions that yield higher fitness scores in GPL compared to WT, with dark grey being those causing lower fitness in GPL (vs WT). CIP treatment used concentrations yielding 20-30% growth inhibition in each parent strain compared to the untreated grown in parallel. WT: wildtype. GPL: *gyrAparCadeL*.

**Supplemental Figure 18:** Fitness of transposon insertion mutants in GPN vs WT. Heatmaps of the normalized fitness of each transposon gene knockout in GPN and WT grown in the presence of CIP (A) or in the absence of drug pressure (B) alongside the difference in fitness scoring between the two strain backgrounds. Scatter plots show the fitness scores for each gene transposon knockout across the genome post CIP treatment (C) or without drug (D) for the drug susceptible WT strain alongside GPN. The noted genes are those that cause significant fitness changes in GPN compared to WT (where fitness differences ≥10%), with the exclusion of genes characterized as essential and genes associated with prophages or plasmids. Gene names are followed by accession numbers which are preceded by “NDEONHPJ_” per the ATCC 17978 strain annotations. Genes highlighted in teal are transposon disruptions that yield higher fitness scores in GPN compared to WT, with dark grey being those causing lower fitness in GPN (vs WT). CIP treatment used concentrations yielding 20-30% growth inhibition in each parent strain compared to the untreated grown in parallel. WT: wildtype. GPN: *gyrAparCadeN*.

**Supplemental Figure 19:** Fitness of transposon insertion mutants in GPS vs WT. Heatmaps of the normalized fitness of each transposon gene knockout in GPS and WT grown in the presence of CIP (A) or in the absence of drug pressure (B) alongside the difference in fitness scoring between the two strain backgrounds. Scatter plots show the fitness scores for each gene transposon knockout across the genome post CIP treatment (C) or without drug (D) for the drug susceptible WT strain alongside GPS. The noted genes are those that cause significant fitness changes in GPS compared to WT (where fitness differences ≥10%), with the exclusion of genes characterized as essential and genes associated with prophages or plasmids. Gene names are followed by accession numbers which are preceded by “NDEONHPJ_” per the ATCC 17978 strain annotations. Genes highlighted in teal are transposon disruptions that yield higher fitness scores in GPS compared to WT, with dark grey being those causing lower fitness in GPS (vs WT). CIP treatment used concentrations yielding 20-30% growth inhibition in each parent strain compared to the untreated grown in parallel. WT: wildtype. GPS: *gyrAparCadeS*.

**Supplemental Figure 20:** Nonadditive effects of overproducing two efflux pumps. At least three biological replicates of each strain were tested, all grown in LB with CIP concentrations yielding comparable growth inhibition of the parent strains, 0.12µg/mL or 60µg/mL (A) or 0.14µg/mL (B). After growing to early post-exponential, all cultures were diluted to A_600_~0.003 in LB broth and culture density monitored every hour for five hours (A) or every 15min overnight in a 96-well plate, though the first 8 hours are shown (B). Displayed is the mean of the replicates ± SEM. To note, WGS results showed that the *adeL*(∆I335A336) ∆*adeIJK* strain, which has no detectable alteration in drug sensitivity relative to the parental *adeL*(∆I335A336) strain, also has a mutation in *rpsC* (E170K).

**Supplemental Figure 21:** The observed MICs of CIP for the indicated null mutation strains are displayed as relative to their parent strains’ MIC (given a value of 1). WT: wildtype. GPL: *gyrAparCadeL*. GPN: *gyrAparCadeN*. GP: *gyrAparC*. ND: not determined.

**Supplemental Table 1:** Bacterial strains, plasmids, and primers used in this study.

**Supplemental Data Set 1:** TRANSIT Resampling data with comparisons made between FQR vs. WT strains.

**Supplemental Data Set 2:** TnSeq data with fitness comparisons made between FQR vs. WT strains.

**Supplemental Data Set 3:** RNAseq data with comparisons made for transcriptomes between the FQR vs. GP parent strains during CIP exposure.

**Supplemental Data Set 4:** RNAseq data with comparisons made for transcriptomes between the FQR vs. GP parent strains in the absence of drug treatment.
