## Supplementary Figures for "Identification of Determinants that Allow Maintenance of High-Level Fluoroquinolone Resistance in *Acinetobacter baumannii*"

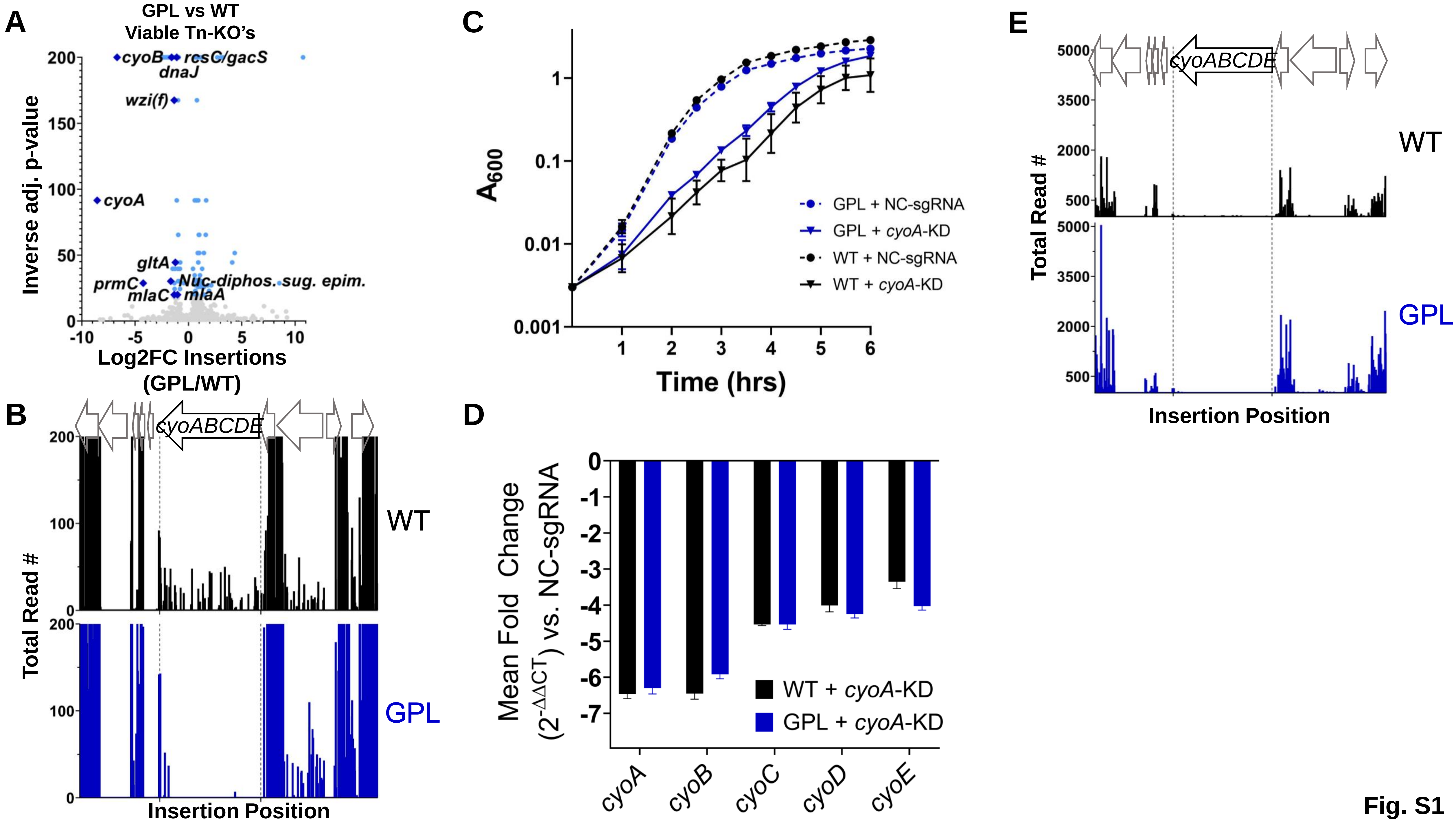
A
GPL vs WT
Viable Tn-KO's
Inverse adj. p-value
cyoB
resC/gacS
dnaJ
wzi(f)
cyoA
gltA
prmC
Nuc-diphos. sug. epim.
mlaC
mlaA
Log2FC Insertions (GPL/WT)
B
cyoABCDE
WT
GPL
Total Read #
Insertion Position
C
A600
Time (hrs)
GPL + NC-sgRNA
GPL + cyoA-KD
WT + NC-sgRNA
WT + cyoA-KD
D
Mean Fold Change (2-ΔΔCT) vs. NC-sgRNA
WT + cyoA-KD
GPL + cyoA-KD
cyoA
cyoB
cyoC
cyoD
cyoE
E
cyoABCDE
WT
GPL
Total Read #
Insertion Position
Fig. S1

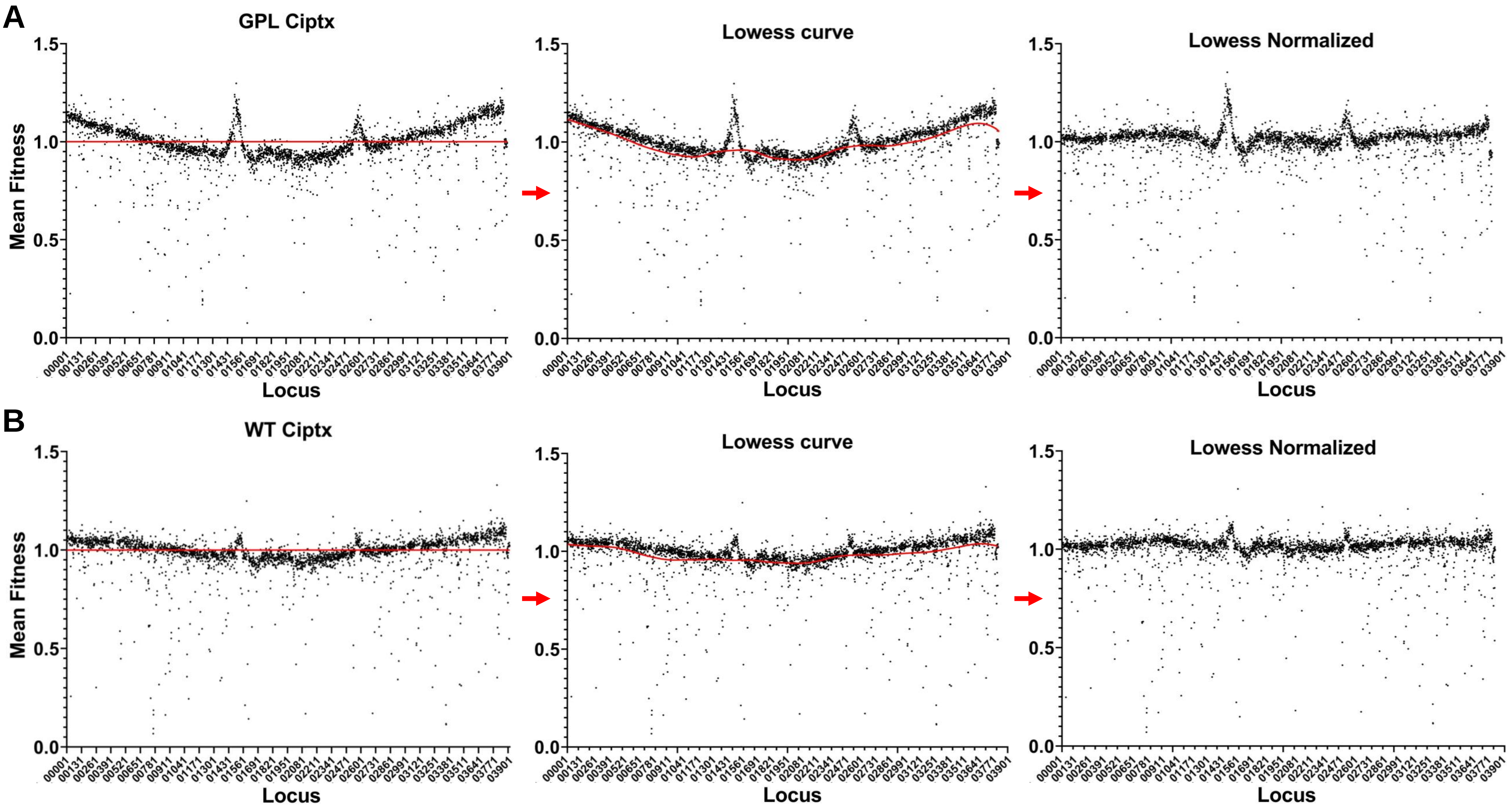
A
GPL Ciptx
Lowess curve
Lowess Normalized
B
WT Ciptx
Lowess curve
Lowess Normalized
Mean Fitness
Locus
1.5
1.0
0.5
0.0
00001
00131
00261
00391
00521
00651
00781
00911
01041
01171
01301
01431
01561
01691
01821
01951
02081
02211
02341
02471
02601
02731
02861
02991
03121
03251
03381
03511
03641
03771
03901

Fig. S2

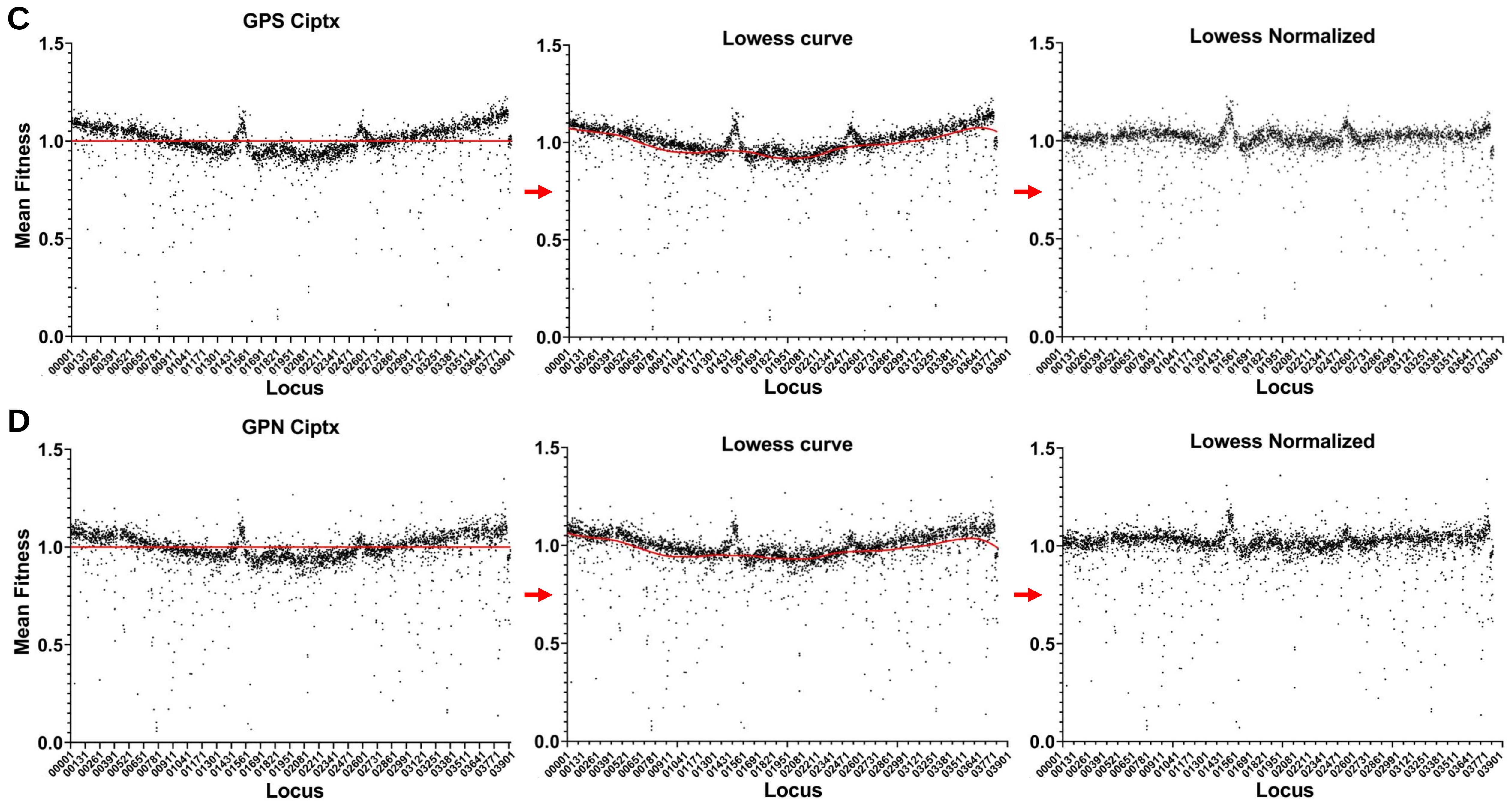
C
GPS Ciptx
Lowess curve
Lowess Normalized
Mean Fitness
Locus
D
GPN Ciptx
Lowess curve
Lowess Normalized
Mean Fitness
Locus

Fig. S2

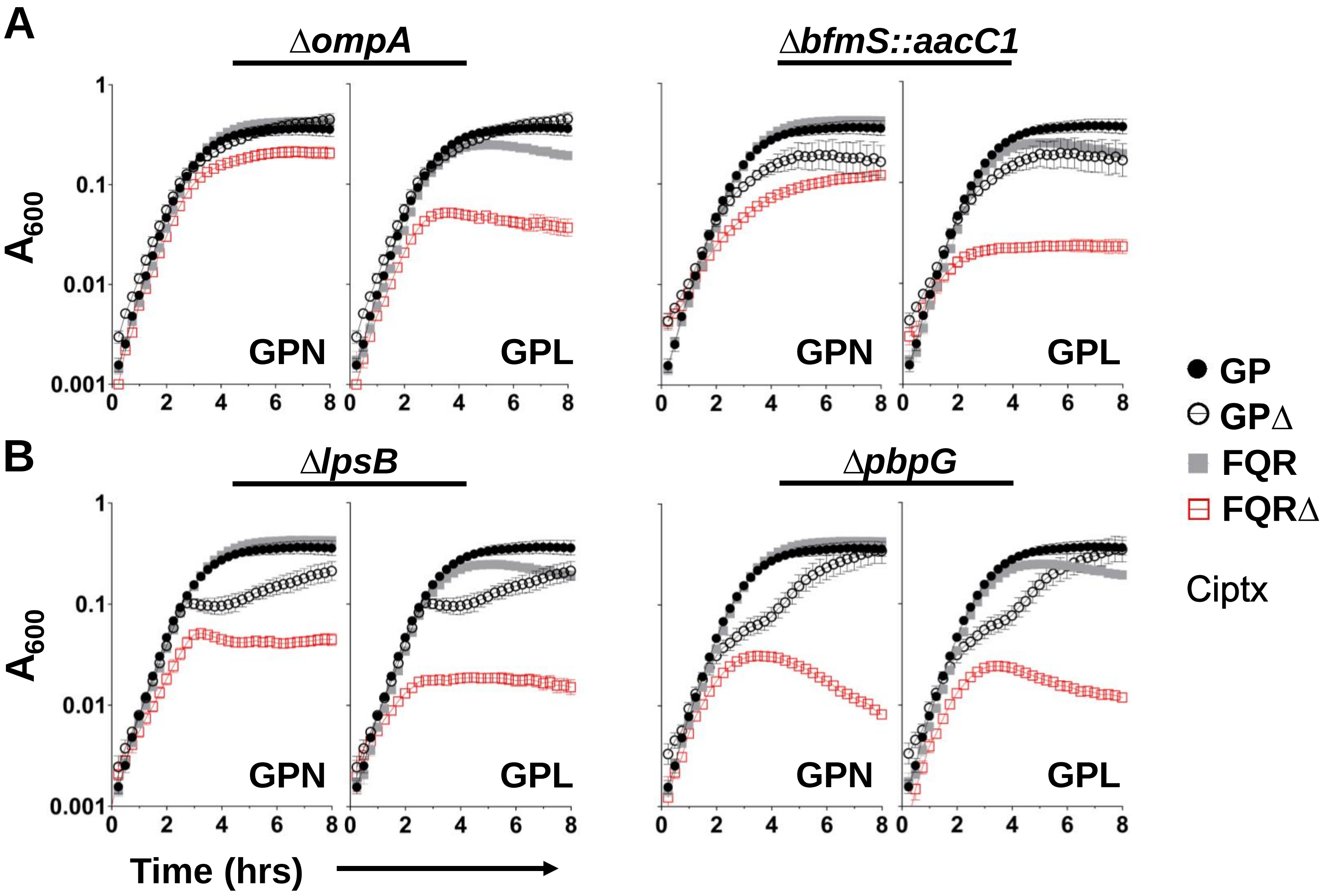
A
ΔompA
ΔbfmS::aacC1
A600
GPN
GPL
GPN
GPL
B
ΔlpsB
ΔpbpG
GPN
GPL
GPN
GPL
Time (hrs)
GP
GPΔ
FQR
FQRΔ
Ciptx

Fig. S3

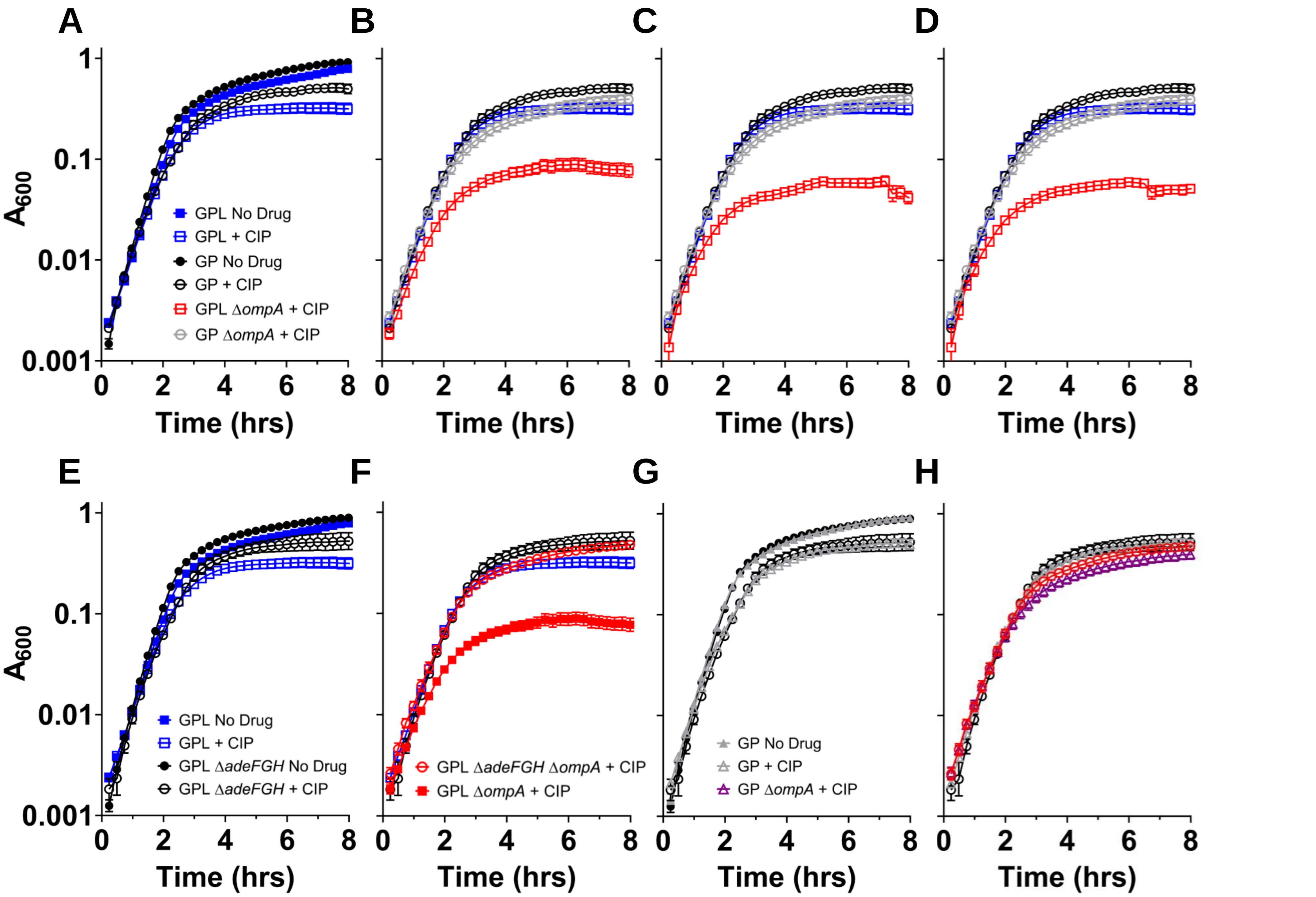
A
B
C
D
E
F
G
H
A600
Time (hrs)
GPL No Drug
GPL + CIP
GP No Drug
GP + CIP
GPL ΔompA + CIP
GP ΔompA + CIP
GPL ΔadeFGH No Drug
GPL ΔadeFGH + CIP
GPL ΔadeFGH ΔompA + CIP
GPL ΔompA + CIP
GP No Drug
GP + CIP
GP ΔompA + CIP

Fig. S4

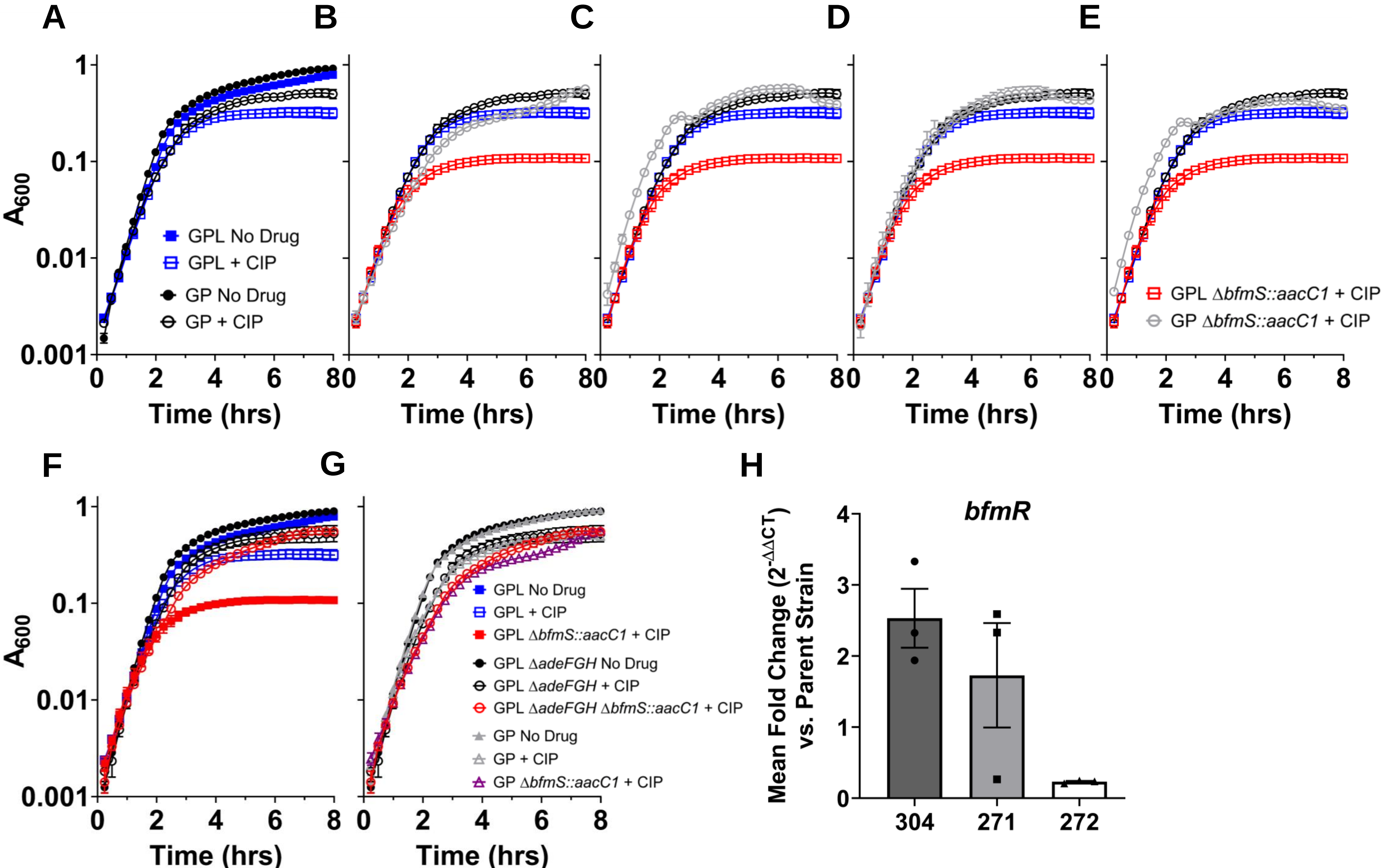
A
B
C
D
E
A600
GPL No Drug
GPL + CIP
GP No Drug
GP + CIP
GPL ΔbfmS::aacC1 + CIP
GP ΔbfmS::aacC1 + CIP
Time (hrs)
F
G
H
GPL No Drug
GPL + CIP
GPL ΔbfmS::aacC1 + CIP
GPL ΔadeFGH No Drug
GPL ΔadeFGH + CIP
GPL ΔadeFGH ΔbfmS::aacC1 + CIP
GP No Drug
GP + CIP
GP ΔbfmS::aacC1 + CIP
bfmR
Mean Fold Change (2^-ΔΔCT) vs. Parent Strain
304
271
272

Fig. S5

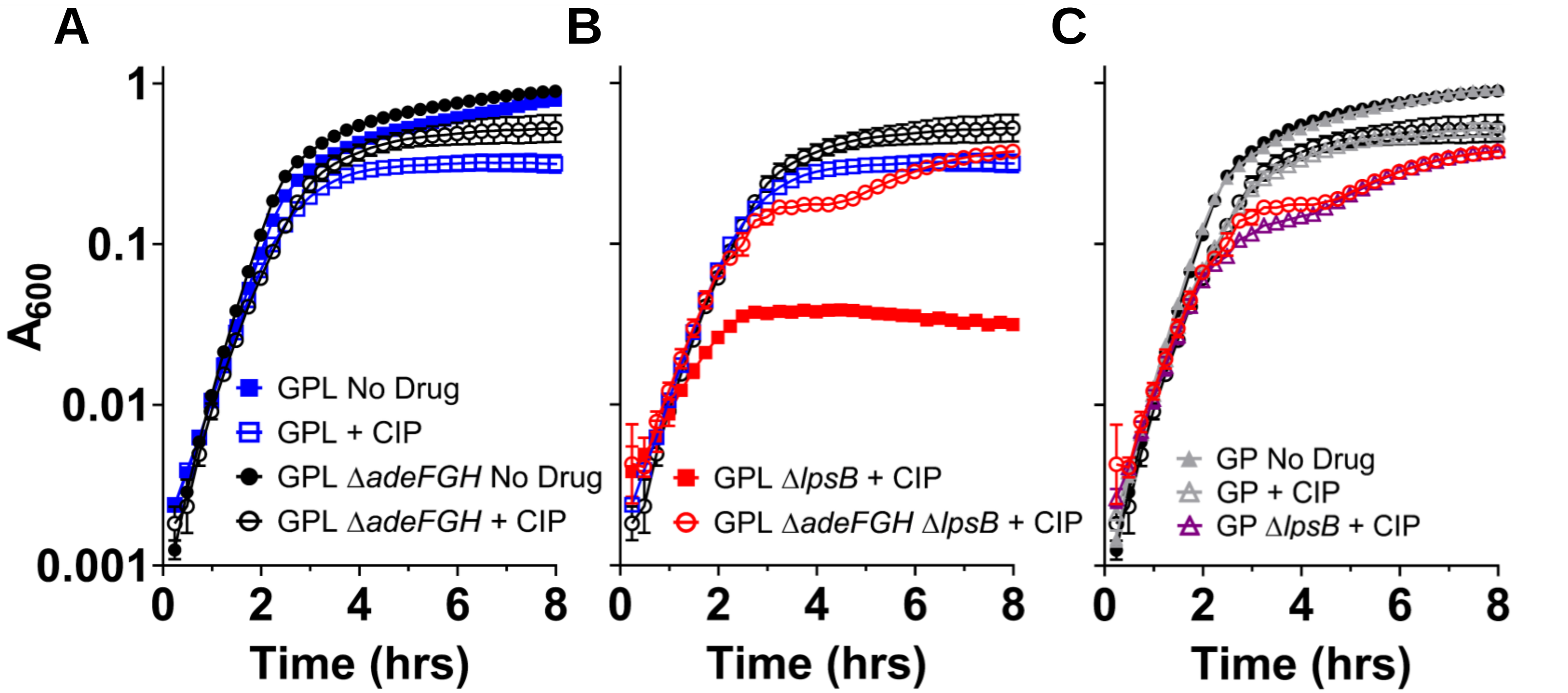
A
B
C
A600
1
0.1
0.01
0.001
0
2
4
6
8
Time (hrs)
GPL No Drug
GPL + CIP
GPL ΔadeFGH No Drug
GPL ΔadeFGH + CIP
GPL ΔlpsB + CIP
GPL ΔadeFGH ΔlpsB + CIP
GP No Drug
GP + CIP
GP ΔlpsB + CIP

Fig. S6

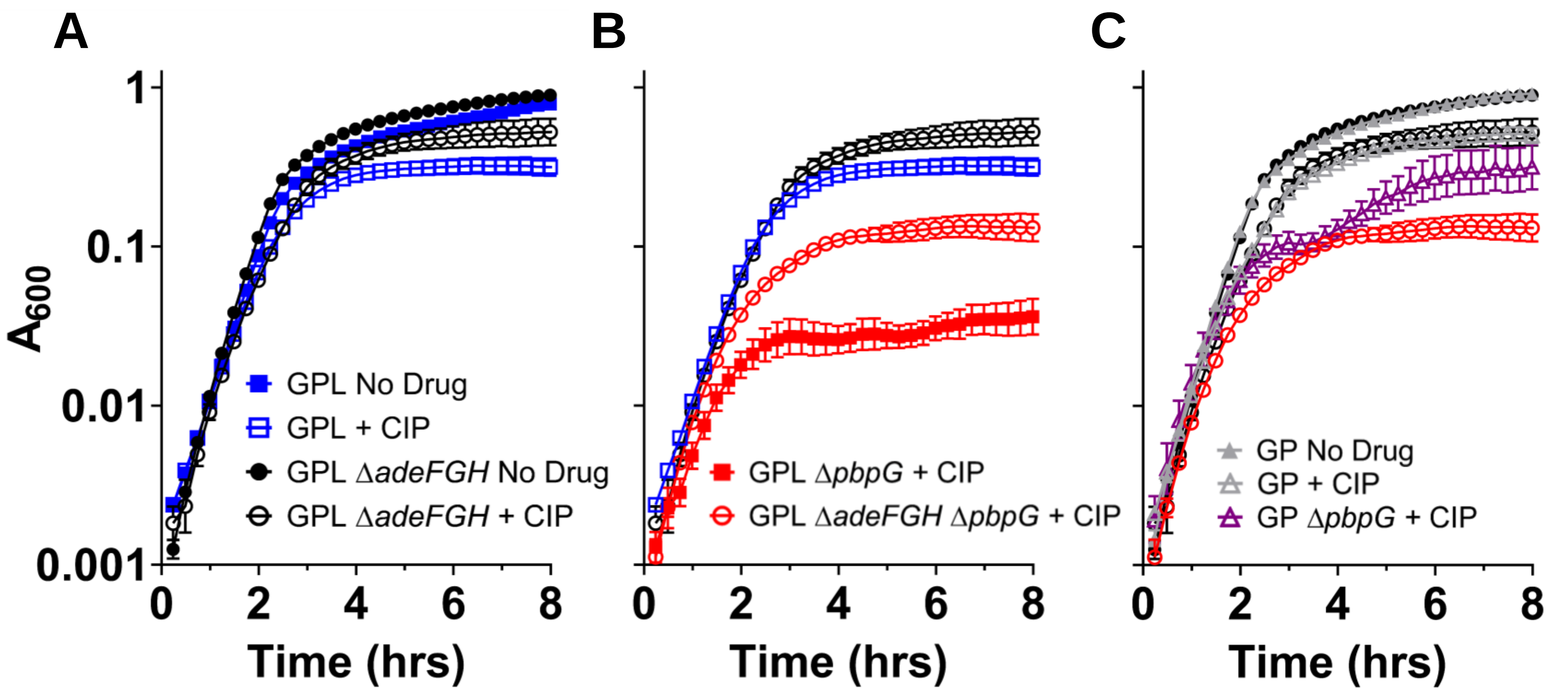
A
B
C
1
0.1
0.01
0.001
A600
GPL No Drug
GPL + CIP
GPL ΔadeFGH No Drug
GPL ΔadeFGH + CIP
GPL ΔpbpG + CIP
GPL ΔadeFGH ΔpbpG + CIP
GP No Drug
GP + CIP
GP ΔpbpG + CIP
0
2
4
6
8
Time (hrs)

**Fig. S7**

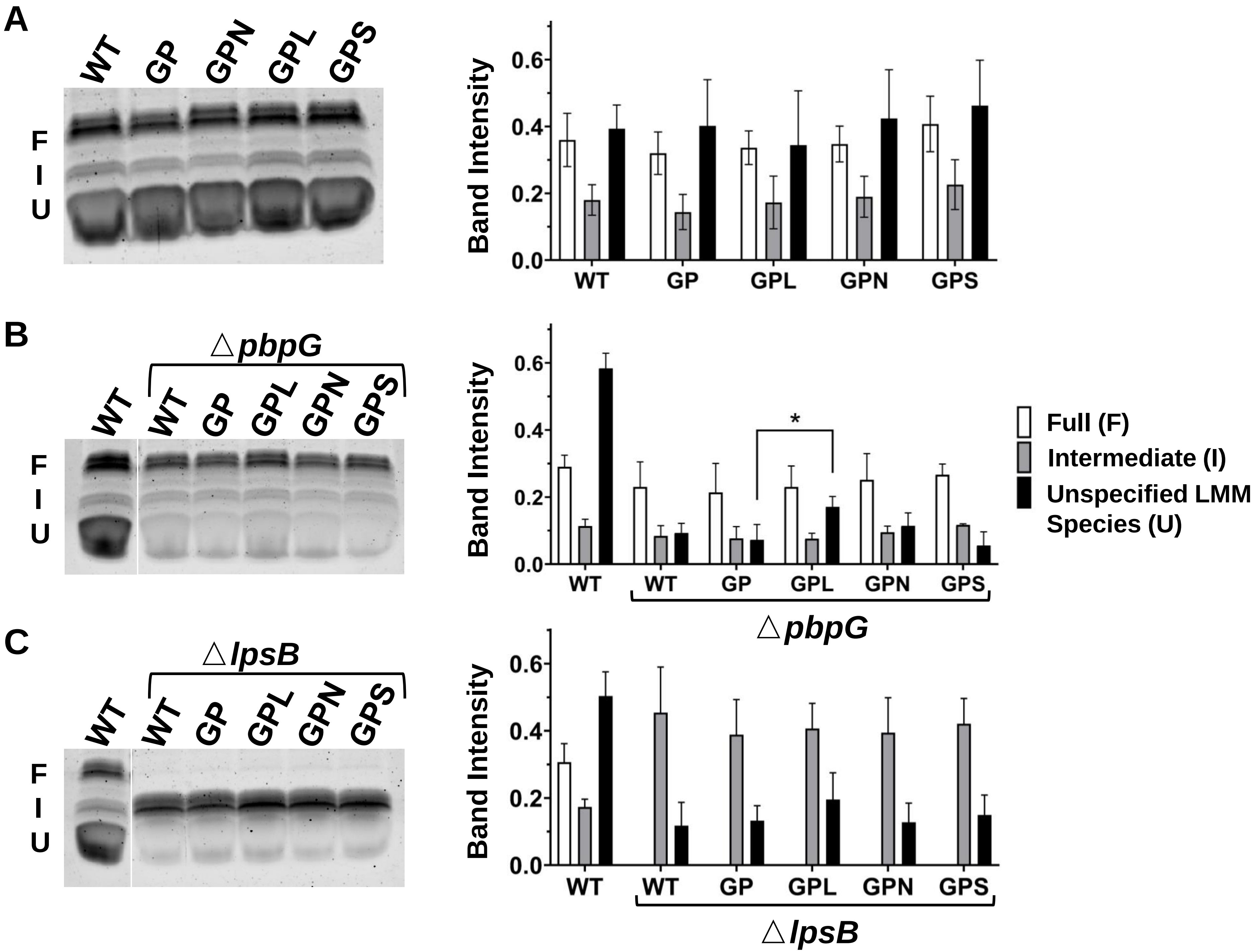
A
WT GP GPN GPL GPS
F
I
U
Band Intensity
0.0 0.2 0.4 0.6
WT GP GPL GPN GPS
B
△pbpG
WT WT GP GPL GPN GPS
F
I
U
Band Intensity
0.0 0.2 0.4 0.6
*
WT WT GP GPL GPN GPS
△pbpG
Full (F)
Intermediate (I)
Unspecified LMM Species (U)
C
△lpsB
WT WT GP GPL GPN GPS
F
I
U
Band Intensity
0.0 0.2 0.4 0.6
WT WT GP GPL GPN GPS
△lpsB

Fig. S8

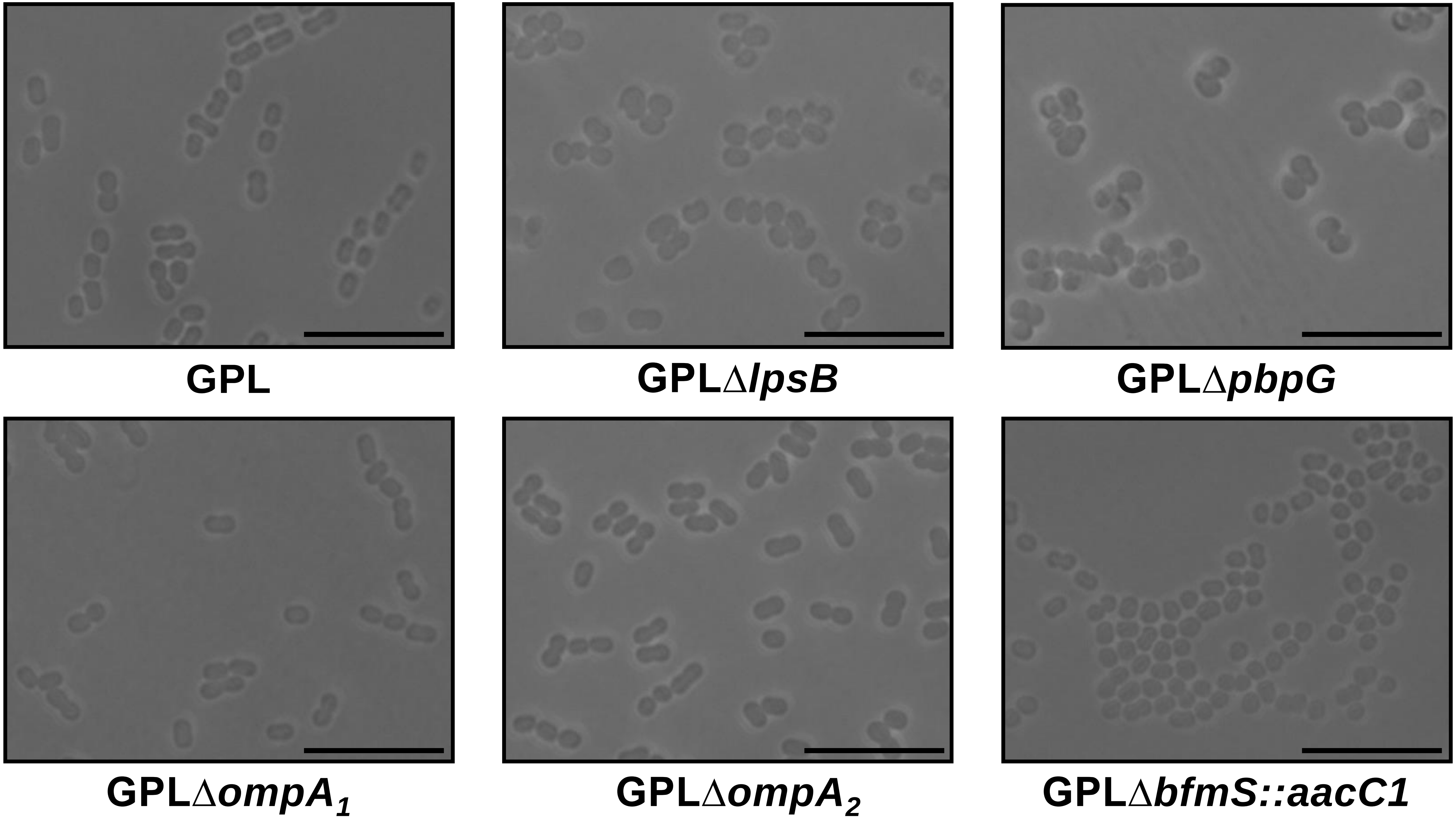
GPL
GPLΔ*lpsB*
GPLΔ*pbpG*
GPLΔ*ompA*$_1$
GPLΔ*ompA*$_2$
GPLΔ*bfmS::aacC1*

**Fig. S9**

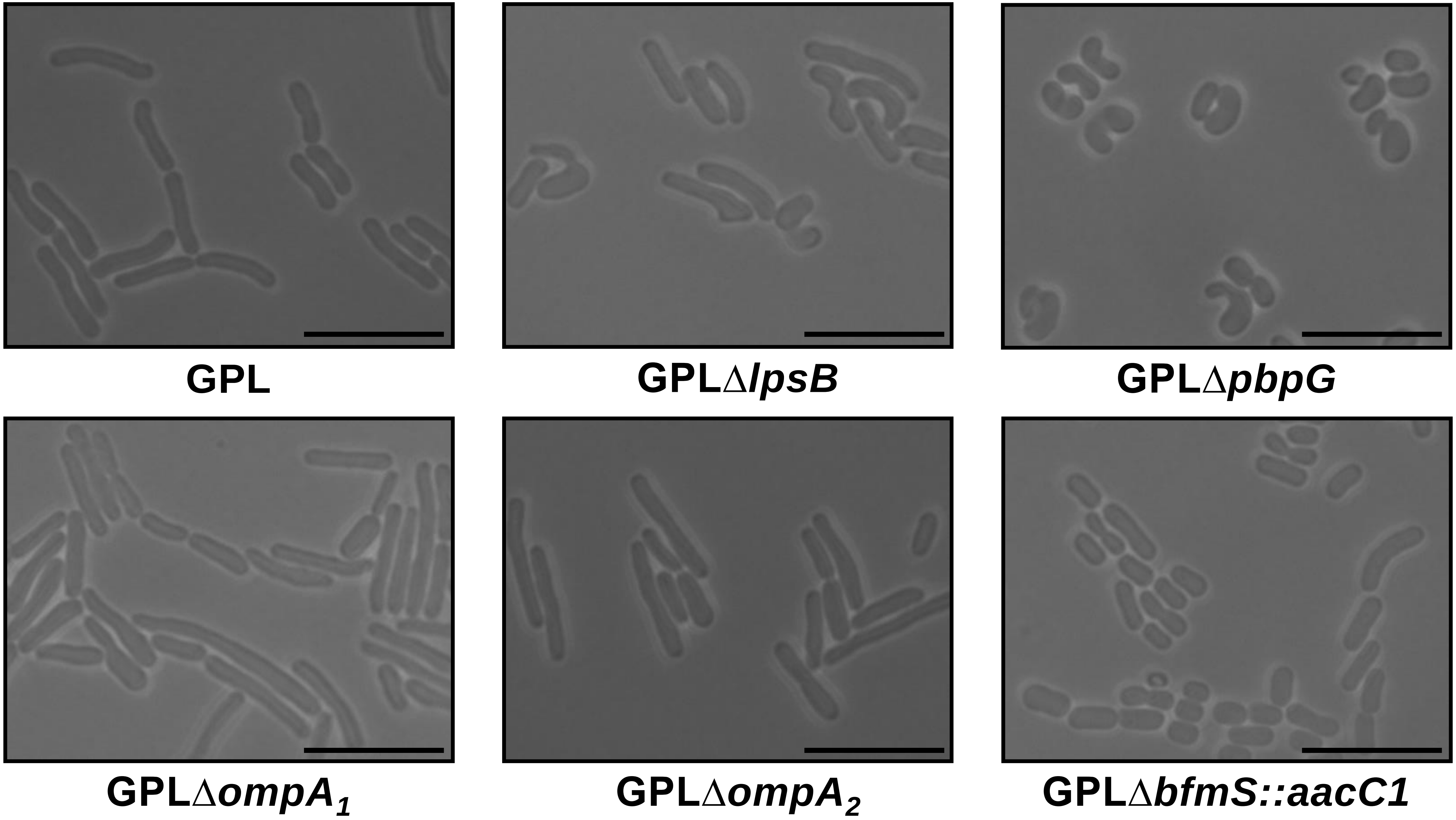
GPL
GPLΔ*lpsB*
GPLΔ*pbpG*
GPLΔ$ompA_1$
GPLΔ$ompA_2$
GPLΔ*bfmS::aacC1*

**Fig. S10**

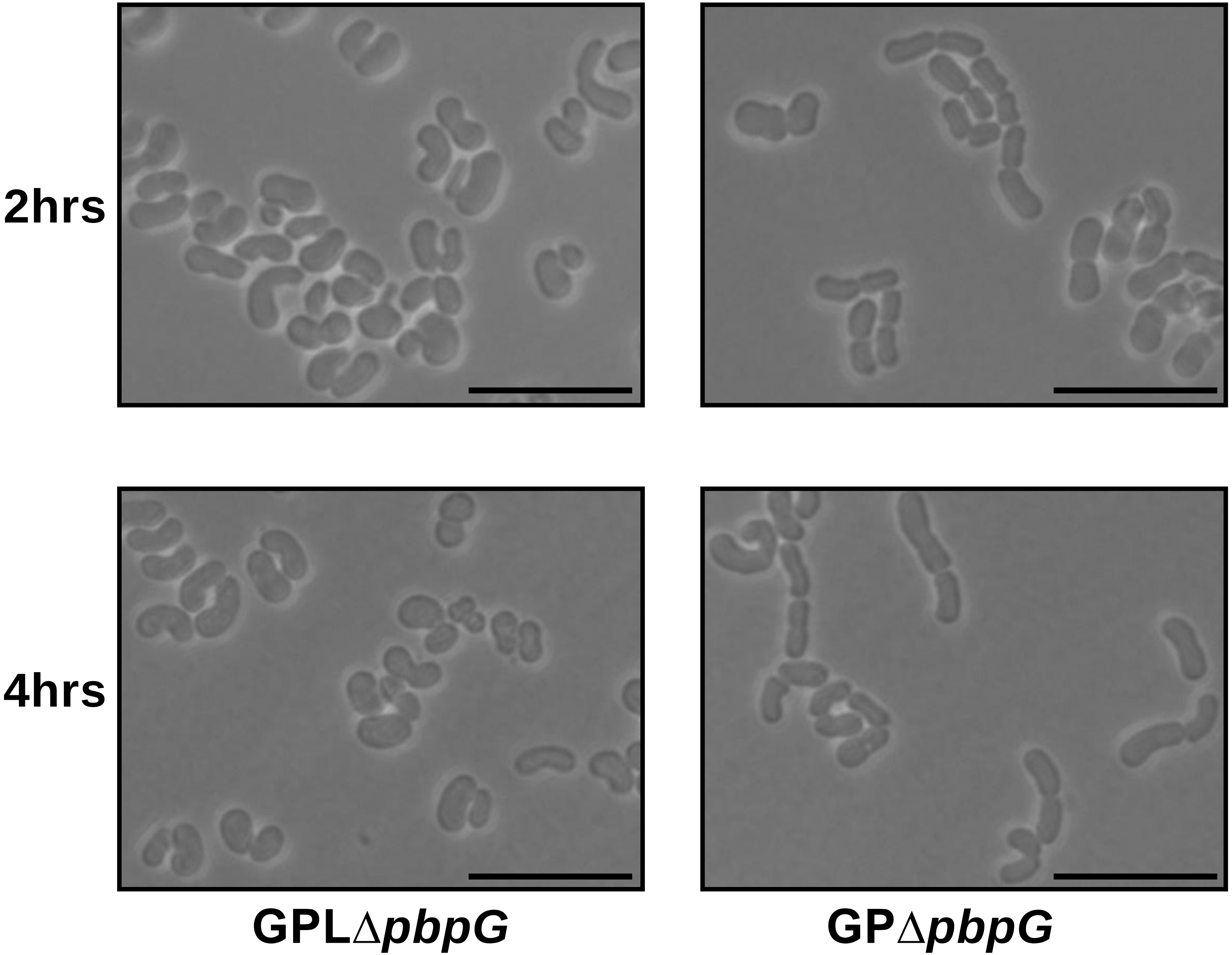


**Fig. S11**

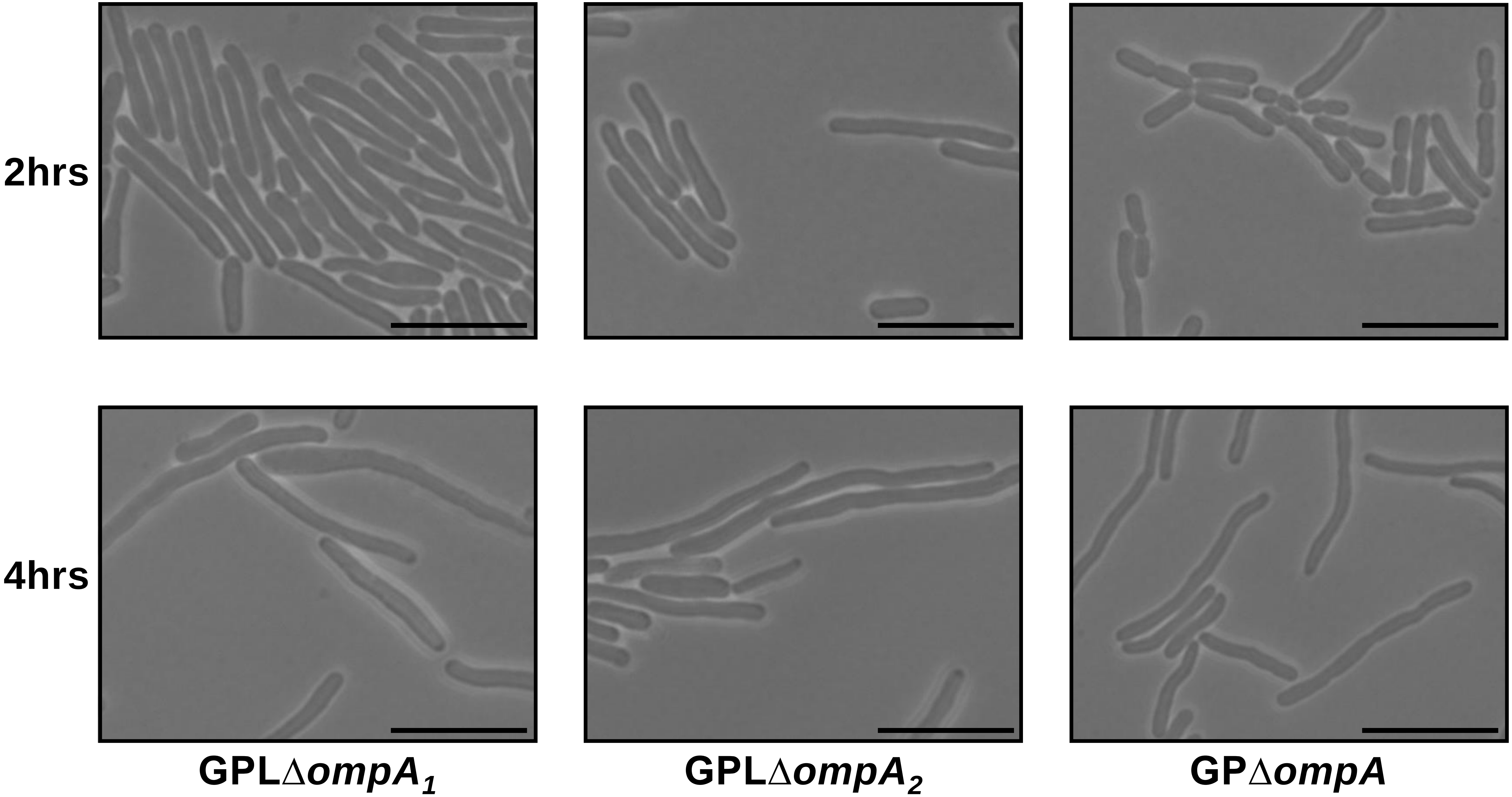


**Fig. S12**

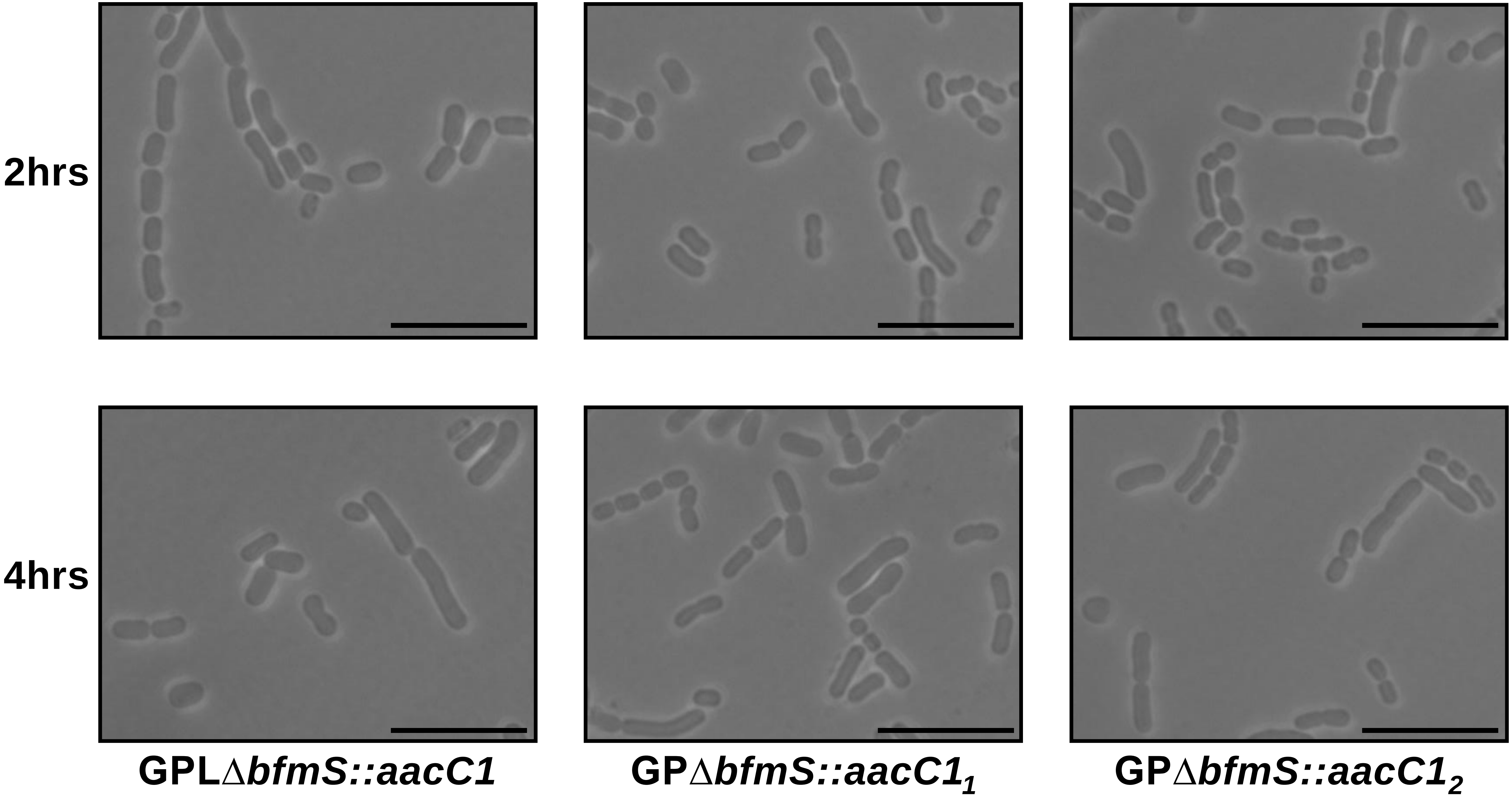


Fig. S13

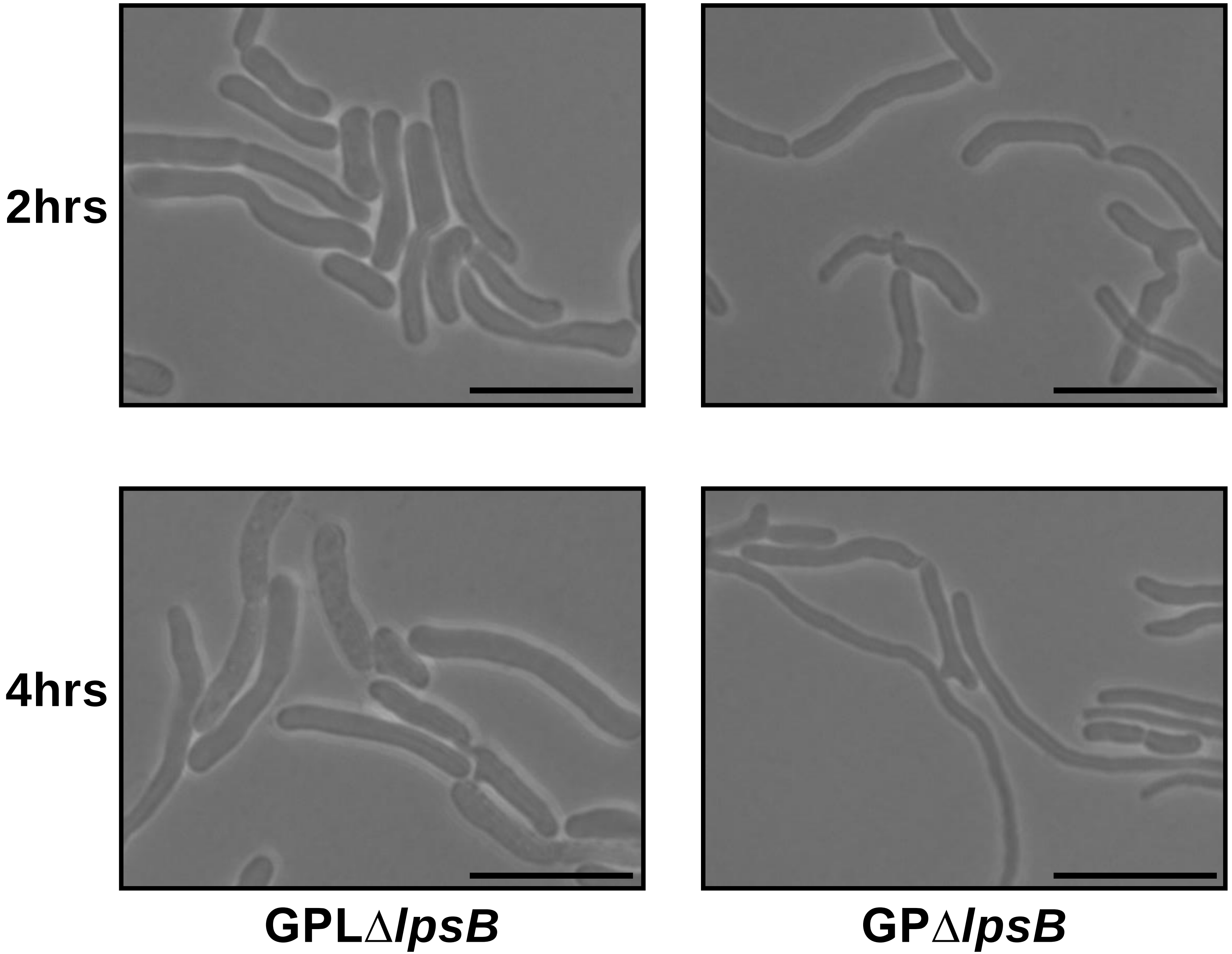


**Fig. S14**

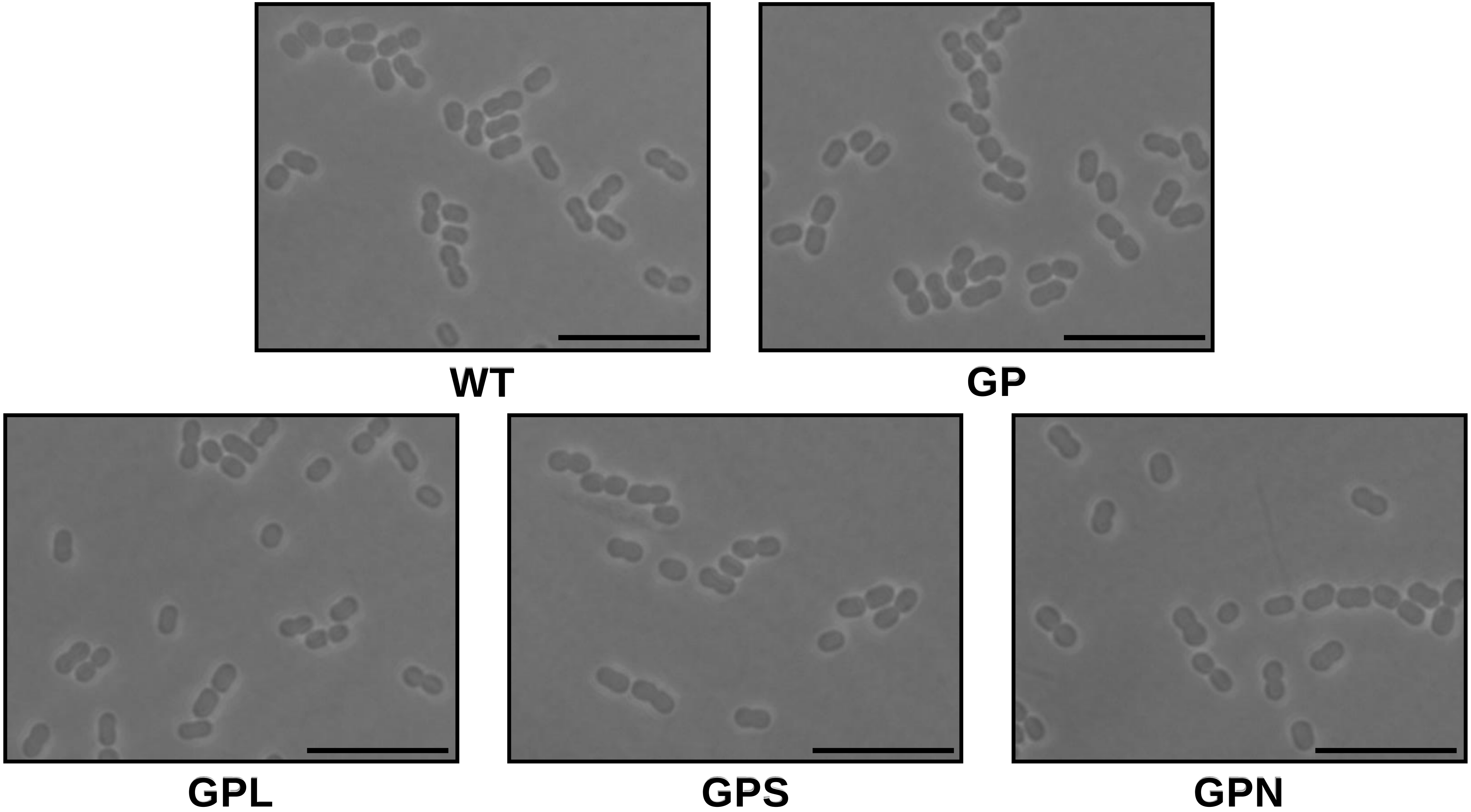


Fig. S15

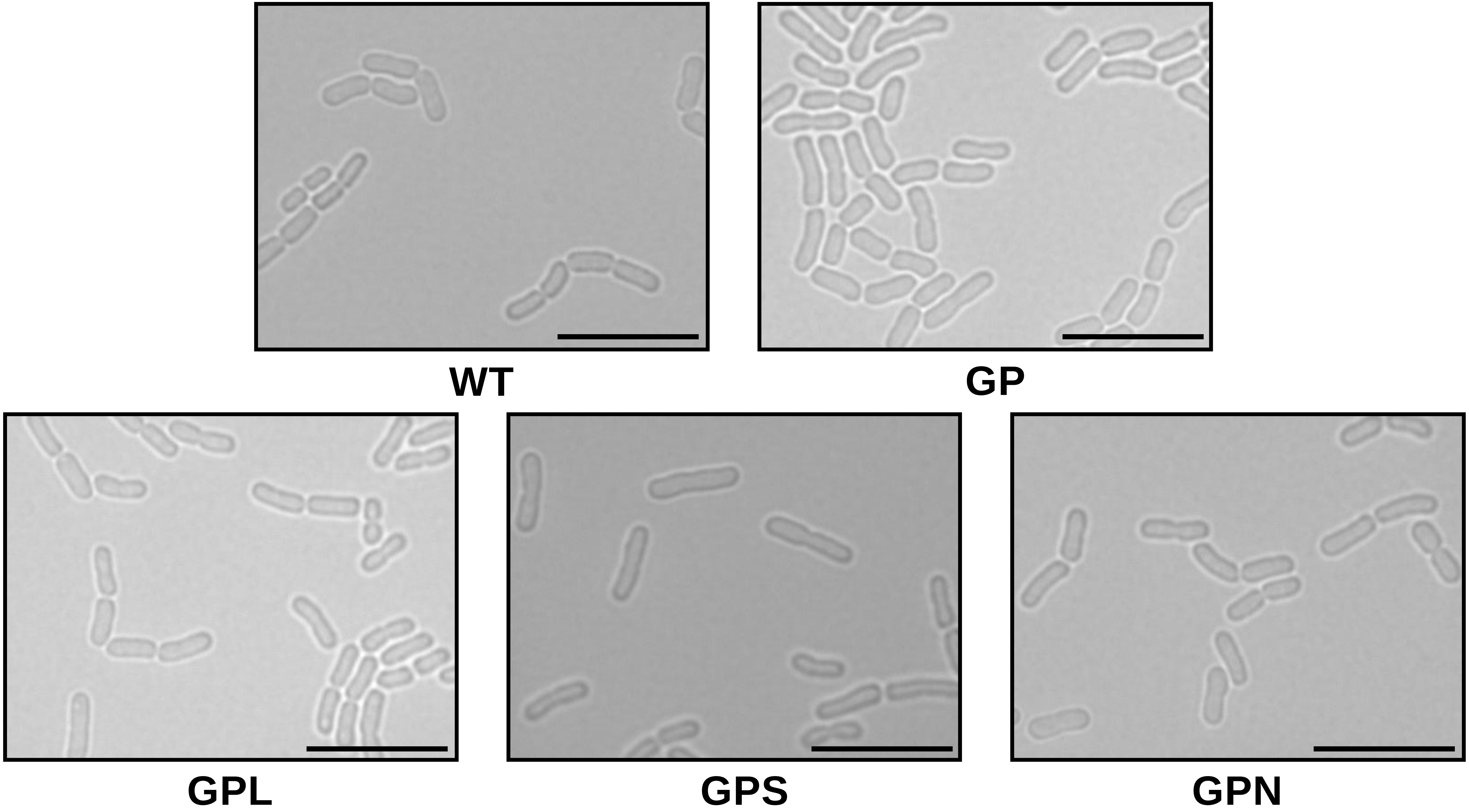


Fig. S16

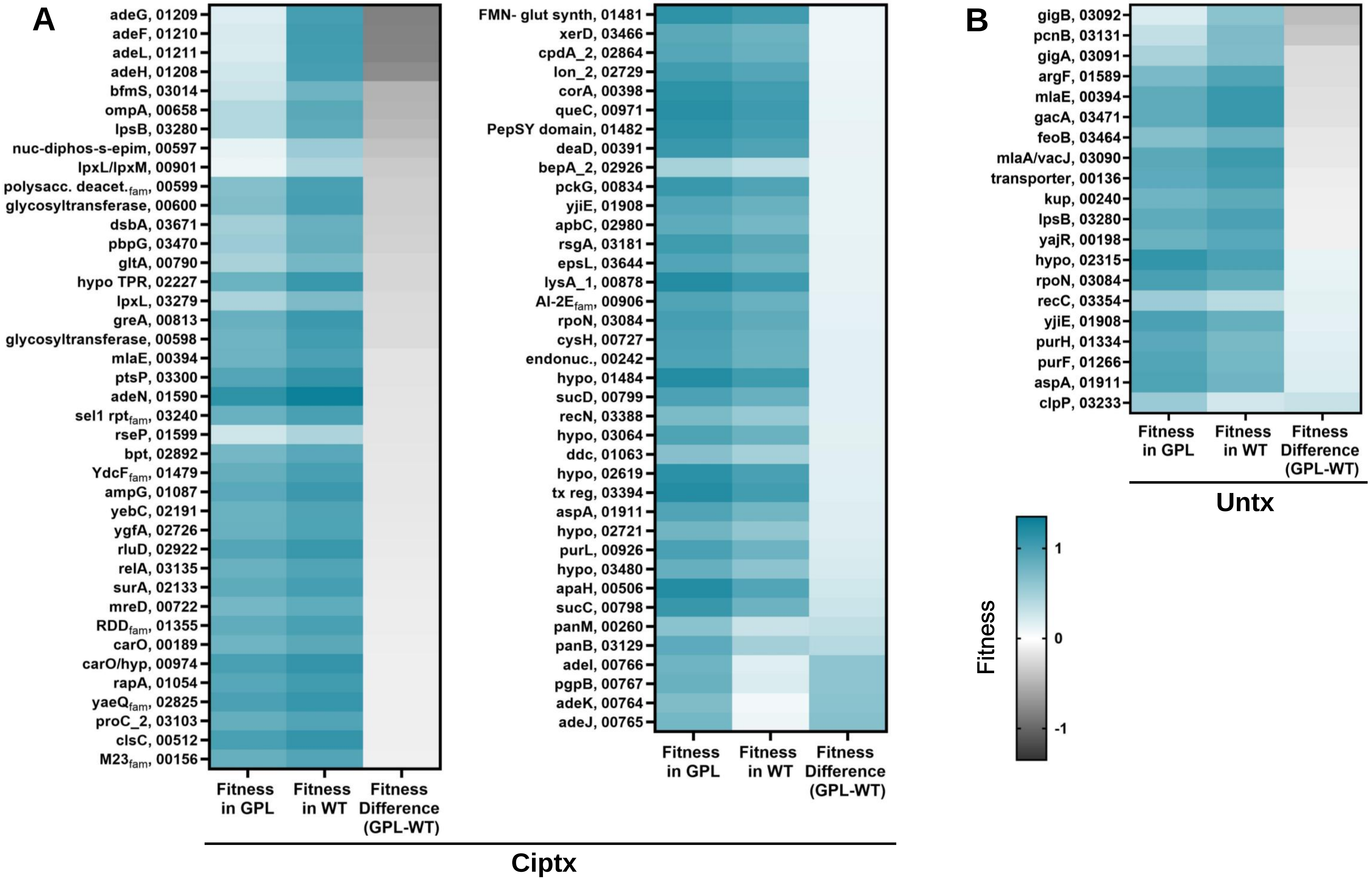
A
adeG, 01209
adeF, 01210
adeL, 01211
adeH, 01208
bfmS, 03014
ompA, 00658
lpsB, 03280
nuc-diphos-s-epim, 00597
lpxL/lpxM, 00901
polysacc. deacet.fam, 00599
glycosyltransferase, 00600
dsbA, 03671
pbpG, 03470
gltA, 00790
hypo TPR, 02227
lpxL, 03279
greA, 00813
glycosyltransferase, 00598
mlaE, 00394
ptsP, 03300
adeN, 01590
sel1 rptfam, 03240
rseP, 01599
bpt, 02892
YdcFfam, 01479
ampG, 01087
yebC, 02191
ygfA, 02726
rluD, 02922
relA, 03135
surA, 02133
mreD, 00722
RDDfam, 01355
carO, 00189
carO/hyp, 00974
rapA, 01054
yaeQfam, 02825
proC_2, 03103
clsC, 00512
M23fam, 00156
FMN- glut synth, 01481
xerD, 03466
cpdA_2, 02864
lon_2, 02729
corA, 00398
queC, 00971
PepSY domain, 01482
deaD, 00391
bepA_2, 02926
pckG, 00834
yjiE, 01908
apbC, 02980
rsgA, 03181
epsL, 03644
lysA_1, 00878
AI-2Efam, 00906
rpoN, 03084
cysH, 00727
endonuc., 00242
hypo, 01484
sucD, 00799
recN, 03388
hypo, 03064
ddc, 01063
hypo, 02619
tx reg, 03394
aspA, 01911
hypo, 02721
purL, 00926
hypo, 03480
apaH, 00506
sucC, 00798
panM, 00260
panB, 03129
adeI, 00766
pgpB, 00767
adeK, 00764
adeJ, 00765
Fitness in GPL
Fitness in WT
Fitness Difference (GPL-WT)
Ciptx
B
gigB, 03092
pcnB, 03131
gigA, 03091
argF, 01589
mlaE, 00394
gacA, 03471
feoB, 03464
mlaA/vacJ, 03090
transporter, 00136
kup, 00240
lpsB, 03280
yajR, 00198
hypo, 02315
rpoN, 03084
recC, 03354
yjiE, 01908
purH, 01334
purF, 01266
aspA, 01911
clpP, 03233
Fitness in GPL
Fitness in WT
Fitness Difference (GPL-WT)
Untx
Fitness
1
0
-1

Fig. S17

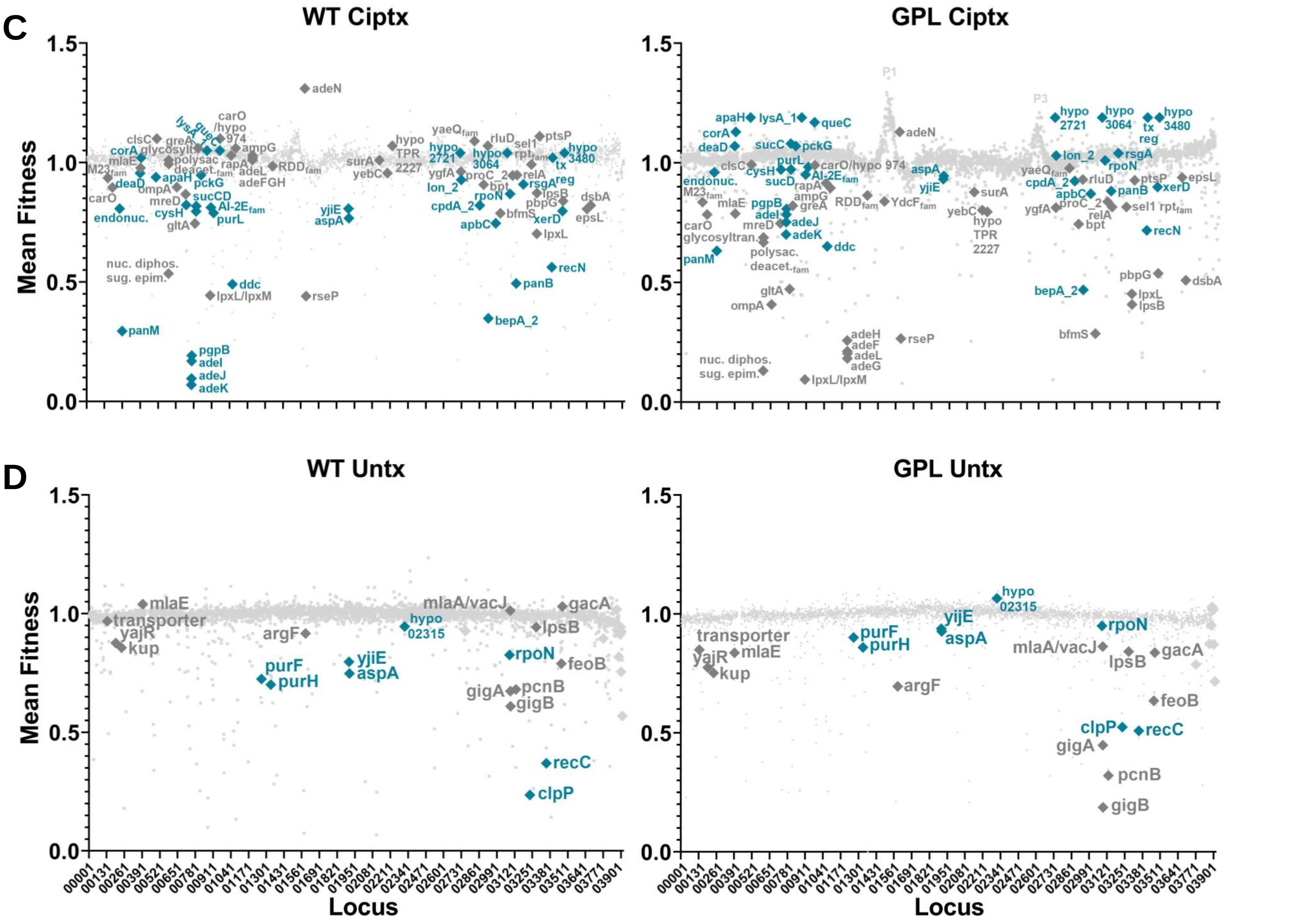
C
WT Ciptx
GPL Ciptx
D
WT Untx
GPL Untx
Mean Fitness
Locus
1.5
1.0
0.5
0.0
00001
00131
00261
00391
00521
00651
00781
00911
01041
01171
01301
01431
01561
01691
01821
01951
02081
02211
02341
02471
02601
02731
02861
02991
03121
03251
03381
03511
03641
03771
03901
adeN
carO
/hypo
974
clsC
greA
glycosylt.
corA
mlaE
M23fam
deaD
apaH
pckG
polysac
deacet.fam
rapA
ampG
adeL
adeFGH
RDDfam
carO
ompA
mreD
sucCD
cysH
AI-2Efam
purL
endonuc.
gltA
nuc. diphos.
sug. epim.
ddc
lpxL/lpxM
rseP
panM
pgpB
adeI
adeJ
adeK
hypo
TPR
2227
surA
yebC
yaeQfam
hypo
2721
hypo
3064
rluD
sel1
rptfam
ptsP
hypo
3480
tx
reg
ygfA
proC_2
relA
bpt
lon_2
rsgA
rpoN
lpsB
pbpG
dsbA
cpdA_2
bfmS
xerD
epsL
apbC
lpxL
yjiE
aspA
recN
panB
bepA_2
P1
P3
lysA_1
queC
sucC
sucD
purL
carO/hypo 974
glycosyltran.
mreD
polysac.
deacet.fam
RDDfam
YdcFfam
adeH
adeF
adeG
rpoN
recN
epsL
lpxB
mlaE
transporter
yajR
kup
argF
purF
purH
hypo
02315
mlaA/vacJ
gacA
feoB
pcnB
gigA
gigB
recC
clpP

Fig. S17

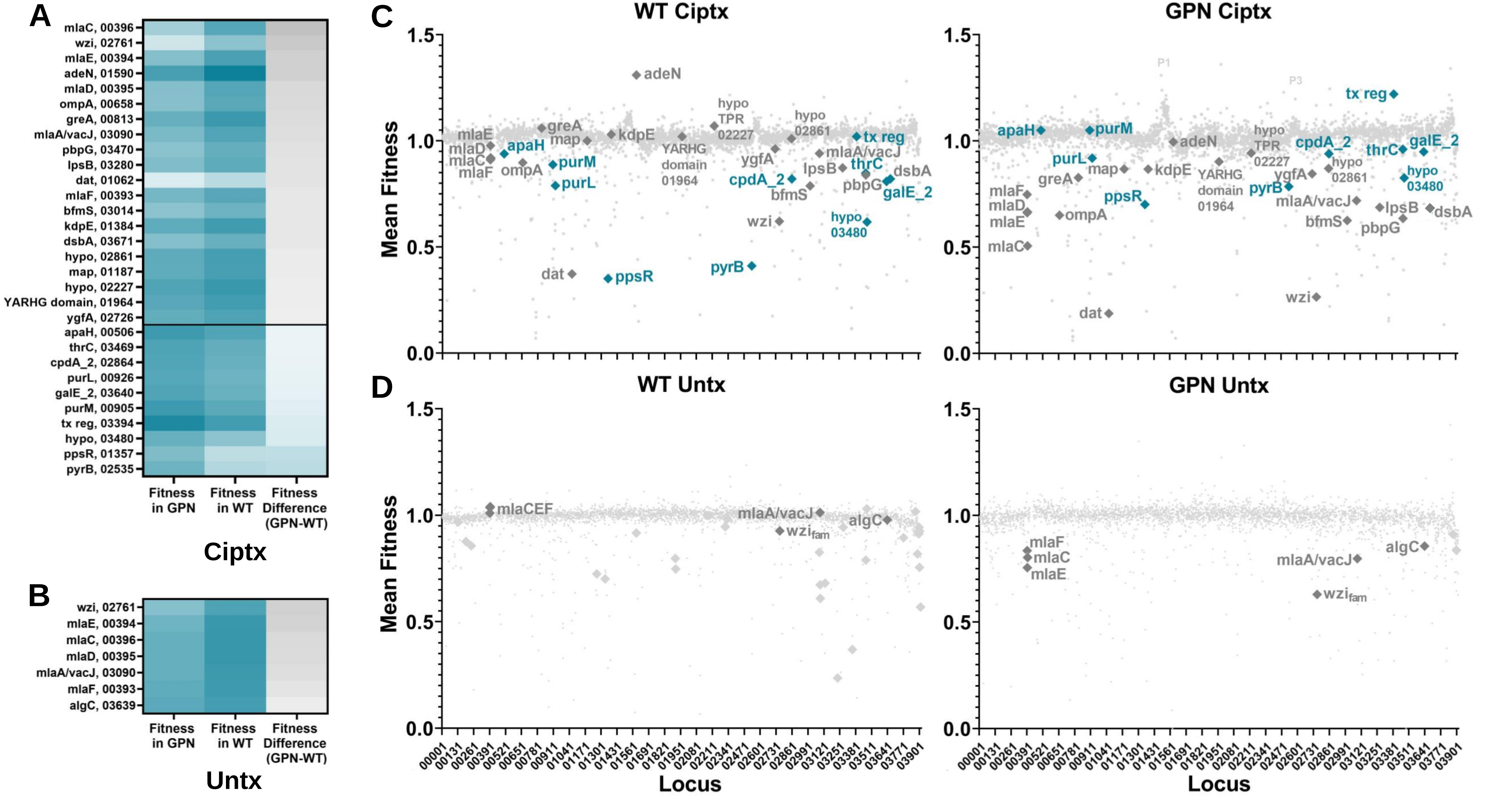
A
mlaC, 00396
wzi, 02761
mlaE, 00394
adeN, 01590
mlaD, 00395
ompA, 00658
greA, 00813
mlaA/vacJ, 03090
pbpG, 03470
lpsB, 03280
dat, 01062
mlaF, 00393
bfmS, 03014
kdpE, 01384
dsbA, 03671
hypo, 02861
map, 01187
hypo, 02227
YARHG domain, 01964
ygfA, 02726
apaH, 00506
thrC, 03469
cpdA_2, 02864
purL, 00926
galE_2, 03640
purM, 00905
tx reg, 03394
hypo, 03480
ppsR, 01357
pyrB, 02535
Fitness in GPN
Fitness in WT
Fitness Difference (GPN-WT)
Ciptx
B
wzi, 02761
mlaE, 00394
mlaC, 00396
mlaD, 00395
mlaA/vacJ, 03090
mlaF, 00393
algC, 03639
Fitness in GPN
Fitness in WT
Fitness Difference (GPN-WT)
Untx
C
WT Ciptx
GPN Ciptx
D
WT Untx
GPN Untx
Mean Fitness
Locus

Fig. S18

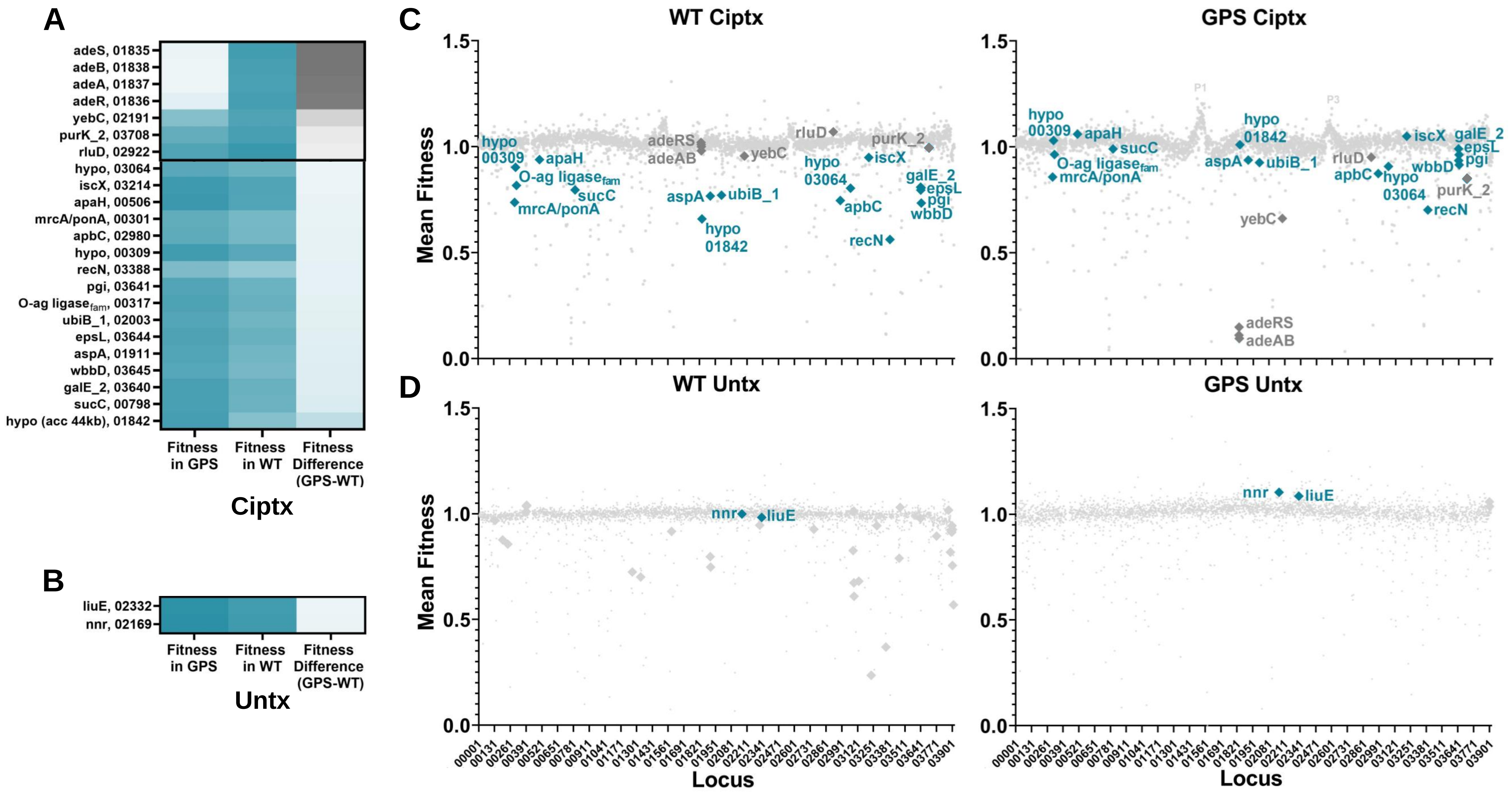
A
adeS, 01835
adeB, 01838
adeA, 01837
adeR, 01836
yebC, 02191
purK_2, 03708
rluD, 02922
hypo, 03064
iscX, 03214
apaH, 00506
mrcA/ponA, 00301
apbC, 02980
hypo, 00309
recN, 03388
pgi, 03641
O-ag ligase$_{fam}$, 00317
ubiB_1, 02003
epsL, 03644
aspA, 01911
wbbD, 03645
galE_2, 03640
sucC, 00798
hypo (acc 44kb), 01842
Fitness in GPS
Fitness in WT
Fitness Difference (GPS-WT)
Ciptx
B
liuE, 02332
nnr, 02169
Fitness in GPS
Fitness in WT
Fitness Difference (GPS-WT)
Untx
C
WT Ciptx
GPS Ciptx
D
WT Untx
GPS Untx
Mean Fitness
Locus
P1
P3
adeRS
adeAB
yebC
rluD
purK_2
hypo 00309
apaH
O-ag ligase$_{fam}$
sucC
mrcA/ponA
aspA
ubiB_1
hypo 01842
hypo 03064
iscX
apbC
recN
galE_2
epsL
pgi
wbbD
nnr
liuE

Fig. S19

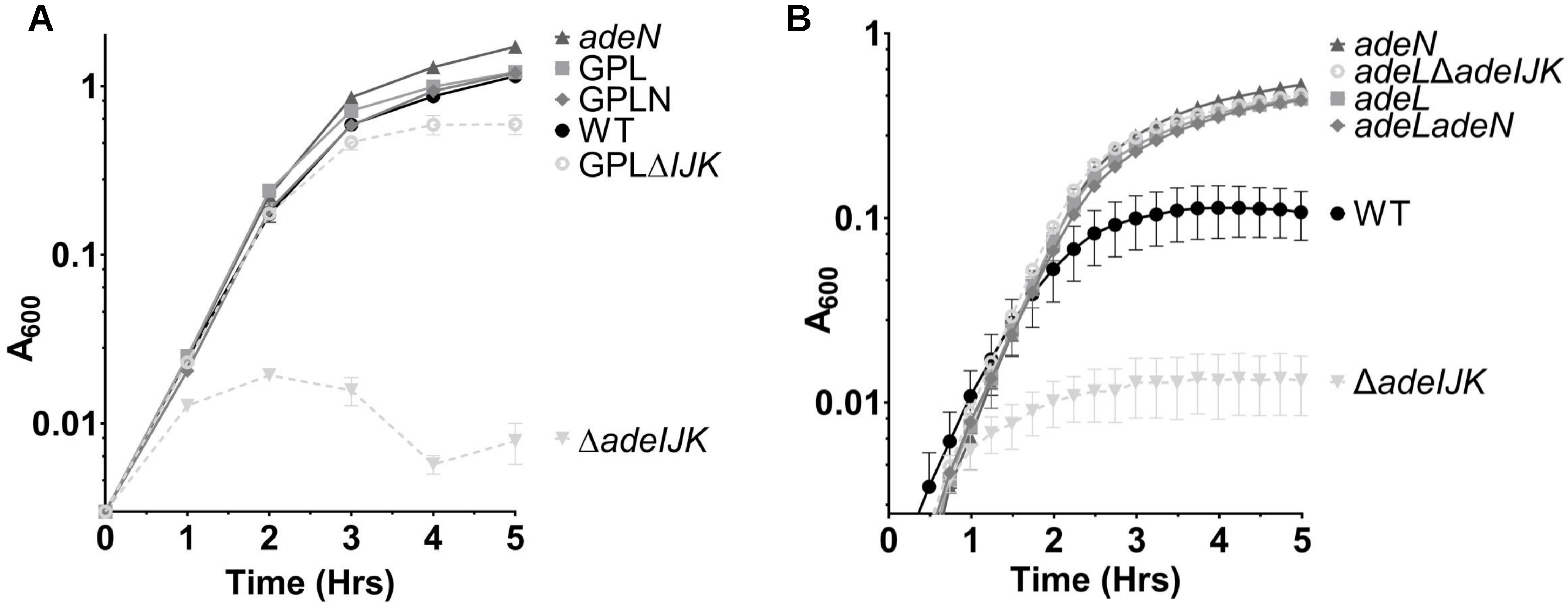
A
adeN
GPL
GPLN
WT
GPLΔIJK
ΔadeIJK
A600
Time (Hrs)
B
adeN
adeLΔadeIJK
adeL
adeLadeN
WT
ΔadeIJK
A600
Time (Hrs)

Fig. S20

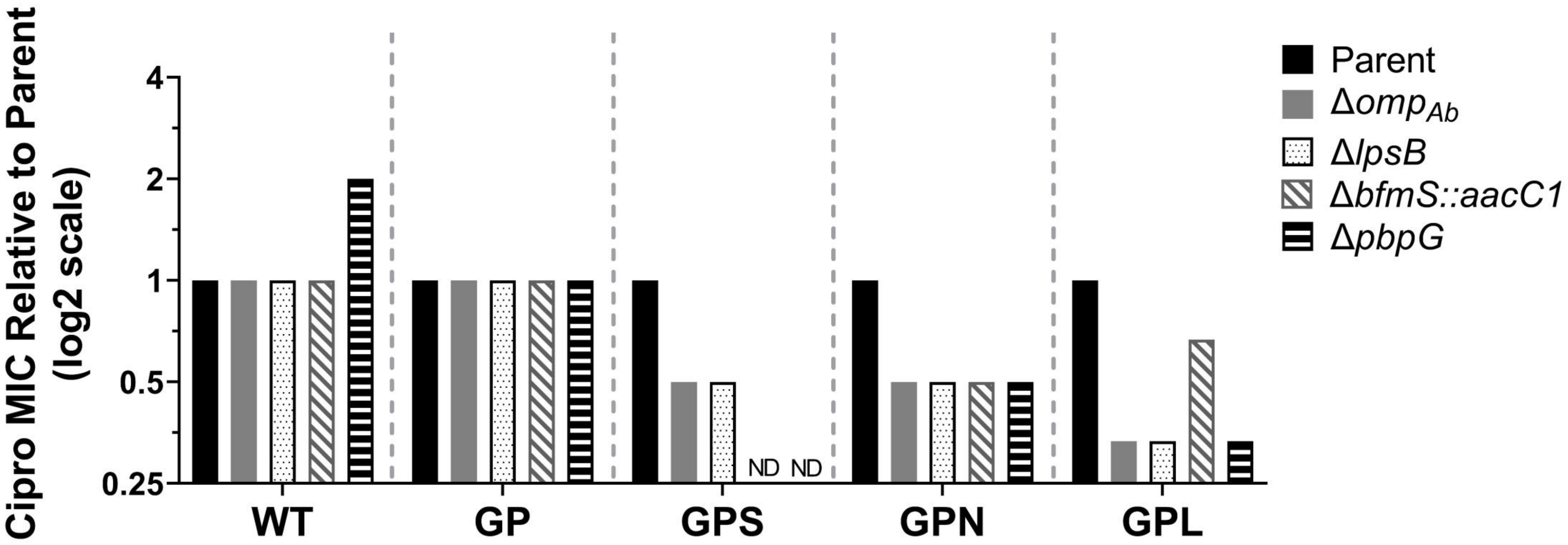
Cipro MIC Relative to Parent (log2 scale)
4
2
1
0.5
0.25
ND
ND
WT
GP
GPS
GPN
GPL
Parent
Δomp$_{Ab}$
ΔlpsB
ΔbfmS::aacC1
ΔpbpG

**Fig. S21**
